# A lymph node-to-tumour PDL1^+^macrophage circuit antagonizes dendritic cell immunotherapy

**DOI:** 10.1101/2023.03.14.532534

**Authors:** Jenny Sprooten, Isaure Vanmeerbeek, Angeliki Datsi, Jannes Govaerts, Daniel M Borràs, Stefan Naulaerts, Raquel S. Laureano, Anna Calvet, Marc Kuballa, Michael C. Sabel, Marion Rapp, Christiane Knobbe-Thomsen, Peng Liu, Liwei Zhao, Oliver Kepp, Guido Kroemer, Louis Boon, Sabine Tejpar, Jannie Borst, Susan Schlenner, Steven De Vleeschouwer, Rüdiger V. Sorg, Abhishek D Garg

**Affiliations:** Laboratory of Cell Stress & Immunity, Department of Cellular & Molecular Medicine, KU Leuven, Belgium; Institute for Transplantation Diagnostics and Cell Therapeutics, Medical Faculty, Heinrich Heine University Hospital, Düsseldorf, Germany; Department of Neurosurgery, Medical Faculty, Heinrich Heine University Hospital, Düsseldorf, Germany; Metabolomics and Cell Biology Platforms, Gustave Roussy Cancer Center, Université Paris Saclay, Villejuif, France; Centre de Recherche des Cordeliers, Equipe labellisée par la Ligue contre le cancer, Université de Paris, Sorbonne Université, Inserm U1138, Institut Universitaire de France, Paris, France; Institut du Cancer Paris CARPEM, Department of Biology, Hôpital Européen Georges Pompidou, AP-HP, Paris, France; JJP Biologics, Warsaw, Poland; Laboratory for Molecular Digestive Oncology, Department of Oncology, KU Leuven, Belgium; Department of Immunology and Oncode Institute, Leiden University Medical Center, Leiden, Netherlands; Department of Microbiology, Immunology and Transplantation, KU Leuven-University of Leuven, Leuven, Belgium; Department of Neurosurgery, University Hospitals Leuven, Leuven, Belgium; Laboratory of Experimental Neurosurgery and Neuroanatomy, Department of Neurosciences, KU Leuven, Belgium; Leuven Brain Institute (LBI), Leuven, Belgium

**Keywords:** macrophages, damage-associated molecular patterns (DAMPs), tumour-associated antigens (TAAs), apoptosis, necroptosis, mature-regulatory DCs (mregDC), immune checkpoint blockers, tumour-infiltrating lymphocytes, glioblastoma

## Abstract

Immune-checkpoint blockers (ICB) provide limited benefit against T cell-depleted tumours, calling for therapeutic innovation. Here, we aimed at designing a new type of dendritic cell (DC) vaccine by unbiased computational integration of multi-omics data from cancer patients. In a first attempt, a DC vaccine designed to present tumor antigens from cancer cells succumbing to immunogenic cancer cell death (ICD) and to elicit high type I interferon (IFN) responses failed to induce the regression of mouse tumors lacking T cell infiltrates. In lymph nodes (LNs), instead of activating CD4^+^ and CD8^+^T cells, DCs stimulated immunosuppressive PD-L1^+^LN-associated macrophages (LAMs) via a type I IFN response. Moreover, DC vaccines of this type stimulated pre-existing, T cell-suppressive, PD-L1^+^tumour-associated macrophages (TAMs). This created a T cell-suppressive circuit of PD-L1^+^macrophages, spanning across LNs and tumours. Accordingly, DC vaccines synergised with PD-L1 blockade to deplete PD-L1^+^macrophages, suppress myeloid inflammation affecting the tumor bed and draining lymph nodes, and de-inhibit effector/stem-like memory T cells, eventually causing tumour regression. The synergistic interaction between the DC vaccine and PD-L1 blockade was lost when DCs were manipulated to lose *Ifnar1*or *Ccr7* or when macrophages were depleted. Interestingly, clinical DC vaccines also potentiated lymphocyte-suppressive PD-L1^+^TAMs in patients bearing T cell-depleted tumours. Altogether, our results reveal the existence of a novel PD-L1^+^LAM/TAM-driven immunosuppressive pathway that can be elicited by DC vaccines, yet can be subverted for improving the outcome of immunotherapy.

## INTRODUCTION

Cancer immunotherapy via immune-checkpoint blockers (ICBs) has improved the clinical outlook for many patients (Sharma and Allison, 2020). However, not all patients or cancer-types respond to ICBs (Sharma and Allison, 2020). The success of ICBs relies on rejuvenating effector/stem-like memory CD8^+^T cells, that are typically pre-enriched in T cell-infiltrated tumours (Blank et al., 2019; Naulaerts et al., 2021; Thommen and Schumacher, 2018; Vanmeerbeek et al., 2021). Therefore, terminally exhausted, or severely dysfunctional CD8^+^T cells (with effector/memory-function disabilities and increased propensity to cell death), together with poor antigenicity within T cell-depleted tumours, pose a strong barrier to ICBs (Blank et al., 2019; Naulaerts et al., 2021; Thommen and Schumacher, 2018). This immuno-resistant barrier is further reinforced by tumour-associated macrophages (TAMs) with anti-inflammatory activity (M2-like) (Beyranvand Nejad et al., 2020; Goswami et al., 2022; Nakamura and Smyth, 2020).

But, such ICB-resistant tumours (Naulaerts et al., 2021) do present a niche for T cell-enhancing cellular immunotherapies like dendritic cell (DC)-based vaccines (Goswami et al., 2022; Naulaerts et al., 2021; Perez and De Palma, 2019). Such vaccines utilize patient-derived autologous DCs (most commonly, monocytes-derived DCs, or moDCs), that are pulsed with cancer antigens and stimulated with maturation stimuli *ex vivo*, before re-injection into the patient (Anguille et al., 2014; van Beek et al., 2020). The potential of DC vaccines in activating antigen-specific immunogenicity is well-established since DCs are highly proficient antigen-presenting cells (Wculek et al., 2020). This is due to their unique ability to cross-present antigens to both CD4^+^/CD8^+^T cells in the lymph nodes (LNs). Thereby, DC vaccines mobilize antigen-specific CD4^+^/CD8^+^T cells (Kvedaraite and Ginhoux, 2022). These features have inspired the rationale of anti-cancer DC vaccines (Garg et al., 2017).

Unfortunately, these pro-immunogenic capacities of DC vaccines have not consistently translated into clinically-meaningful tumour regression (Anguille et al., 2014; Garg et al., 2017). However recently, the lack of efficacy of ICBs against T cell-depleted tumours, has spurred the interest in multi-modal immunotherapy regimen integrating ‘next-generation’ DC vaccines (Fucikova et al., 2022; Garg et al., 2017; van Beek et al., 2020). For such a regimen to succeed, it is important to understand the exact immunological disparities affecting DC vaccines, to inform combinations with other immunotherapies. It is still unclear which of the following reasons are behind the dismal performance of DC vaccines: limited immunogenic potential, inefficacious maturation, gaps between mouse vs. human translation of vaccine designs, and/or as-yet-unknown immuno-resistance pathways. Addressing these challenges simultaneously via reverse translational (human-to-mice and back) research is urgently needed to properly position DC vaccines in specific immuno-oncology niches. Hence, the overarching aim of this study was to use a human (pan-)cancer omics-driven framework to inform the design of a novel next-generation DC vaccine with highly Immunogenic maturation-Trajectory (DCvax-IT). We wanted to tailor this DC vaccine against ICB non-responsive, T cell-depleted tumours; and then decipher the exact immunological mechanisms mediating the optimal vs. sub-optimal anti-tumour performance of DCvax-IT. Finally, we aimed to confirm our pre-clinical observations in human clinical settings as proof-of-concept for future designing of multi-modal immunotherapy trials integrating DC vaccines.

## RESULTS

### Dysregulated danger signalling, defective type I interferon (IFN)-production and DC depletion distinguishes the human T cell-depleted tumours

The designing of the DC vaccines against T cell-depleted tumours in this study aimed to use a reverse translational approach. For this, we first needed to extract the most dysregulated extra-cellular immune-pathways from human T cell-depleted tumours (apart from the lack of T cell-infiltration) that might be restraining tumour immunogenicity. This informed two crucial steps for our study: (I) the designing of DC vaccines integrating required immunogenic signalling; and (II) the rationale (unbiased) selection of a mouse tumour model that mimics the dominant immune-disparities of human T cell-depleted tumours.

To extract the immune-disparities, we analysed the well-established tumour immune-landscapes in The Cancer Genome Atlas (TCGA) datasets, to delineate the most T cell-depleted tumours (Thorsson et al., 2018). Previously, an immunogenomics analysis had identified six pan-cancer immune-landscape classes (C1-to-C6; for class-labels, see X-axis of the heatmap in **Fig.1A**) (Thorsson et al., 2018). We uniformly analysed them, across 3546 patients spanning 13 cancer-types, for major immune cell fractions or their ratios, and T cell-receptor (TCR) richness (a metric for TCR poly-clonality) (**Fig.1A**) (Naulaerts et al., 2021; Thorsson et al., 2018). As compared to C1-C3/C6, C4/C5-tumour immune-landscapes showed the most striking depletion of CD4^+^/CD8^+^T cell-fractions, TCR richness, and even DCs abundance (**Fig.1A**). Contrastingly, C4/C5-tumours had high ratios of macrophages-to-CD8^+^T cell fractions and M2 (anti-inflammatory) to M1 (pro-inflammatory) macrophage-polarizations (**Fig.1A**). Consequently, in an integrated multi-cancer dataset of 704 patients (spanning 5 cancer-types) whose tumours were transcriptome-profiled before ICBs-treatment (Eddy et al., 2020), overall survival of patients with C4/C5-tumours was significantly shorter than those with immunogenic C2/C3/C6-tumours (**Fig.1B**). This highlighted the suitability of TCGA C4/C5-tumours as transcriptomic representatives of ICB-resistant non-immunogenic tumours, to which we will refer to as T cell-depleted tumours.

**Figure 1.**
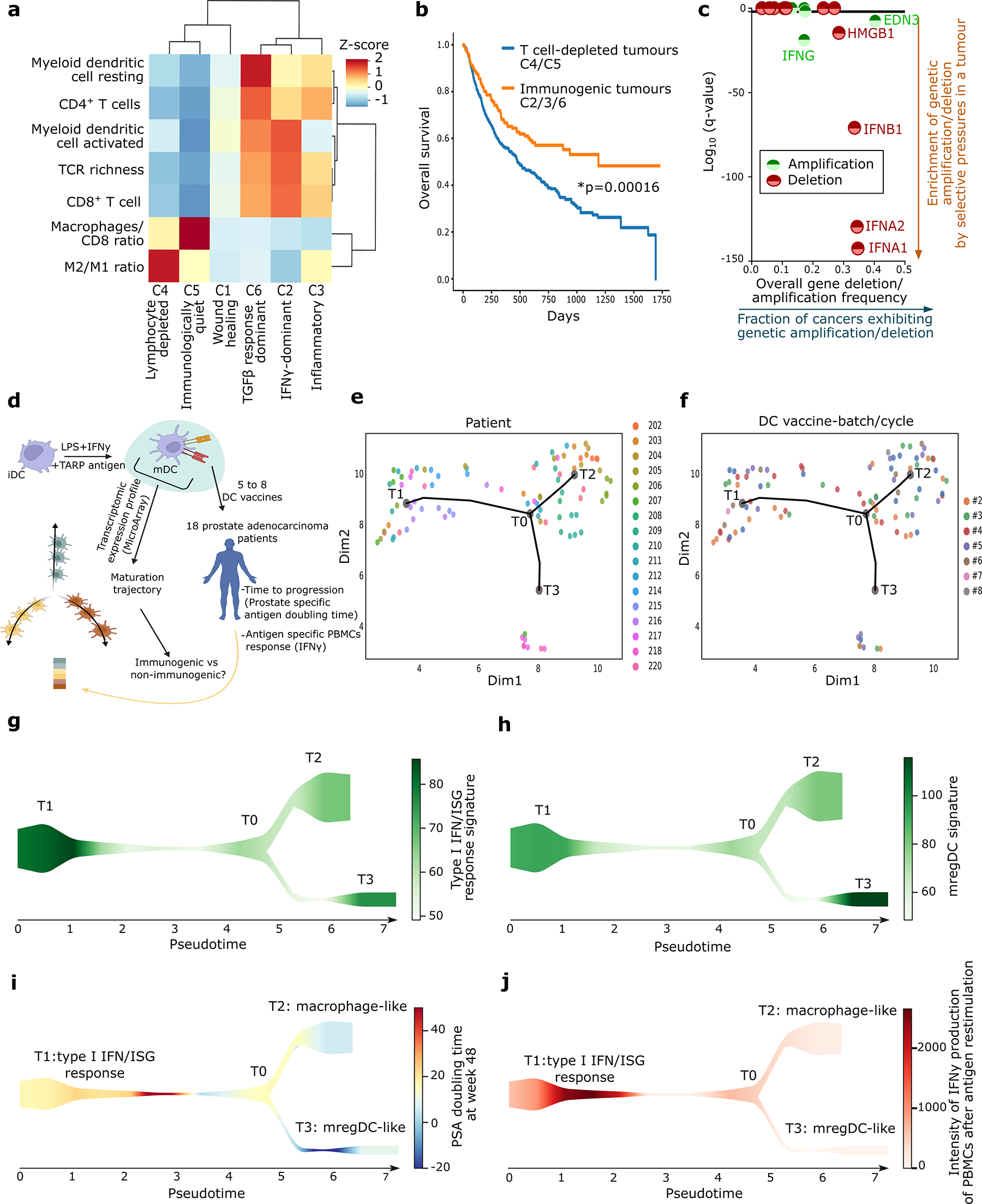
Type I interferon (IFN) pathway disparities distinguish T cell-depleted tumours and are essential for the most clinically efficacious maturation-trajectory of human autologous DC vaccines. (A) Heatmap representation of CIBERSORT deconvolution across all TCGA cancer types. Population abundances were row-normalised over the six immune subtype clusters (C1, n=1313; C2, n=1210, C3, n=688; C4, n=222, C5, n=2; C6, n=111). (B) Kaplan-Meier curve of the overall survival of an integrated multi-cancer dataset (across 12 clinical trials) of patients’ transcriptome-profiled before ICBs-treatment (anti-PD1, anti-CTLA4 or anti-PDL1 ICBs, or combinations thereof) sub-grouped in T cell-depleted C4/C5 tumours (n=667) and immunogenic C2/C3/C6 tumours (n=474). Statistical significance was assessed with the log rank test. (C) Pan-cancer GISTIC 2.0 analysis of the deletion and amplification of the 12 genes selected from the network analysis. Statistical significance was defined as FDR<0.05 (random permutations to background score distribution, BH-adjusted). bladder cancer, n=136; breast cancer, n=880; colorectal adenocarcinomas, n=585; glioblastoma multiforme, n=580; head and neck squamous cell cancer, n=310; kidney clear cell carcinoma, n=497; acute myeloid leukaemia, n=200; lung adenocarcinoma, n=357; lung squamous cell carcinoma, n=344; ovarian serous carcinoma, n=563; uterine endometrial carcinoma, n=496. (D-J) STREAM DC vaccine trajectory analysis of 93 autologous DC vaccines pulsed with LPS, IFNγ and TARP peptide generated for 18 prostate adenocarcinoma cancer patients vaccinated with 5-8 vaccines. (D) Overview of STREAM DC vaccine trajectory analysis. (E-F) Pseudo time was inferred from the transcriptome of the DC vaccines based on variable genes. The principal graph was initiated with epg_alpha=0.01, epg_mu=0.2, epg_lambda=0.03 and epg_n_nodes=5. Dots depict individual DC vaccines and dot colour represents (E) patient number or (F) DC vaccine batch/cycle. (G-H) Signature scores were overlaid on the graph as streamplots. Type I IFN/ISG response signature (G) or mature regulatory DC signature (H) were used as colour intensity of the resulting stream plots. (I-J) Patient outcome measurements were overlaid on the graph as streamplots. PSA doubling time at week 48 (I) and intensity of IFNγ production of peripheral blood mononuclear cell after antigen restimulation (J) were used as colour intensity of the resulting stream plots. See also figure S1.

Next, we looked at the transcriptomic inverse correlation between expression of ligand-receptor, cell-receptors, or cell-ligand, pairs in TCGA C4/C5-tumours e.g., high expression of ligand-coding genes vs. low expression of receptor-coding genes, or vice versa, for all pairs (Eddy et al., 2020; Ramilowski et al., 2015). Such inverse correlation within each pair was visually illustrated by node-to-node connecting arrows in a network (**Suppl. Fig.S1A**), highlighting the immune dysregulation in inter/extra-cellular signalling, as established previously (Eddy et al., 2020; Ramilowski et al., 2015). TCGA C4/C5-tumours exhibited an inverse correlation between various pairs relevant for danger signalling (Garg and Agostinis, 2017) e.g., the TLR4 pathway (inversely correlating with various immune cells or danger signals/cytokines), and type I interferon (IFN) pathway (inverse correlation between *IFNA2* and *IFNAR1*), amongst others (**Suppl. Fig.S1A**). Next, we pursued copy number-variation (CNV) analyses for loci-centric amplification or deletion events of genes short-listed by the above network, to extract the most dominant genetic deletion pressures in pan-cancer TCGA data (Mermel et al., 2011). This revealed that type I IFN-related genes (*IFNA1*/2, *IFNB1*) were under the highest genetic deletion pressures (**Fig.1C**). Overall, this emphasized that human T cell-depleted tumours harbour disparities in immune cell-composition (DCs and T cells’ depletion and enrichment of M2-like macrophages) and danger signalling (dysregulation of TLR4-driven and type I IFN pathways).

### Type I IFN-response distinguishes the most immunogenic and clinically efficacious maturation-trajectory of human autologous DC vaccines

Above results suggested that TLR4-agonist/IFN-stimulated DC vaccines could be a practical formulation to rejuvenate anti-cancer immunity. However, previous clinical trials have applied DC vaccines stimulated with type I/II IFNs (i.e., IFNα/β/γ, inducing interferon-stimulated genetic [ISG]-response) and/or TLR4-agonists (e.g., bacterial LPS, that also stimulates ISG-responses) yet with only limited clinical success (Sprooten et al., 2019a). This made us question if, despite TLR4/IFN-stimulation, there might be heterogeneity in the ‘maturation trajectories’ of human DC vaccines that undermines their efficacy.

To address this, we analysed the single DC vaccine transcriptomes from a clinical trial, wherein DC vaccines were TLR4/IFN-stimulated and applied against ICB-resistant and/or T cell-depleted tumours. Therein 18 prostate cancer patients were treated with 6-8 cycles of autologous moDC vaccines pulsed with TARP antigen-peptide and stimulated with LPS + IFNγ (**Fig.1D**) (Castiello et al., 2017). Herein each DC vaccine’s bulk-transcriptome and matching patient response was available (prostate-specific antigen [PSA] doubling time, as time-to-tumour progression [TTP]); and IFNγ production by TARP antigen-pulsed PBMCs as antigen-specific immune response (Castiello et al., 2017).

We performed an unbiased pseudo-time trajectory analysis via STREAM (Chen et al., 2019) on transcriptomes of 93 DC vaccines (**Fig.1D**). This revealed that despite a simple maturation-stimuli and a homogenous antigen-source (peptide rather than lysate), DC vaccines had three completely distinct trajectories (T1, T2 or T3) (**Fig.1E-F**). Herein while the distribution of patients was relatively trajectory-specific (**Fig.1E**), DC vaccine batches/cycles (**Fig.1F**) were mostly randomly distributed. Next, we performed a REACTOME biological pathway enrichment per trajectory with remarkable results. We found that T1 enriched for type I IFN-response and immunogenic/pro-inflammatory signalling (*IRF3, IFI44L, MAVS, CCR7, GBP4/5, MX2, RELB, OASL, HLA-DRA, CD40, TNFSF4*) (**Suppl. Fig.S1B-C**), T2 enriched for macrophage-like pathways (phagocytosis, scavenger receptor pathway, M2 macrophages; *CD68, CXCL1, CXCL12, CCL26, CEACAM3, CCL2, CCR1*) (**Suppl. Fig.S1B, S1D**), while T3 was enriched for hyper-maturation and immune-checkpoint signalling (*CLEC10A, TLR5, TGFA, SIGLEC6/9/12, CD36, IL12R, IL13RA1, ENTPD1, IL1R1, CMTM6, TNFRSF10B, REL, S1PR1*) (**Suppl. Fig.S1B, S1E**).

An anti-inflammatory macrophage-like T2 trajectory might be explained by the monocytic origins of the DC vaccines (Kvedaraite and Ginhoux, 2022). However, the T1 implied a specific ISG-response^+^DC vaccine-subset whereas T3 implied maturation-associated regulatory (mreg)-DC program (i.e., hyper-maturation accompanied by immune-checkpoints signalling) (Kvedaraite and Ginhoux, 2022; Maier et al., 2020). To verify these, we applied previously published ISG response-signature (**Supplementary Table 1**) and mregDC-signature (**Supplementary Table 1**) to these trajectories (Echebli et al., 2018; Maier et al., 2020). Indeed, the ISG response-signature had the highest enrichment in T1 (**Fig.1G**) while the mregDC-signature had the highest enrichment in T3 (**Fig.1H**).

These distinct DC vaccine trajectories were unprecedented and hence we pursued their alignment with tumoural (TTP) (**Fig.1I**) or antigen-specific (PBMC) responses to DC vaccines (**Fig.1J**). We found that the T1-trajectory enriched for high antigen-specific responses and longer TTP whereas the T3-trajectory didn’t enrich for antigen-specific responses while associating with shorter TTP (**Fig.1I-J**). The macrophage-like T2 trajectory resembled T3 trends to a larger extent. These data suggested that DC vaccines form contradictory maturation-trajectories such that a potential DCvax-IT approach must favour type I IFN/ISG-responses over macrophage-like or mregDC-like states.

### Murine TC1-tumours phenocopy key immune-disparities of human T cell-depleted tumours

To formulate a preclinical DCvax-IT, it was necessary to delineate a pre-clinical syngeneic tumour model mimicking the major immune-disparities of human T cell-depleted tumours. Therefore, we conducted a reanalysis of an existing dataset composed of tumour transcriptomes of 11 commonly used (subcutaneous) murine tumour models in immuno-oncology (Mosely et al., 2017) (**Fig.2A**). This comparative expression analyses involved genetic signatures (**Supplementary Table 1**) for pro-lymphocytic IFNγ/effector signalling, macrophages, type I IFN/ISG-response, or DCs (Mosely et al., 2017; Sprooten et al., 2021). This revealed that the triple-oncogene driven c-H-Ras^+^HPV16-E6^+^HPV16-E7^+^TC1-tumours (Smahel et al., 2001) (cancerous lung epithelia-origin, C57BL/6 mice) recapitulated most features of human T cell-depleted tumours (negligible levels of IFNγ/effector signalling, type I IFN/ISG-response and DCs, counter-balanced by enrichment of the macrophage-signature) (**Fig.2A**).

**Figure 2.**
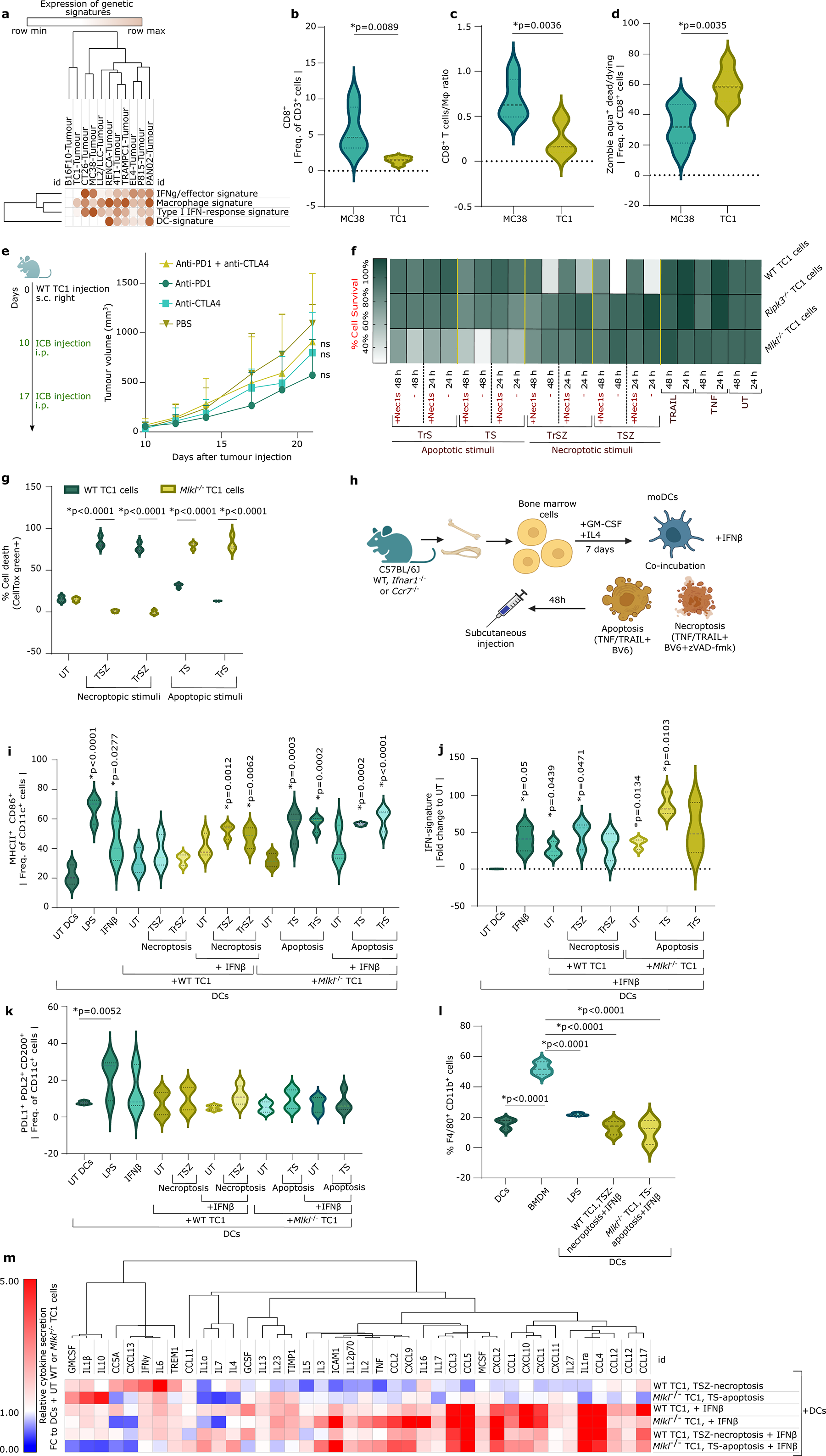
Optimization of DCvax-IT co-integrating immunogenic cell death and interferon(IFN)β-stimulation for the human T cell-depleted tumours phenocopy: murine TC1-tumours. (A) Heatmap representation of metagene expression for 4 distinct signatures for the different subcutaneous tumours derived from different cell lines derived from the existing micro-array dataset GSE85509 (B-D) Violin plots of the flow cytometry analysis of CD45^+^ cell fraction obtained from subcutaneous MC38 and wild-type TC1 tumours on day 23 after tumour cell injection showing (B) the percentage of CD8^+^ of CD3^+^ cells (C) CD8^+^ T cell-to-macrophage ratio and (D) percentage of zombie aqua^+^ death/dying in CD8^+^ T cells. (n=6; two-tailed student’s t test) (E) Tumour volume curve of wild-type TC1 tumour-bearing mice treated with anti-PD1 and/or anti-CTLA4 on day 9 and 16 after wild-type TC1 injection. (n=6; area under curve; One-way ANOVA, Kruskal-Wallis test) (F) Heatmap of survival of wild-type, *Ripk3*^-/-^ and *Mlkl*^-/-^ TC1 cells 24h and 48h after treatment with different cell death stimuli with or without Nec1s. Values represent 3 to 4 repeats (G) Cell death of wild-type and *Mlkl*^-/-^ TC1 cells 48h after treated with different cell death stimuli. p-values depict comparison between wild-type and *Mlkl^-/-^* TC1 cells. (n=3; Two-way ANOVA, Sidak’s multiple comparisons test) (H) Schematic overview of the vaccine formulation process. (I-M) Functional analysis of dendritic cells untreated or stimulated with LPS, IFNβ or with untreated or dying TC1 cancer cells with or without IFNβ. (I) Flow cytometry analysis of DC maturation assessed by MHCII^+^ CD86^+^ frequency of CD11c^+^. p-values depict comparison to UT DCs. (n=3; one-way ANOVA, Dunnet’s multiple comparisons test) (J) IFN-signature expression measured by qPCR. p-values depict comparison to UT DCs (n=3; One sample t test). (K) Flow cytometry analysis of frequency of PDL1^+^PDL2^+^CD200^+^ of CD11c^+^ cells in moDCs alone or cocultured with untreated or dying wild type or *Mlkl^-/-^* TC1 cells. p-values depict comparison to UT moDCs (n=4, LPS/IFNβ n=3; One-way ANOVA, Fischer LSD). (L) Flow cytometry analysis of frequency of CD11b^+^F4/80^+^ fractions in moDCs (alone or cocultured with untreated or dying wild type or *Mlkl^-/-^* TC1 cells) or bone-marrow derived macrophages (BMDMs). p-values depict comparison to BMDMs (n=3; one-way ANOVA, Dunnet’s multiple comparisons test) (M) Cytokine secretion assessed by cytokine array. From all values, the background was subtracted. Normalisation was done using moDCs stimulated with untreated cancer cells. (n=3) See also figure S2-S9

Next, we pursued tumour immunophenotyping to functionally validate the choice of TC1-tumours and selected the immunogenic MC38-tumours (Efremova et al., 2018) (cancerous colon epithelia-origin, C57BL/6 mice) for comparison (**Fig.2A**). Compared to MC38-tumours, TC1-tumours had significantly lower CD8^+^CD3^+^T cell-infiltrates (**Fig.2B; Suppl. Fig.S2A**), lower CD8^+^T cell-to-TAMs (CD11b^+^F4/80^+^) ratio (**Fig.2C; Suppl. Fig.S2B**) and a higher CD8^+^T cell death (**Fig.2D; Suppl. Fig.S2A**). This suggested that TC1-tumours harbour a T cell-depleted, pro-TAMs environment. This was further supported by TC1-tumours being non-responsive to anti-PD1 or anti-CTLA4 ICBs, or a combination thereof (**Fig.2E**). Finally, in line with defective type I IFN/TLR danger signalling in human C4/C5-tumours, TC1 cancer cells failed to significantly secrete IFNα/β upon stimulation with agonists of TLR4 (LPS), TLR7 (Imiquimod), RIG-I (5’ppp-dsRNA), STING/cGAS (2’3’cGAMP) and immunogenic (doxorubicin) or non-immunogenic (cisplatin) chemotherapies (**Suppl. Fig.S3A-B**). This underscored the suitability of TC1-tumours as pre-clinical representative of human T cell-depleted C4/C5-tumours.

### TC1-cells undergo apoptotic or necroptotic immunogenic cell death with antigen release

We explored the possibility of providing damage-associated molecular patterns (DAMPs)-based danger signalling and antigen release-based ‘DC pulsing’ via immunogenic cell death (ICD) in TC1 cancer cells (Garg et al., 2017; Perez and De Palma, 2019). ICD is well-established to potentiate anti-cancer DC vaccines (Garg et al., 2016). However, it is currently unknown which underlying cell death pathway (i.e., apoptotic vs. necroptotic) has the strongest potentiating effect (Galluzzi et al., 2020).

To distinguish apoptosis vs. necroptosis, we utilized consensus cell death receptors (DR)-driven necroptotic stimuli (TNF/TRAIL + SMAC mimetic, BV6 + pan-caspase inhibitor, zVAD-fmk) vs. apoptotic stimuli (TNF/TRAIL + BV6) (Galluzzi et al., 2018; Yang et al., 2016). Herein, TNF or TRAIL were comparatively used as DR-agonists because they have distinct pro-inflammatory activity (Yi et al., 2018). Wild-type (WT) TC1 cells exhibited a substantial loss of survival following necroptotic stimuli, which was inhibited by the pharmacological RIPK1-inhibitor, Nec1s (Takahashi et al., 2012). WT TC1 cells did not respond to apoptotic stimuli (**Fig.2F**). To overcome this resistance, we used CRISPR/Cas9-driven knock-out of two essential necroptosis regulators i.e., *Ripk3* or *Mlkl* (**Suppl. Fig.S3C-D**) (Yang et al., 2016). *Ripk3*^-/-^TC1 cells were resistant to both necroptotic and apoptotic stimuli (**Fig.2F**). Instead, *Mlkl*^-/-^ TC1 cells showed resistance to necroptotic-stimuli but reduction in cellular survival following apoptotic stimuli (**Fig.2F**). Subsequently, WT TC1 cells underwent necroptotic cell death (**Fig.2G**) based on: the absence of caspase-3/7 activity (**Suppl. Fig.S3E**), non-sustained annexin-V staining (**Suppl. Fig.S4A-B**) and phosphorylation of MLKL (**Suppl. Fig.S4E-F**) (hereafter referred to as necroptotic TC1 cells or necroptotic^TC1^). Conversely *Mlkl*^-/-^TC1 cells underwent apoptotic cell death (**Fig.2G**) based on: significant caspase-3/7 activity (**Suppl. Fig.S3E**), sustained annexin-V staining (**Suppl. Fig.S4C-D**) and the absence of MLKL phosphorylation (**Suppl. Fig.S4E-F**) (hereafter referred to as apoptotic TC1 cells or apoptotic^TC1^). Of note, we didn’t explicitly use *Casp8*^-/-^TC1 cells as necroptotic system because, like WT TC1 cells with TS+zVAD-fmk, *Casp8*^-/-^ or *Casp*8^-/-^*Casp*9^-/-^ (but not *Casp9*^-/-^) TC1 cells (**Suppl. Fig.S5A**) underwent Nec1s-inhibitable loss of cellular survival upon TS-stimuli (**Suppl. Fig.S5B**), thereby making the latter redundant (and thus unnecessary) as necroptosis-model.

Finally, we analysed the ICD-like DAMP-profiles of cells stimulated as described above (Galluzzi et al., 2020). Interestingly, both apoptotic/necroptotic TC1 cells showed ICD-like DAMPs features i.e., higher enrichment of surface-calreticulin^HIGH^CD47^LOW^cells (**Suppl. Fig.S6A**), high ATP (**Suppl. Fig.S6B-C**) or HMGB1 (**Suppl. Fig.S6D**) release, and high extracellular release of the TC1-specific E7 antigen (**Suppl. Fig.S6D**), wherein necroptotic^TC1^ had a relatively superior DAMP-profile than apoptotic^TC1^. Nevertheless, neither apoptosis nor necroptosis could induce IFNβ secretion from TC1 cells (*data not shown*). Thus, TC1 cells showed proficient DAMP-based apoptotic/necroptotic ICD coupled with antigen-release.

### DCvax-IT co-integrating ICD and IFNβ-stimulation favour type I IFN-responses over macrophages-like or mregDC-like phenotypes

We based our DC vaccines on bone marrow-derived moDCs (**Fig.2H**). Since IFN/ISG responses were pre-requisite for immunogenicity, we did a comparative potency analysis between IFNα/β/γ (due to lack of consensus on prioritization (Sprooten et al., 2019b)) for induction of DC phenotypic maturation (MHC-II^+^CD86^+^CD11c^+^) (**Suppl. Fig.S7A**) and secretion of an ISG-factor, CXCL10. Herein, *E.coli* LPS served as a positive control. Interestingly, IFNβ had the highest potency in inducing DC phenotypic-maturation (**Suppl. Fig.S7B**) and CXCL10 secretion (**Suppl. Fig.S7C**). We selected 2.5 ng/mL IFNβ for rest of the study since it achieved sufficient phenotypic maturation and CXCL10 release. Here, 2.5 ng/mL IFNβ also up-regulated multiple ISGs in DCs: *Irf7, Rsad, Mx1, Cxcl9* or *Cxcl10* (**Suppl. Fig.S7D**).

Subsequently, we pulsed moDCs with apoptotic/necroptotic^TC1^, with or without IFNβ-stimulation (**Fig.2H**) and checked for: successful pulsing (efferocytosis of apoptotic/necroptotic^TC1^ by DCs), DC phenotypic maturation, bulk IFN/ISG-response genetic signature, macrophage-like/mregDC-like phenotype and increase in CCR7^+^DCs (necessary for LN-homing). After pulsing with apoptotic^TC1^ alone, we already saw promising DC stimulation i.e., high efferocytosis of TC1 cells (**Suppl. Fig.S8A**) and high DC phenotypic maturation (**Fig.2I**), which was further optimized or maintained by IFNβ co-stimulation (**Suppl. Fig.S8B;** **Fig.2I**). Co-stimulation with IFNβ was especially useful for necroptotic^TC1^ since it outperformed DC phenotypic maturation by necroptotic^TC1^ alone (**Fig.2I**) (although efferocytosis for necroptotic^TC1^ was lower/more variable; **Suppl. Fig.S8C-D**). Moreover, the combination of ICD and IFNβ increased or maintained CCR7-levels on DCs, especially for apoptotic^TC1^ (**Suppl. Fig.S8E-F**). Thus along with ICD, co-stimulation with IFNβ optimized DC maturation.

Next, to confirm type I IFN/ISG-response, we investigated the ISG-signature behaviour (metagene for *Irf7, Rsad, Mx1, Cxcl9* and *Cxcl10*) in the DC+TC1+IFNβ co-cultures. Herein, IFNβ-induced ISG-response was maintained in necroptotic^TC1^+DCs co-cultures but was potentiated in apoptotic^TC1^+DCs co-cultures (**Fig.2J**). Moreover, TNF-driven apoptosis/necroptosis was better at preserving or potentiating ISG-response than TRAIL (**Fig.2J**). Overall, above data made us prioritize TNF-driven cell death, DCs and IFNβ co-cultures as putative DCvax-IT formulations. Next, we checked for macrophages-like or mregDCs-like phenotypes. Analyses for PDL1^+^PDL2^+^CD200^+^CD11c^+^mregDC (Maier et al., 2020) (**Suppl. Fig.S8G**) showed that LPS (significantly), and to a variable extent IFNβ (non-significantly), enriched for a mregDC-like phenotype (**Fig.2K**). However, the presence of cancer cells (untreated, apoptotic/necroptotic) avoided the mregDC-like phenotype (**Fig.2K**). In line with this, apoptotic^TC1^/necroptotic^TC1^DCvax-IT did not enrich CD11b^+^F4/80^+^macrophage-like markers as compared to *bona fide* (M-CSF-based) bone marrow-derived macrophages (BMDMs) (**Fig.2L**).

Finally, we aimed to unbiasedly validate DCvax-IT secretome’s immunogenic/inflammatory potential via an antibody array combined with GSEA-based Gene Ontology biological pathway analyses. Interestingly, necroptotic^TC1^+DCs secretome was more inflammatory than apoptotic^TC1^+DCs (**Fig.2M**), with enrichment of inflammatory cytokine-related pathways (**Suppl. Fig.S9A-B**). However, the enrichment of diverse pro-inflammatory cytokines was mainly enhanced by IFNβ co-stimulation (**Suppl. Fig S9C-E**) (including immunogenic/ISG-response factors like CXCL10, CXCL9, CCL5, IL12p70) in TC1-DCs co-cultures (**Fig.2M**). The co-stimulation with ICD and IFNβ enriched for specific inflammatory pathways depending on cell death pathway (**Suppl. Fig.S9D-E**). Thus, we successfully optimized a pre-clinical DCvax-IT (integrating cancer cells undergoing apoptotic/necroptotic-ICD and IFNβ-driven type I IFN/ISG stimulation of DCs) that resembled the immunogenic-trajectory of human DC vaccines.

### DCvax-IT induce type I IFN sensing-dependent anti-cancer immunity *in vivo*

For rapid *in vivo* immunogenicity testing of DCvax-IT, we first used a prophylactic vaccination setting (**Fig.3A**). Herein, injections with both apoptotic^TC1^/necroptotic^TC1^ DCvax-IT significantly protected mice from later tumour challenge (compared to PBS-treatment) (**Fig.3A**), which was comparable to antigenic peptides-based DC vaccine (E6/E7 peptides + IFNβ + DCs) (**Fig.3A**). Additionally, when above DCvax-IT vaccinated mice, with tumour-free survival, were re-challenged with TC1 cancer cells (for assessment of stable memory responses), there was no significant tumour formation (**Fig.3B**). This showed that DCvax-IT creates *in vivo* immunogenicity-driven memory responses.

**Figure 3.**
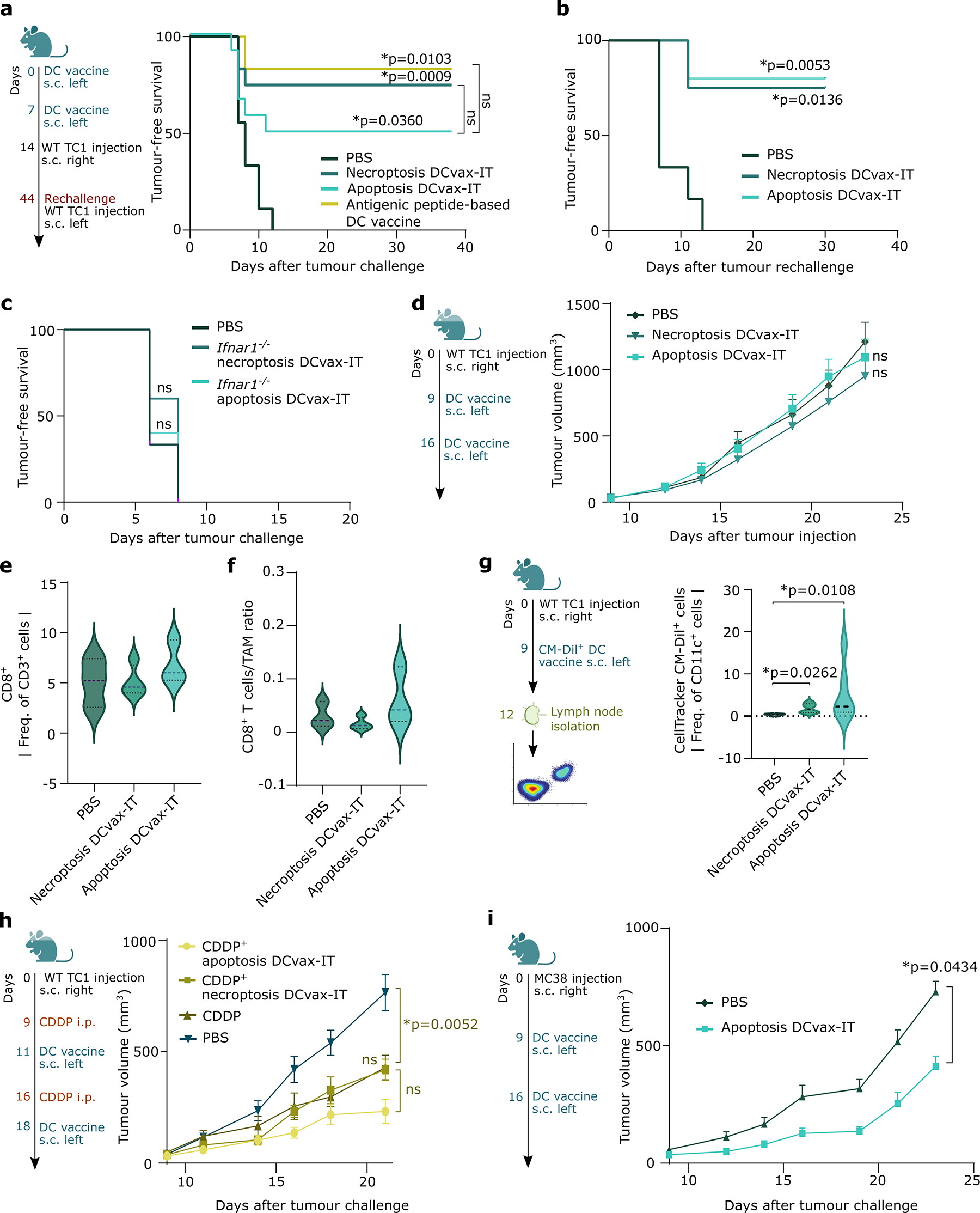
DCvax-IT induces anticancer immunity *in vivo* but fails against T cell-depleted tumours in a curative set-up. (A-C) Kaplan-Meier curves showing survival of mice vaccinated with 2 doses of prophylactic DC vaccines administered at day 0 and 7, followed by subcutaneous wild-type TC1 tumour injection. p-values depict comparison to PBS treated mice. (A) Comparison of necroptosis DCvax-IT, Apoptosis DCvax-IT and antigenic peptide-based DC vaccine to PBS treated mice. (PBS; n=9, DCvax-IT; n=6, Log-rank (Mantel-Cox) test) (B) Rechallenge with wild-type TC1 injection after vaccination with necroptosis DCvax-IT, Apoptosis DCvax-IT wild-type DC vaccines. (PBS,n=6; Apoptosis DCvax-IT, n=5; Necroptosis DCvax-IT, n=4, Log-rank (Mantel-Cox) test) (C) Comparison of necroptosis DCvax-IT, Apoptosis DCvax-IT *Ifnar1-/-* DC vaccines to PBS treated mice. (PBS; n=6, Apoptosis DCvax-IT, Necroptosis DCvax-IT: n=5, Log-rank (Mantel-Cox) test) (D) Tumour volume curve of wild-type TC1 tumour-bearing mice treated with DCvax-IT on day 9 and 16 after wild-type TC1 injection. comparison to PBS treated mice (n=12, area under curve; Kruskal-Wallis test) (E-F) Violin plots of flow cytometry analysis of CD45^+^ cell fraction obtained from untreated or DCvax-IT-treated subcutaneous wild-type TC1 tumours on day 23 after tumour cell injection: (E) Frequency of CD8^+^ T cells or (F) CD8^+^ T cells-to-TAM ratio. comparison to PBS treated mice (UT; n=3, Necroptosis DCvax-IT; n=4, Apoptosis DCvax-IT; n=3, one-way ANOVA, Dunnet’s multiple comparisons test) (G) Frequency of Celltracker CM-Dil^+^ CD11c^+^ cells in the lymph nodes of DCvax-IT vaccinated mice. p-values depict comparison to PBS treated mice (UT; n=4, Necroptosis DCvax-IT; n=6, Apoptosis DCvax-IT; n=6, one-way ANOVA, Kruskal-Wallis test) (H) Tumour volume curve of wild-type TC1 tumour bearing mice treated with cisplatin on day 9 and 16 alone or in combination with DCvax-IT on day 11 and 18 and after wild-type TC1 injection. p-values depict comparison to cisplatin treated mice. (n=8; area under curve, one-way ANOVA, Dunnet’s multiple comparisons test) (I) Tumour volume curves of MC38 tumour bearing mice treated with apoptosis DCvax-IT on day 9 and 16 after MC38 injection. p-values depict comparison to PBS treated mice. (PBS,n=8; Apoptosis DCvax-IT; n=10, area under curve; Mann-Whitney test) See also figure S9-S11

Next, we wondered if type I IFN-sensing (i.e., interaction of IFNβ with its cognate receptor complex of IFNAR1::IFNAR2 on DCs) was indeed necessary for the induction of anti-cancer immunity by DCvax-IT, especially beyond ICD. Hence, we pursued testing of apoptotic^TC1^/necroptotic^TC1^DCvax-IT with *Ifnar1*^-/-^DCs. The efferocytosis potential of *Ifnar1*^-/-^DCs for apoptotic^TC1^/necroptotic^TC1^ was measurable (**Suppl. Fig.S10A-B**) but their phenotypic-maturation in the DCvax-IT setting was not proficient (**Suppl. Fig.S10C**). Thus, mice vaccinated with *Ifnar1*^-/-^DCs-based apoptotic^TC1^/necroptotic^TC1^DCvax-IT failed to resist tumour challenge (**Fig.3C**). Altogether, DCvax-IT induced proficient *in vivo* anti-cancer immunity that was dependent on the DCs’ ability to sense IFNβ, irrespective of ICD/cell death.

### Therapeutic DCvax-IT fails against a T cell-depleted tumour but effectively overcomes a T cell-infiltrated tumour

For pre-clinical testing, we pursued the curative or therapeutic vaccination setting (**Fig.3D**). Surprisingly, DCvax-IT did not decrease TC1-tumour growth (**Fig.3D**). On the level of tumour immunophenotype, DCvax-IT failed to significantly increase CD8^+^T cell-infiltrates (**Fig.3E**), or substantially shift the CD8^+^T cell-to-TAMs ratio (**Fig.3F**), thereby emphasizing that despite high immunogenic potential, DCvax-IT failed to sufficiently activate anti-tumour immunity.

It was not clear if the ineffectiveness of the therapeutic DCvax-IT was due to the vaccine formulation or the immune-disparities in TC1-tumours. The moDCs may exhibit limited LN-homing (Perez and De Palma, 2019), which can be particularly troublesome in a therapeutic (rather than prophylactic) setting, because pre-existing tumours can further inhibit moDC migration to the LNs (Imai et al., 2012; Ito et al., 2006). Hence, we fluorescently labelled the DCvax-IT via CellTracker™CM-DiI to investigate their mobility. Labelled cells were injected in tumour-bearing mice and their accumulation in inguinal/axillary LNs draining the subcutaneous injection-site was measured (**Fig.3G**). There was significant LN-enrichment of CM-DiI^+^DCvax-IT in treated mice (**Fig.3G**).

Given that LN-homing of DCvax-IT was not impaired, we next investigated if the immuno-resistant burden of TC1-tumours was too high to be overcome by monotherapy, therefore requiring a combinatorial chemotherapy for debulking. However, combining DCvax-IT with cisplatin, a chemotherapy with immunotherapy-synergizing properties in TC1-tumours (Beyranvand Nejad et al., 2020; Bruchard et al., 2022), did not add significant therapeutic efficacy beyond cisplatin’s own tumour regressing capacity (**Fig.3H**). We wondered if the non-immunogenic nature of TC1-tumour formed a specific barrier against DCvax-IT. Hence, we therapeutically applied DCvax-IT against the more T cell-enriched MC38-tumours. Of note, MC38 cells were only susceptible to apoptotic but not necroptotic-stimuli (**Suppl. Fig.S11A-B**), as reported previously (Yu et al., 2022). We therefore focused on apoptotic^MC38^DCvax-IT. Here, we observed that DCs efficiently executed efferocytosis (**Suppl. Fig.S11C-D**) and phenotypically matured (**Suppl. Fig.S11E**) when exposed to apoptotic^MC38^ plus IFNβ. Interestingly, apoptotic^MC38^DCvax-IT significantly reduced MC38-tumour growth (**Fig.3I**). Thus, despite the high immunogenicity and proficient LN-homing abilities of DCvax-IT, T cell-depleted (but not T cell-infiltrated) tumours were immuno-resistant to DCvax-IT.

### TC1-tumours enrich for M2-like PDL1^+^TAMs that partly suppress T cells

To improve therapeutic DCvax-IT efficacy, it was critical to find the dominant immuno-resistant pathway in the TC1-tumours and neutralize it. Hence, we pursued differential gene-expression (DGE) analyses between TC1 (DCvax-IT immuno-resistant) and MC38 (DCvax-IT susceptible) tumours using above bulk-tumour transcriptome data (**Fig.2A**). The DGE analysis highlighted a set of major immune-relevant genes (see **Supplementary Table 2**) that disclosed enrichment of myeloid anti-inflammatory (*Axl, Gas2, Areg, Figf, Ptgs1/2, Thbs1/2*) and TAM-associated genes (*Cd33, Mrc1*) in TC1-tumours (**Fig.4A)**. MC38-tumours enriched for IFN/ISG-response, T cell-relevant or effector signalling-linked genes (*Oas2/l2/l1, Rsad2, Stat1/2, Ccl5, Mx1/2, Gzmf/d/b, Cxcl9/10*). *In vivo* tumour immunophenotyping confirmed TC1-tumours had relatively higher M2-to-M1 TAM-ratio (**Fig.4B**) and significantly lower M1-like MHC-II^HIGH^CD206^LOW^TAMs, when compared to MC38-tumours (**Fig.4C**). Of note, TC1-tumours enriched higher TAMs than DCs or cDC1 and cDC2 subsets (**Suppl. Fig.S12A-B**).

**Figure 4.**
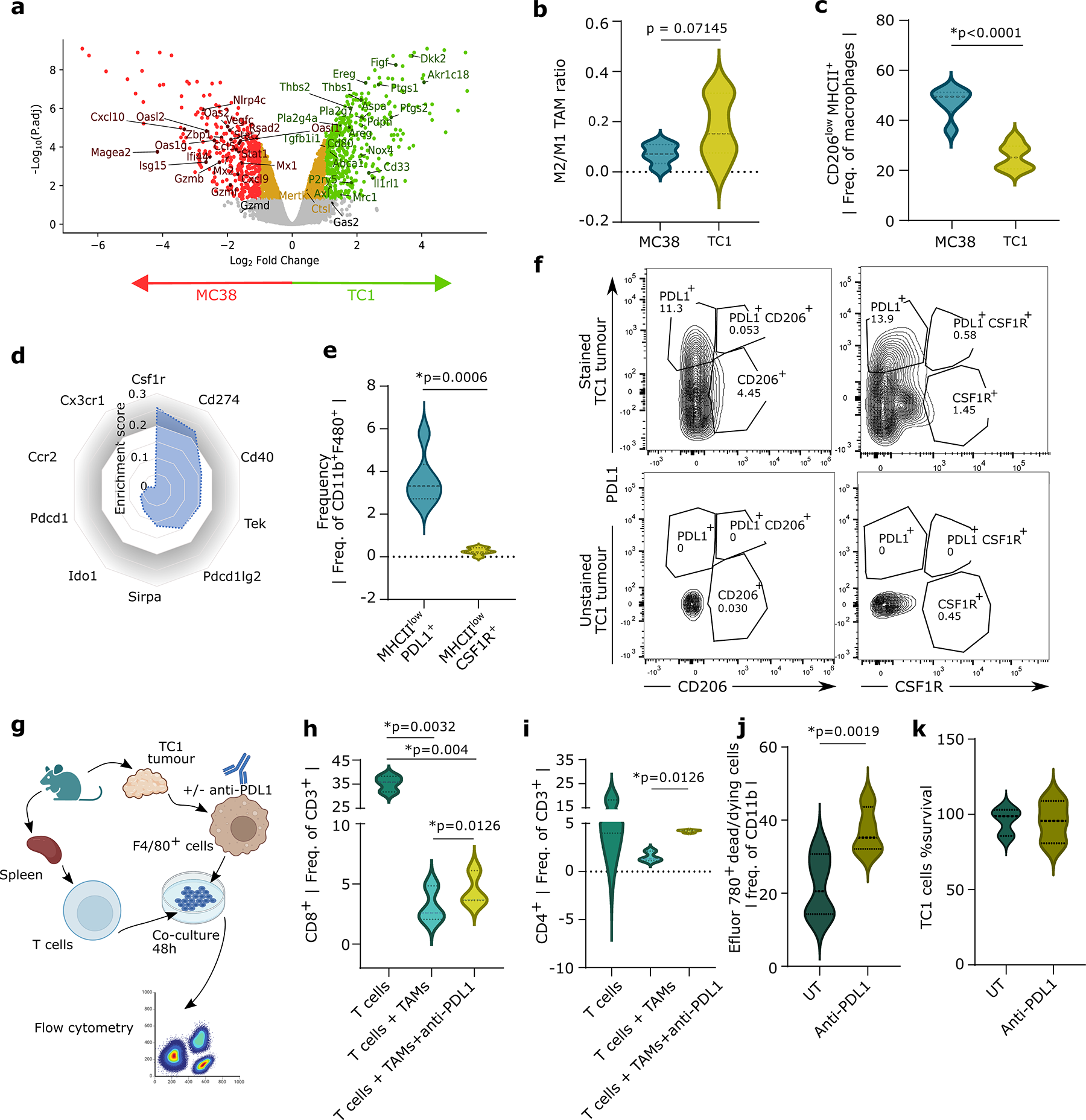
TC1-tumours enrich for M2-like PDL1^+^ tumour associated macrophages (TAMs) that partly antagonize T cells. (A) Volcano plot capturing fold changes and statistical significance of gene expression alterations between MC38 and wild-type TC1 subcutaneous tumours in micro-array (GSE85509). (B-C) Violin plots of the CD45^+^ cell fraction obtained from subcutaneous MC38 and TC1 tumours on day 23 after tumour cell injection. (B) M2/M1 ratio present in MC38 and wild-type TC1 tumours (TC1; n=5, MC38; n=6; two-tailed student’s t test) (C) Frequency of CD206^low^ MHCII^high^ cells of CD11b^+^ F4/80^+^ (n=6; two-tailed student’s t test) (D) Radar plot illustrating enrichment scores of indicated genes compared to a reference macrophage transcriptomic profile (see materials & methods for more details on computational approach). (E) Violin plot of the CD45^+^ cell fraction obtained from subcutaneous wild-type TC1 tumours on day 23 after tumour cell injection. Frequency of TAM subtypes. (n=6; two-tailed paired t test) (F) Contour plots of flow cytometry analysis of PDL1^+^, CSF1R^+^ and CD206^+^ (gating based on unstained samples) on TAM derived from wild-type TC1 tumours isolated on day 23 after tumour cell injection. (G-I) Flow cytometry analysis of T cell recovery, involving cocultures of TAMs isolated from wild-type TC1 subcutaneous tumours on day 23 after tumour cell injection and pre-incubated with or without anti-PDL1 for 48h, together with paired T cells obtained from the spleen (G) Schematic representation of TAM/T cell coculture experimental setup. (H-I) Frequency of (H) CD8^+^ T cells and (I) CD4^+^ T cells. (n=3; two-tailed paired t test) (J) Flow cytometry analysis of Efluor 780^+^ dead/dying cell in untreated and anti-PDL1 treated wild-type TC1-derived TAMS isolated on day 23 after tumour cell injection (n=4; two-tailed paired t-test) (K) Survival of untreated and anti-PDL1 treated wild-type TC1 cells measured by MTT. (n=3; two-tailed paired t test) See also figure S12

Several TAM-linked targets are currently undergoing translational analyses e.g., CSF1R, CCR2, CX3CR1, SIRPA, TIE2 (*Tek*), PDL1 (*Cd274*), PDL2 (*Pdcd1lg2*), CD40, IDO1, and PD1 (*Pdcd1*) (Goswami et al., 2022; Nakamura and Smyth, 2020). To prioritise the most dominant targets, we used the differential TC1-specific myeloid genes (**Fig.4A**) as input for a correlation-GSEA against a reference murine macrophage transcriptome-profile (Vandenbon et al., 2016). This predicted that *Csf1r*, followed by *Cd274* (PDL1) (**Fig.4D**), might have the highest associations with TC1-tumour associated myeloid genes. However, tumour immunophenotyping emphasized that TC1-tumours were dominated by immunoregulatory (MHC-II^LOW^) TAMs expressing significant PDL1 (PDLI^+^MHC-II^LOW^) rather than CSF1R (CSF1R^+^MHC-II^LOW^) (**Fig.4E**). In TC1-tumours, PDLI^+^MHC-II^LOW^TAMs were strongly enriched and were distinct from CSF1R^+^MHC-II^LOW^TAMs or the canonical M2-like CD206^+^MHC-II^LOW^TAMs (**Fig.4F**). Also, there were more PDL1^+^TAMs than PDL1^+^DCs, PDL1^+^cDC1, PDL1^+^cDC2 (**Suppl. Fig.S12C**) or PDL1^+^TC1 cells (**Suppl. Fig.S12D**) thus positioning TAMs as dominant source of tumoural PDL1 signalling.

M2-like TAMs are well-established T cell suppressors (Güç and Pollard, 2021; Huber et al., 2010) wherein PDL1^+^TAMs can reduce T cell viability (Shan et al., 2020). To evaluate if this was applicable here, we isolated TAMs from TC1-tumours and splenic lymphocytes from the same mice and co-cultured them with or without PDL1 blocking antibody (anti-PDL1 ICB) (**Fig.4G**). Expectedly, the presence of TAMs dramatically reduced the recovery of live CD8^+^/CD4^+^T cells while anti-PDL1 ICB partially but significantly increased live CD8^+^/CD4^+^T cell-recovery (**Fig.4H-I**). This phenotype was PDL1-specific, rather than dependent on the classical PDL1::PD1 interactions (Wei et al., 2018), because anti-PD1 ICB (**Suppl. Fig.S12E-F**) failed to increase T cell-recovery (**Suppl. Fig.S12F**). Thus, we wondered how anti-PDL1 ICB was modulating macrophage’s function (Cha et al., 2019; Hartley et al., 2018) and tested effects of PDL1 blockade on macrophage/TAM’s M2/M1 polarization. Anti-PDL1 ICB did not significantly affect the M2-like (MHC-II^LOW^CD206^HIGH^) vs. M1-like (MHC-II^HIGH^CD206^LOW^) ratio for macrophages (**Suppl. Fig.S12G**) or TAMs (**Suppl. Fig.S12H**). PDL1 can modulate survival of various cells including myeloid or cancer cells (Cha et al., 2019; Hartley et al., 2018). Accordingly, we saw that TAMs underwent significant cell death upon anti-PDL1 ICB-treatment (**Fig.4J**). Moreover, this pro-death effect of anti-PDL1 ICB was TAMs-centric, because TC1 cells did not change their survival upon anti-PDL1 ICB-treatment (**Fig.4K**). Thus, TC1-tumours harbour M2-like PDL1^+^TAMs that partly suppress T cells *ex vivo*, such that this suppression can be ameliorated via pro-death activity of anti-PDL1 ICB against TAMs.

### DCvax-IT and anti-PDL1 ICB synergistically suppress TC1-tumours by overcoming immunosuppressive macrophages

Since PDL1^+^TAMs were partly T cell-suppressive and susceptible to pro-death depletion via anti-PDL1 ICB, we tested if treating TC1-tumours with anti-PDL1 ICB can suppress tumour growth irrespective of DCvax-IT (**Fig.5A**). While anti-PDL1 ICB indeed slightly suppressed TC1-tumour growth this was not significant (**Fig.5A**). The combination of anti-PDL1 ICB with both apoptotic^TC1^ and necroptotic^TC1^DCvax-IT synergistically enforced significantly greater TC1-tumour suppression (**Fig.5A**), thereby highlighting the importance of PDL1-signalling as a barrier to DCvax-IT.

**Figure 5.**
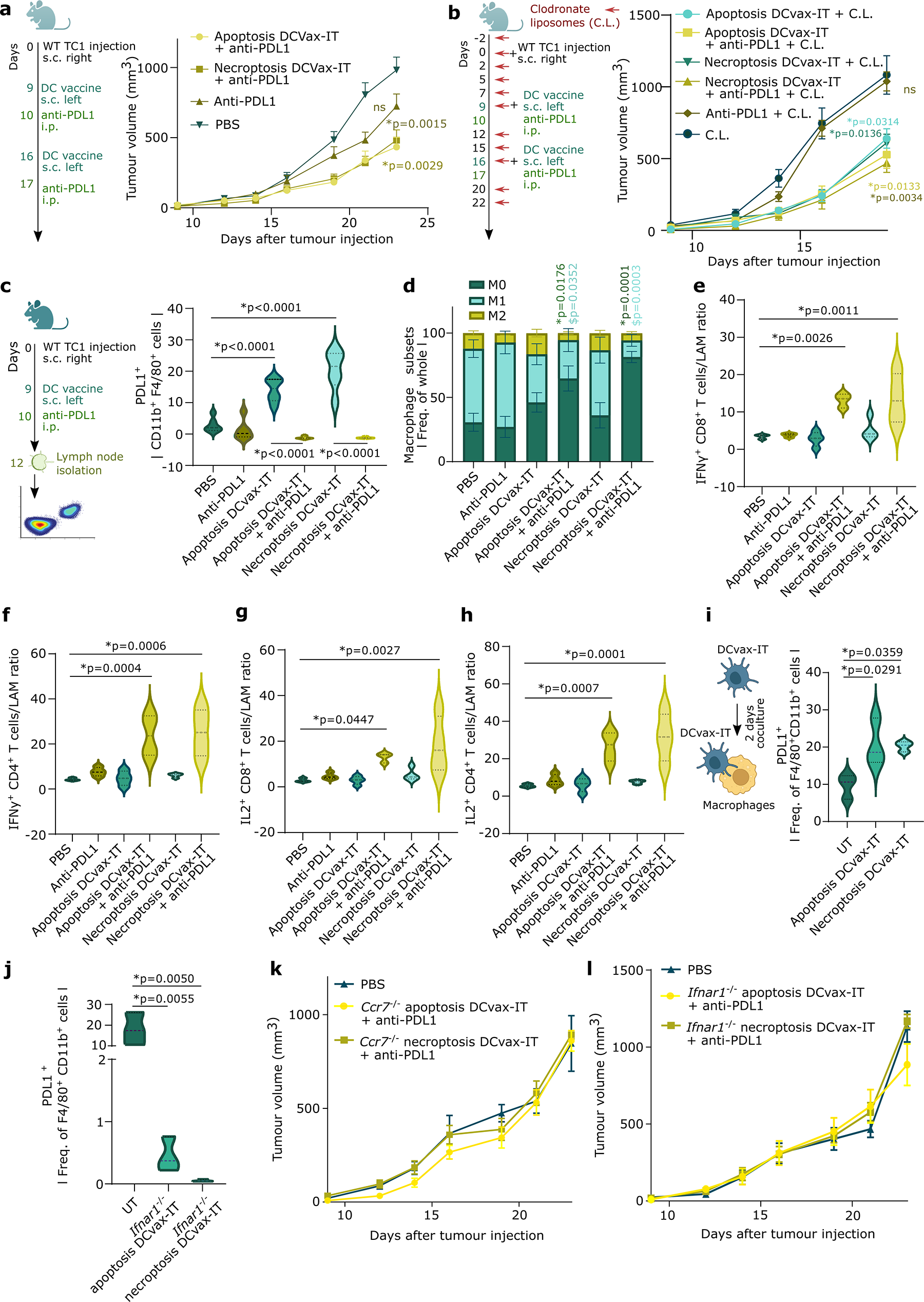
DCvax-IT mobilizes PDL1^+^macrophages in lymph nodes which can be overcome by DCvax-IT and anti-PDL1 ICB combination. (A) Tumour volume curve of wild-type TC1 tumour bearing mice treated with DCvax-IT on day 9 and 16 and/or anti-PDL1 on day 10 and 17. p-values depict comparison to PBS treated mice. (PBS; n=12, Apoptosis DCvax-IT + anti-PDL1/Necroptosis DCvax-IT + anti-PDL1; n=12, area under curve, one-way ANOVA Kruskal-Wallis test) (B) Tumour volume curve of wild-type TC1 tumour bearing mice treated with DCvax-IT on day 9 and 16 and/oranti-PDL1 on day 10 and 17 in combination with clodronate liposomes (C.L.). p-values depict comparison to clodronate liposome treated mice. (clodronate liposome/Apoptosis DCvax-IT/Necroptosis DCvax-IT; n=5, PDL1/Apoptosis DCvax-IT + anti-PDL1/Necroptosis DCvax-IT+ anti-PDL1; n=6, area under curve; one-way ANOVA, Dunnet’s multiple comparisons test) (C-H) Lymph node analysis of wild-type TC1 tumour bearing mice, 3 days after treatment with DCvax-IT with or without anti-PDL1. (C) Frequency of PDL1^+^ of CD11b^+^F4/80^+^ cells. p-values depict comparison to PBS treated mice, unless otherwise specified. (n=6; one-way ANOVA, Dunnet’s multiple comparisons test) (D) Frequency of M1 (MHC-II^HIGH^CD206^LOW^), M2 (MHC-II^LOW^CD206^HIGH^) or M0 (MHC-II^LOW^CD206^LOW^) macrophage-subsets within the whole macrophage population. *p-values depict comparison of M0 to PBS treated mice. ^$^p-values depict comparison of M1 to PBS treated mice. (n=6; two-tailed student’s t test) (E) IFNγ^+^ CD8^+^ T cells-to-TAMs ratio. p-values depict comparison to PBS treated mice. (PBS/ Apoptosis DCvax-IT + anti-PDL1; n=3, anti-PDL1/Apoptosis DCvax-IT /Necroptosis DCvax-IT+ anti-PDL1;n=4, One-way ANOVA, Fischer’s LSD test) (F) IFNγ^+^ CD4^+^ T cells-to-TAMs ratio. p-values depict comparison to PBS treated mice. (PBS; n=3 anti-PDL1/Apoptosis DCvax-IT/ Apoptosis DCvax-IT + anti-PDL1/Necroptosis DCvax-IT+ anti-PDL1;n=4, One-way ANOVA, Fischer’s LSD test) (G) IL2^+^ CD8^+^ T cells-to-TAMs ratio. p-values depict comparison to PBS treated mice. (PBS/ Apoptosis DCvax-IT + anti-PDL1; n=3, anti-PDL1/Apoptosis DCvax-IT /Necroptosis DCvax-IT+ anti-PDL1;n=4, One-way ANOVA, Fischer’s LSD test) (H) IL2^+^ CD4^+^ T cells-to-TAMs ratio. p-values depict comparison to PBS treated mice. p-values depict comparison to PBS treated mice. (PBS; n=3 anti-PDL1/Apoptosis DCvax-IT/ Apoptosis DCvax-IT + anti-PDL1/Necroptosis DCvax-IT+ anti-PDL1;n=4, One-way ANOVA, Fischer’s LSD test) (I-J) Frequency of PDL1^+^ cells of CD11b^+^ F4/80^+^ cells after coculturing BMDMs with (I) wild-type or (J) *Ifnar1^-/-^* DCvax-IT for 48h. p-values depict comparison to untreated DCs. (n=3; one-way ANOVA, Dunnet’s multiple comparisons test) (K-L) Tumour volume curve of wild-type TC1 tumour bearing mice treated with (K) *Ccr7*^-/-^ (L) or *Ifnar1*^-/-^ DCvax-IT on day 9 and 16 and with anti-PDL1 on day 10 and 17. p-values depict comparison to PBS treated mice (n=6; area under curve; One-way ANOVA, Kruskal-Wallis test) See figure S13-14

Since the immunomodulation via PDL1 was supposedly independent of PD1-engagement, and more dominant than CSF1R, we combined anti-PD1 or anti-CSF1R ICBs with DCvax-IT. Indeed, unlike anti-PDL1 ICB (**Fig.5A**), both anti-PD1 (**Suppl. Fig.S13A**) and anti-CSF1R (**Suppl. Fig.S13B**) ICBs did not synergize with DCvax-IT thereby excluding any roles for PD1::PDL1 interactions or CSF1R^+^TAMs. Of note, anti-CSF1R ICB did not suppress TC1-tumour growth despite achieving significant depletion of CSF1R^+^TAMs (**Suppl. Fig.S13C**) without disturbing the general TAMs-compartment (**Suppl. Fig.S13D**), thereby emphasizing the complexities of TAMs heterogeneity.

Above results positioned TAMs/macrophages as the key immuno-resistance barrier to DCvax-IT, that could be overcome with anti-PDL1 ICB-based targeting of PDL1^+^TAMs. To confirm this association, we depleted the macrophages via clodronate liposomes (C.L.) (**Fig.5B**) (Nguyen et al., 2021). Herein, C.L. were applied at a carefully titrated dose that significantly depleted macrophages (**Suppl. Fig.S13E**) but did not affect DCs (**Suppl. Fig.S13F**). Importantly, C.L.-based macrophage depletion completely reshaped the immunotherapy-susceptibility of TC1-tumours. Upfront, after macrophages depletion, the DCvax-IT were able to autonomously exert a significant suppressive effect on TC1-tumour growth (**Fig.5B**), thereby confirming macrophages as the main immuno-resistant barrier to DCvax-IT. Contrastingly, the superiority of DCvax-IT plus anti-PDL1 ICB synergism over DCvax-IT alone completely disappeared after macrophages were depleted (**Fig.5B**). This confirmed the macrophage-targeting properties of anti-PDL1 ICB as the primary reason behind its synergism with DCvax-IT. Accordingly, under C.L.-based macrophage depletion, DCvax-IT were able to autonomously boost the tumoural CD8^+^T cells (**Suppl. Fig.S13G**). This stimulus to CD8^+^T cell infiltration was not further potentiated by DCvax-IT plus anti-PDL1 ICB (**Suppl. Fig.S13G**). In conclusion, DCvax-IT plus anti-PDL1 ICB combo significantly suppressed TC1-tumours; such that the failure of DCvax-IT alone, and low tumoural CD8^+^T cells were all attributable to macrophage-mediated immunosuppression.

### DCvax-IT induce PDL1^+^macrophages in lymph nodes and combination with anti-PDL1 ICB depletes them to facilitate effector T cells

It has been well-established that higher PDL1 abundance can proportionally improve anti-PDL1/PD1 ICB-efficacy (Davis and Patel, 2019). This made us question if DCvax-IT were in fact further boosting PDL1^+^macrophages, thereby allowing PDL1 signalling to reach critical abundance thresholds that are therapeutically relevant for synergism.

DC vaccines’ major *modus operandi* involves anti-tumour priming of LN-associated T cells (Garg et al., 2017; Perez and De Palma, 2019; Sabado et al., 2017), however very little is known about the impact of DC vaccines on LN-associated macrophages (LAMs). Since DCvax-IT showed good LN-homing, we investigated if they perhaps induced PDL1^+^LAMs. We immunophenotyped the inguinal/axillary LNs (that drain the subcutaneous vaccination-site) of tumour-bearing mice (**Fig.5C**). While DCvax-IT did not directly cause mobilization of macrophages (**Suppl. Fig.S14A**) or cDC1/cDC2 (**Suppl. Fig.S14B-C**) in LNs, they did induce a strong enrichment of PDL1^+^LAMs (**Fig.5C**). Similar enrichment was not observed for CSF1R^+^LAMs (**Suppl. Fig.S14D**), PDL1^+^cDC1 (**Suppl. Fig.S14E**) or PDL1^+^cDC2 (**Suppl. Fig.S14F**). However, DCvax-IT plus anti-PDL1 ICB (but not anti-PDL1 ICB alone) caused a remarkably strong depletion of not only PDL1^+^LAMs (**Fig.5C**), but also other myeloid cells i.e., macrophages (**Suppl. Fig.S14A**), cDC1/cDC2 (**Suppl. Fig.S14B-C**), CSF1R^+^LAMs (**Suppl. Fig.S14D**) and PDL1^+^cDC1/cDC2 (**Suppl. Fig.S14E-F**), thereby indicating PDL1-specific as well as bystander modulation of the LN myeloid environment. Indeed, anti-PDL1 ICB plus DCvax-IT caused significant reduction in both M1-like and M2-like LAMs thereby ‘re-wiring’ overall LAMs-compartment toward homeostatic M0-like LAMs (MHC-II^LOW^CD206^LOW^) (**Fig.5D**). Contrastingly, anti-PDL1 ICB plus DCvax-IT had no clear increase in phenotypic-maturation of cDC1/cDC2 (**Suppl. Fig.S14G-H**) thereby suggesting a macrophage-centric modulation of LN-associated inflammation.

Next, we analysed the LN-associated T cells and their effector cytokine-activity (IFNγ/IL2 production). Upfront, none of the treatment conditions caused any enrichment of LN-associated CD4^+^/CD8^+^T cells. (**Suppl. Fig.S14I-J**). Hence, we focused on the qualitative re-structuring of the LN immune-milieu from the viewpoint of IFNγ^+^ or IL2^+^ CD4^+^/CD8^+^T cells exceeding LAMs-enrichments. Notably, DCvax-IT or anti-PDL1 ICB alone did not enrich for IFNγ^+^CD4^+^/CD8^+^T cells (**Fig.5E-F**) or IL2^+^CD4^+^/CD8^+^T cells (**Fig.5G-H**) compared to LAMs, thus substantiating their therapeutic failure. Remarkably, DCvax-IT plus anti-PDL1 ICB strongly mobilized IFNγ^+^CD4^+^/CD8^+^T cells (**Fig.5E-F**) and IL2^+^CD4^+^/CD8^+^T cells (**Fig.5E-F**) thereby supporting the combination’s anti-tumour efficacy.

Classically, T cell-derived IFNγ is a dominant inducer of PDL1 (Cha et al., 2019). Since DCvax-IT alone did not enrich for IFNγ^+^CD4^+^/CD8^+^T cells (**Fig.5E-F**), we questioned if DCvax-IT were directly inducing PDL1 on macrophages (**Fig.2M**). Interestingly, co-culturing macrophages with DCvax-IT resulted in significantly more PDL1^+^macrophages (**Fig.5I**), but not CSF1R^+^macrophages (**Suppl. Fig.S14K**). Since DCvax-IT secreted several type I IFN/ISG-response driven cytokines (**Fig.2M**) that can redundantly induce PDL1 on macrophages (Cha et al., 2019; Zerdes et al., 2018), we ablated this pathway by using type I IFN non-responsive *Ifnar1*^-/-^DCvax-IT. Confirmatively, *Ifnar1*^-/-^DCvax-IT did not induce PDL1 on macrophages (**Fig.5J**). This suggested that DCvax-IT’s IFNβ-driven pro-inflammatory/immunogenic activity (**Fig.2M**) created PDL1^+^macrophages in LNs.

To mechanistically confirm the direct involvement of DCvax-IT’s IFNβ-driven immunogenicity and their eventual LN-homing (via CCR7) in enhancing PDL1-based immuno-resistance, we repeated the therapeutic experiments with *Ifnar1*^-/-^DCvax-IT or *Ccr7*^-/-^DCvax-IT, respectively. Expectedly, *Ccr7*^-/-^ DCvax-IT (**Fig.5K**) or *Ifnar1*^-/-^DCvax-IT (**Fig.5L**) failed to synergize with anti-PDL1 ICBs in regressing TC1-tumours. This confirmed that DCvax-IT induces PDL1-based immuno-suppression (led by PDL1^+^LAMs) in LNs, that limits effector CD4^+^/CD8^+^T cells, thus creating an immuno-resistant macrophage niche susceptible to only anti-PDL1 ICBs.

### DCvax-IT mobilizes PDL1^+^TAMs and combination with anti-PDL1 ICB depletes them to promote anti-tumour T cell immunity

Since TC1-tumours pre-enriched for T cell-suppressive PDL1^+^TAMs, we wondered if DCvax-IT also enhanced PDL1^+^TAMs to weaken anti-tumour T cell immunity. Indeed, DCvax-IT themselves caused a significant enrichment of not only TAMs (**Fig.6A**) but also PDL1^+^TAMs (**Fig.6B**). In line with its effects in LNs, DCvax-IT plus anti-PDL1 ICB (but not anti-PDL1 ICB alone) caused depletion of not only TAMs (**Fig.6A**), but also PDL1^+^TAMs (**Fig.6B**). An increase in the CD8^+^T cells-to-TAMs ratio was also observed (**Fig.6C**). This confirmed that DCvax-IT facilitates both PDL1^+^LAMs and PDL1^+^TAMs, thereby creating a PDL1-dominant immunosuppressive circuit traversing both LNs and tumours, with specific susceptibility to combinatorial anti-PDL1 ICB.

**Figure 6.**
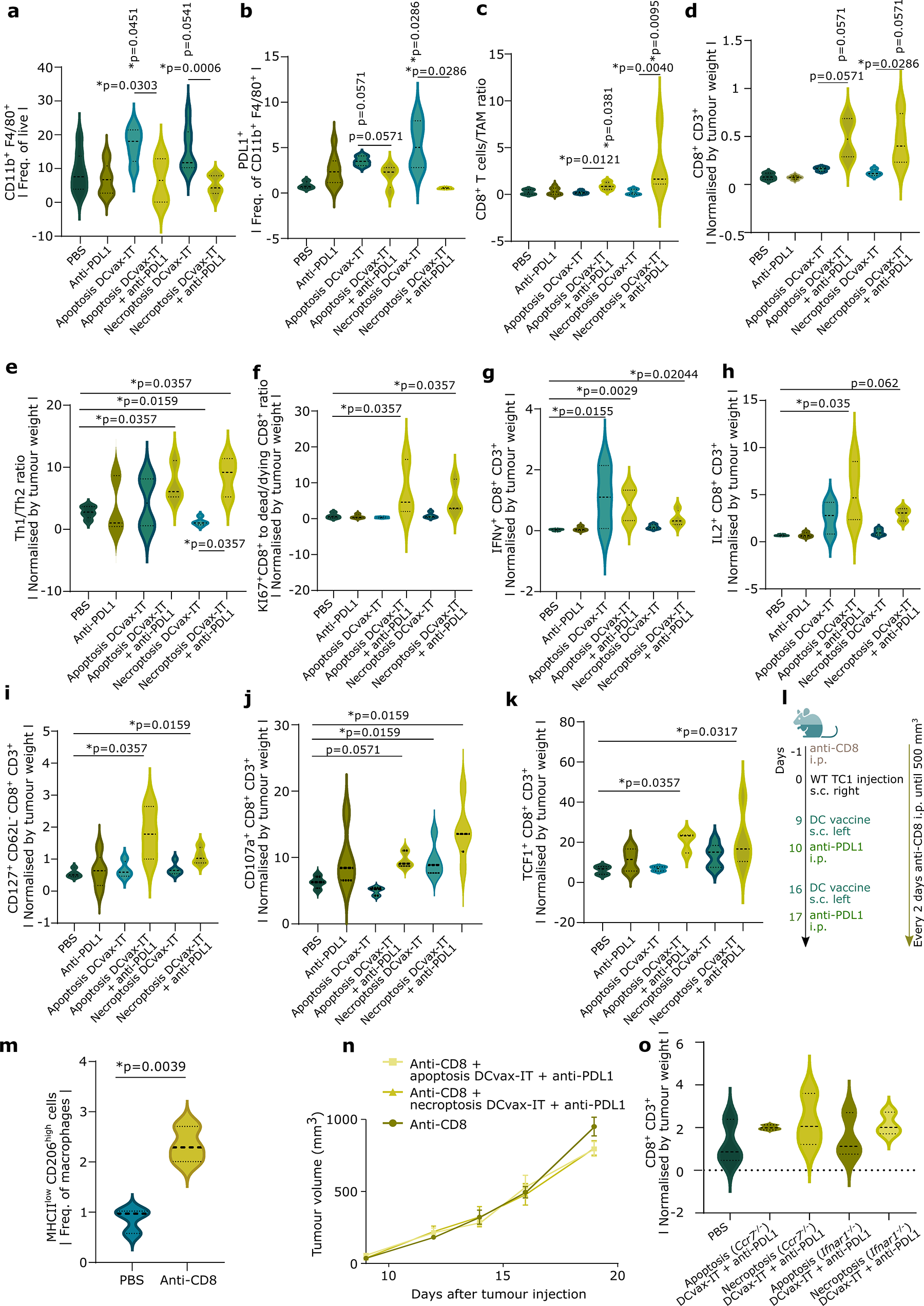
DCvax-IT mobilizes PDL1^+^TAMs and combination with anti-PDL1 immune checkpoint blockade depletes them to facilitate Th1 and effector/stem-like memory CD8^+^T cells. (A-C) Flow cytometry TIL analysis of the CD45^+^ fraction from subcutaneous wild-type TC1 tumours isolated on day 23 after tumour cell injection, treated with DCvax-IT on day 9 and 16 and/or anti-PDL1 on day 10 and 17. (A) Percentage of TAMs assessed by CD11b^+^ and F4/80^+^ (B) Percentage of PDL1^+^ TAMS (C) CD8^+^ to TAM ratio (A-C) p-values depict comparison to PBS treated mice, unless otherwise specified. (n=4-9; Mann-Whitney test) (D-J) Flow cytometry TIL analysis of the CD45^+^ fraction from wild-type TC1 tumour on day 23 after tumour injection treated with DCvax-IT on day 9 and 16 with or without anti-PDL1 injections on day 10 and 17. normalised by tumour weight at day of isolation. (D) Th1-to Th2 ratio. (E) KI67^+^CD8^+^ to dead CD8^+^ ratio. (F)IFNγ^+^ of CD8^+^ T cells. (G) IL2^+^ of CD8^+^ T cells. (H) CD127^+^ CD62^-^ ofcells. (I) CD107a^+^ of CD8^+^ cells. (J) TCF^+^ of CD8^+^ cells. (D,E,H,I,J) p-values depict comparison to PBS treated mice, unless otherwise specified. (n=3-5; Mann-Whitney test) (F-G) p-values depict comparison to PBS treated mice, unless otherwise specified. (n=3-4; One-way ANOVA, Kruskal-Wallis test) (K-L) Wild-type TC1 tumour bearing mice were treated with DCvax-IT on day 9 and 11, with anti-PDL1 on day 10 and 11 and with anti-CD8 one day before tumour injection and every other day until 500mm^3^. p-values depict comparison to PBS treated mice, unless otherwise specified. (K) Percentage of MHCII^low^ CD206^high^ of CD11b^+^ F4/80^+^ in the CD45^+^ fraction from wild-type TC1 tumour on day 23 after tumour injection. (n=3; two-tailed student’s t test) (L) Tumour volume curve (n=7; area under curve; One-way ANOVA, Kruskal-Wallis test) (M) TIL analysis of the CD45^+^ fraction isolated from wild-type TC1 tumour on day 23 after tumour injection. Tumours were treated with *Ifnar1^-/-^* or *Ccr7^-/-^* DCvax-it on day 9 and 16 in combination with anti-PDL1 injection on day 10 and 17. Frequency of CD8^+^ of CD3^+^ cells. Comparison to PBS treated mice. (n=3; One-way ANOVA, Kruskal-Wallis test) See also figure S15-S16

Confirmatively, we did not see any differences in tumoural enrichment of DCs, cDC1, cDC2, PDL1^+^cDC1 or PDL1^+^cDC2 (**Suppl. Fig.S15A-E**). Similarly, there was no significant differences in phenotypic maturation of DCs, cDC1 or cDC2 (**Suppl. Fig.S15F-H**).

We further analysed the tumoural T cell-infiltrates. Upfront, only DCvax-IT plus anti-PDL1 ICB caused a significant enrichment of CD8^+^T cells (**Fig.6D**). Contrastingly, all conditions caused a non-significant increase in CD4^+^T cell-infiltrates (**Suppl. Fig.S15I**). Unlike CD8^+^T cells, CD4^+^T cell-compartment is simultaneously composed of both anti-tumorigenic T helper type-1 (Th1) cells as well as pro-tumorigenic/pro-TAMs Th type-2 (Th2) cells (Thorsson et al., 2018). Hence, we analysed the Th1-to-Th2 ratio, and found that only DCvax-IT plus anti-PDL1 ICB significantly skewed the tumoural CD4^+^T cell-milieu toward the Th1 cells (**Fig.6E**).

Next, we analysed the survival, effector function, and tumour-sensitisation states of CD8^+^T cells. Only the DCvax-IT plus anti-PDL1 ICB, but not the individual monotherapies, increased the proliferative Ki67^+^CD8^+^T cells over dead/dying CD8^+^T cells (**Fig.6F**). Accordingly, only this combination caused significant and consistent enrichment of all effector CD8^+^T cell-subsets, compared to PBS-treated mice (Blank et al., 2019; McLane et al., 2019; Naulaerts et al., 2021) i.e., effector IFNγ^+^ (**Fig.6G**) or IL2^+^ (**Fig.6H**) CD8^+^T cells, effector-memory CD127^+^CD62L^-^CD8^+^T cells (**Fig.6I**), and cytotoxic CD107a^+^CD8^+^T cells (**Fig.6J**). Finally, we pondered if the combination increased tumour-sensitisation of CD8^+^T cells marked by enrichment of chronically exhausted and stem-like memory CD8^+^T cells (Blank et al., 2019; McLane et al., 2019; Naulaerts et al., 2021; Oliveira et al., 2021). We observed that the combination indeed increased enrichment of exhausted PD1^+^TIM3^+^CD8^+^T cells (**Suppl. Fig.S15J**), although its strength was variable. In parallel, we noticed a significant enrichment of stem-like memory TCF1^+^CD8^+^T cells, compared to PBS-treated mice (**Fig.6K**), that took place without increasing terminally differentiated (memory) EOMES^+^TCF1^-^CD8^+^T cells (**Suppl. Fig.S15K**). This proved that DCvax-IT plus anti-PDL1 ICB’s deleterious effects on TAMs are paralleled by an anti-tumorigenic re-orientation of the T cell-compartment.

To confirm the M2-like TAMs vs. CD8^+^T cell antagonistic relationship, we depleted CD8^+^T cells via anti-CD8 depleting antibodies (**Fig.6L**). Interestingly, TC1-tumors depleted of CD8^+^T cells (**Suppl. Fig.S16A**) showed higher M2-like TAMs-enrichment (**Fig.6M**) and lower enrichment of M1-like TAMs (**Suppl. Fig.S16B**). This was accompanied by a much faster tumour growth (**Suppl. Fig.S16C**). This emphasized an antagonism between M2-like TAMs and CD8^+^T cells and indicated an association between enrichment of M2-like TAMs and TC1-tumour’s accelerated growth. As expected, the anti-tumour effect of DCvax-IT plus anti-PDL1 ICB disappeared in absence of CD8^+^T cells (**Fig.6N**). Finally, to confirm the pro-CD8^+^T cells role for DCvax-IT’s type I IFN-based immunogenicity and LN-homing, independent of anti-PDL1 ICB, we immunophenotyped the tumours from above therapeutic experiments administering *Ifnar1*^-/-^DCvax-IT or *Ccr7*^-/-^DCvax-IT plus anti-PDL1 ICB. Confirmatively, despite combination with anti-PDL1 ICB, *Ccr7*^-/-^DCvax-IT or *Ifnar1*^-/-^DCvax-IT (**Fig.6O**) did not enrich CD8^+^T cells. Above data confirmed that DCvax-IT mobilizes PDL1^+^macrophage circuit spanning both tumours and LNs thereby suppressing T cell immunity, such that combination with anti-PDL1 ICB depletes these TAMs and revives anti-tumour T cell immunity.

### PDL1 and TAMs co-association is a negative prognostic niche in human cancers that specifically predicts clinical response to anti-PDL1 ICBs

Above observations deserved further physiological or clinical validation. Herein, we wished to use above TC1-tumour derived data as putative ‘biomarker profiles’ for clinical validation. Accordingly, a large-scale bioinformatics analysis with 21 existing single-cell (sc)RNAseq datasets spanning 13 diverse cancer-types, 287 patients and 2 species (humans or mice) showed that, at single-cell resolution, TAMs where the most dominant source of *CD274* (PDL1) (**Fig.7A**), amongst various immune, stromal or cancer cells.

**Figure 7.**
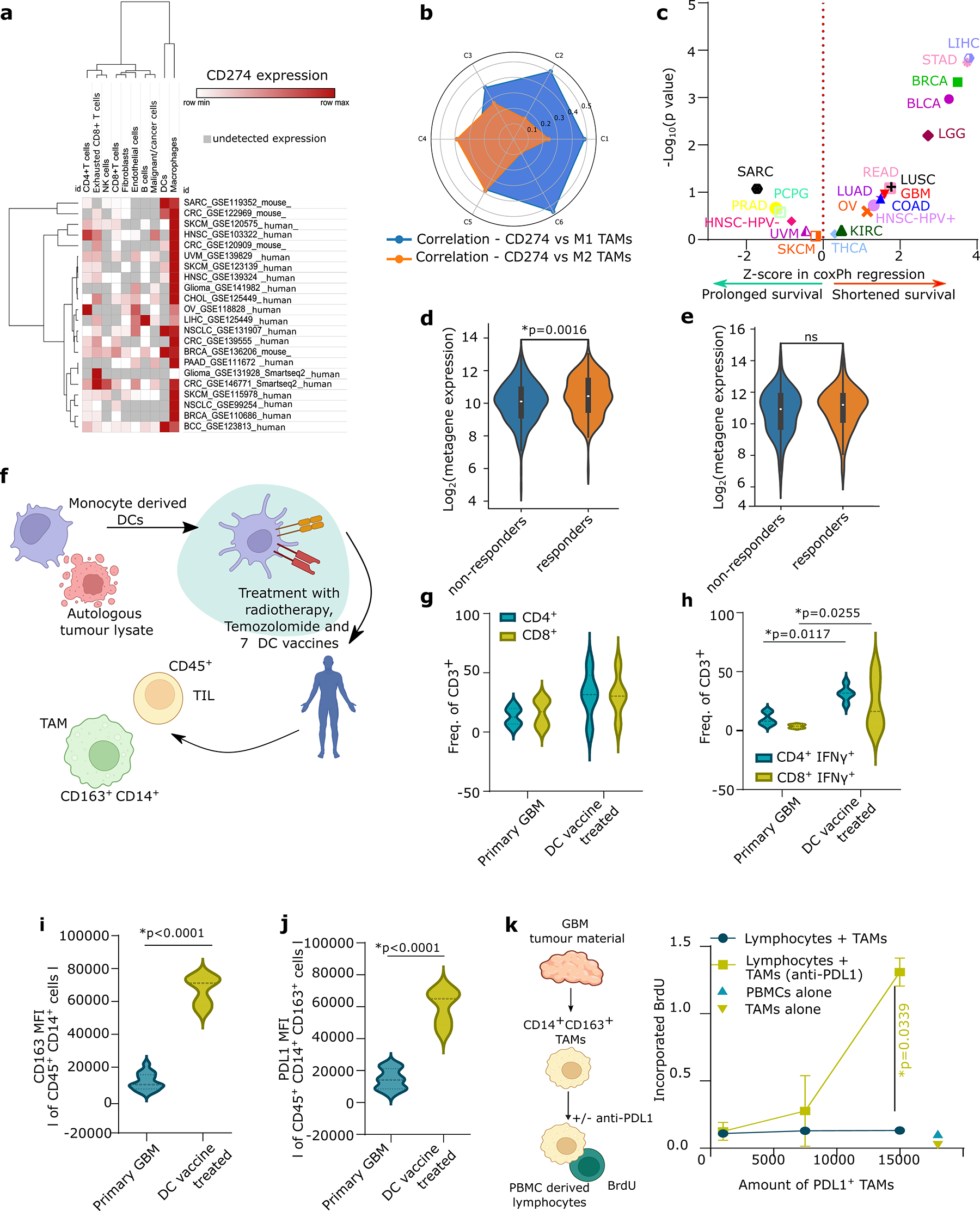
PDL1 and tumour associated macrophages (TAMs) are negative prognostic markers in human cancers and are also mobilized by DC vaccines in a prototypical human T cell-depleted tumour. (A) Heatmap representation of the expression of *CD274*/*cd274* (PDL1) in different immune, cancer or stromal cells across 22 single-cell cohorts covering 13 cancer types (n= 287 patients). (B) Radar plot of the correlation between *CD274* gene expression and M1 or M2 macrophage fraction in across TCGA cancer types (C1, n=1313; C2, n=1210, C3, n=688; C4, n=222, C5, n=2; C6, n=111). (C) Z-scores associated with CoxPH regression of *CD274*^HIGH^macrophages^HIGH^ subgrouping in different cancer types, correcting for age, gender, tumour-stage (Bladder urothelial carcinoma BLCA; n=408, breast invasive carcinoma BRCA; n=1100, colon adenocarcinoma COAD; n=458, glioblastoma GBM; n=153, head and neck squamous cell carcinoma-human papillomavirus^-^ HNSC-HPV^-^; n=422, head and neck squamous cell carcinoma-human papillomavirus^+^ HNSC-HPV^+^; n=98, kidney chromophobe KICH; n=66, kidney renal clear cell carcinoma KIRC; n=533, kidney renal papillary cell carcinoma KIRP; n=290, low grade glioma LGG; n=516, liver hepatocellular carcinoma LIHC; n=371, lung adenocarcinoma LUAD; n=515, lung squamous cell carcinoma LUSC; n=501, ovarian serous cystadenocarcinoma OV; n=303, pancreatic adenocarcinoma PAAD; n=179, pheochromocytoma PCG; n=181, prostate adenocarcinoma PRAD; 498, rectum adenocarcinoma READ; n=166, sarcoma SARC; n=260, skin cutaneous melanoma SKCM; n=471, stomach adenocarcinoma STAD; n=415, thyroid carcinoma THCA; n=509, uveal melanoma UVM; n=80, Mantel-Cox test) (D-E) Violin plots of Log2(metagene expression) of CD274, CD163, CD14 and CD68. (D) Responders versus non-responders to anti-PDL1 therapy (atezolizumab or durvalumab). (responders; n=185, non-responders; n=269, of which ureter/renal pelvis cancer; n=4, urothelial cancer; n=345, bladder cancer; n=31, esophageal cancer; n=72, renal cell carcinoma; n=2; Mann-Whitney U test) (E) Responders versus non-responders to anti-PD1 therapy (nivolumab or pembrolizumab). (responders; n=183, non-responders; n=323, of which lung cancer; n=19, glioblastoma; n=19, ureter/renal pelvis cancer; n=7, gastric cancer; n=45, colorectal cancer; n=5, melanoma; n=415, bladder cancer; n=59, hepatocellular carcinoma; n=22, breast cancer; n=14, renal cell carcinoma; n=31, head & neck cancer n=110; Mann-Whitney U test) (F-J) Flow cytometry analysis of the CD45^+^ fraction of primary and DC vaccinated glioblastoma patients included in clinical trial NCT03395587. Tumour material was obtained at the day of resection at first diagnosis (primary) or at recurrence after dendritic cell vaccination. (F) Overview of clinical trial NCT03395587. (G) Frequency of CD4^+^ and CD8^+^ of CD3^+^ cells. (H) Frequency of IFNγ^+^ of CD4^+^ or CD8^+^ CD3^+^T cells. (I) Mean fluorescent intensity of CD163 on CD14^+^ cells. (J) Mean fluorescent intensity of PDL1 on CD14^+^ cells. (G-H) primary; n=6, progress vaccine; n=5, two-way ANOVA, Bonferroni’s multiple comparison) (I-J) primary; n=15, recurrent DC vaccine; n=5, two-tailed student’s t test (K) Bromodeoxyuridine incorporation in cocultures of PBMC-derived lymphocytes (CD14 depleted PBMC) and TAMs obtained from primary glioblastoma samples with and without anti-PDL1 blocking. (n=3; area under curve-driven two-tailed paired t test). See also figure S16

Next, we wanted to understand if *CD274* indeed dominantly co-associated with M2-like TAMs in T cell-depleted human tumours. For this, we interrogated *CD274*’s correlation to M1 vs. M2-like macrophage fractions across TCGA’s pan-cancer immune-landscape classes. Interestingly, the more immunogenic C1/C2/C6-tumours showed a higher correlation between *CD274* and M1-macrophages (**Fig.7B**). Strikingly, in line with TC1 data, the T cell-depleted C4/C5-tumours displayed a higher correlation between *CD274* and M2-like macrophages (**Fig.7B**). Finally, in terms of survival impact, a large TCGA-based pan-cancer multi-variate prognostic analysis (spanning 19 cancer-types and 8493 patients), emphasized that *CD274*^HIGH^ and macrophage-fractions^HIGH^ co-enrichment associated with increased hazard ratio (i.e., shorter overall survival) in majority of cancer types (**Fig.7C**).

Finally, we saw that the M2-like PDL1^+^TAMs in TC1-tumours created a specific immunotherapy-resistant niche that was partially susceptible to only anti-PDL1 ICB, but not anti-PD1 ICB. Hence, we wondered if also in a human multi-cancer setting, M2-like PDL1^+^TAM’s signature exclusively predicted positive clinical responses to only anti-PDL1 ICBs. Hence, we analysed if a simple M2-like PDL1^+^TAM-metagene (detected in pre-treatment tumour transcriptome, and composed of *CD274, CD68, CD163, CD14*) can distinguish clinical responders from non-responders, in a large-scale immunotherapy clinical cohort integrating treatments with either anti-PDL1 ICB (454 patients, 5 cancer-types) or anti-PD1 ICB (761 patients, 11 cancer-types). Interestingly, PDL1^+^TAM-metagene was more significantly expressed in patients responding to anti-PDL1 ICB (**Fig.7D**) than anti-PD1 ICB (**Fig.7E**). This emphasized PDL1 and TAMs co-association as a pan-cancer negative prognostic niche such that PDL1^+^TAM-metagene predicted positive clinical responses to only anti-PDL1 ICB. This also confirmed that our major pre-clinical conclusions derived from TC1-tumour model had multi-cancer (predictive) clinical implications.

### DC vaccines mobilize lymphocytes-suppressive PDL1^+^TAMs in a prototypical human T cell-depleted tumour

Above preclinical phenotypes were expected to have a pan-cancer (biomarker-like) ‘predictive’ implication and hence be operational in any prototypical human T cell-depleted tumour. Thus, the observation of DC vaccines-driven facilitation of T cell-suppressive PDL1^+^TAMs, needed functional validation in suitable DC vaccinated patients. A cancer-type distribution analyses for C4/C5-tumours found that many of the T cell-depleted tumours were glioblastoma (GBM) or gliomas (LGG) (**Suppl. Fig.S16D**), which are well-established to be T cell-depleted, TAMs-enriched, and ICB resistant (Naulaerts et al., 2021; Pombo Antunes et al., 2021; Woroniecka et al., 2018).

Hence for validation, we accessed samples from the GlioVax clinical trial (Rapp et al., 2018) (for patient clinical details, see **Supplementary Table 3**). GlioVax is a randomized phase II clinical trial involving newly diagnosed GBM-patients treated with tumour lysate-loaded DC vaccines combined with radio/chemo-therapy (**Fig.7F**). Further details of the trial/DC-vaccines are described elsewhere (Rapp et al., 2018). Whereas DC vaccines indeed mobilized increased CD4^+^/CD8^+^T cells (**Fig.7G**) and effector IFNγ^+^CD4^+^/CD8^+^T cells (**Fig.7H; Fig.S17A**) within GBM-tissue of vaccinated patients (compared to unvaccinated newly-diagnosed GBM patients) these patterns weren’t significant for CD4^+^/CD8^+^T cell-infiltrates. Interestingly, DC vaccines also mobilized a significant enrichment of M2-like CD163^+^CD14^+^CD45^+^ TAMs (**Fig.7I**) as well as PDL1^+^CD163^+^CD14^+^CD45^+^TAMs (**Fig.7J**) within GBM tissue of vaccinated patients. To assess if GBM-infiltrating PDL1^+^TAMs indeed suppressed proliferation of lymphocytes in a PDL1-dependent fashion, we isolated GBM-infiltrating human PDL1^+^TAMs and co-cultured them (with increasing PDL1^+^TAM amounts) with human allogeneic PBMC-derived lymphocytes in presence or absence of anti-PDL1 ICB (**Fig.7K**). Remarkably, blockade of PDL1 caused massive spike in lymphocytes proliferation, in proportion to gradually increasing amounts of PDL1^+^TAMs (**Fig.7K**). Of note, human GBM-infiltrating M2-like TAMs expressed significantly higher PDL1 than CSF1R (**Suppl. Fig.S17B**). Above observations provided strikingly consistent clinical validation of our preclinical results. Taken altogether this emphasized that DC vaccines also facilitate lymphocytes-suppressive, M2-like, PDL1^+^TAMs in human T cell-depleted tumours.

## DISCUSSION

Preclinical evidence suggests that future DC vaccination studies need to focus on improving DCs’ antigen presentation, DCs’ LN-homing, DC maturation, and combination with other immunotherapies (Goswami et al., 2022). But it is necessary to also understand the reasons behind failures of previous DC vaccines, to rationally guide DC vaccine design and improvement, and to avoid repeating the same mistakes with future vaccines (Laureano et al., 2022).

We found that the biggest limitation with human DC vaccines was not overall maturation *per se* but their contradictory maturation-trajectories. In fact, once the most immunogenic DC vaccine maturation-trajectory was distinguished (type I IFN/ISG response-driven) from the less immunogenic ones (macrophage-like or mregDC-like), it immediately associated with high antigen-directed immunity and favourable tumour responses in patients (**Fig.8**). Eventually we designed a preclinical DCvax-IT, informed by human DC vaccine’s type I IFN/ISG-response trajectory. Although some studies have investigated the single DC vaccine transcriptome (Castiello et al., 2017; Maurer et al., 2020), they have rarely used these data to drive the creation of an optimized pre-clinical DC vaccine. Nevertheless, while our (pre-)clinical focus was limited to moDCs, our results are potentially applicable to cDC1/cDC2/pDC-based vaccines. For example, type I IFN/ISG-response is also crucial for anti-cancer immunity driven by pDC-vaccines in melanoma patients (van Beek et al., 2020). Similarly, the clinically unfavourable mregDC-phenotype is more enriched in cancer antigens-exposed cDC1/cDC2, than moDCs (Maier et al., 2020).

**Figure 8.**
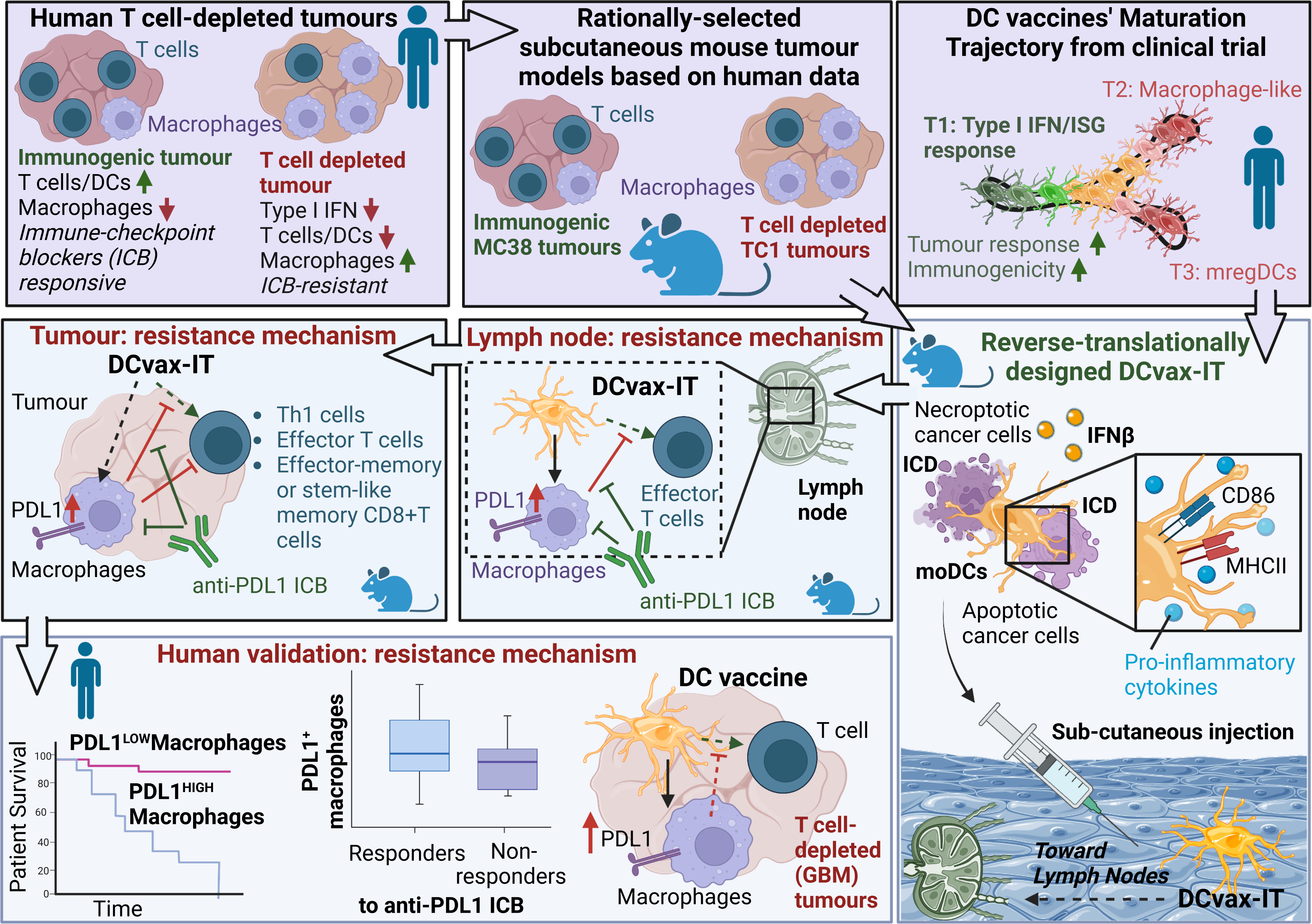
A schematic overview of our study’s main conclusions. Please see discussion for further details.

Despite above significant optimization efforts and a tailored approach toward targeting T cell-depleted TC1-tumours, DCvax-IT alone were not sufficient. In fact, contrary to expectations, DCvax-IT established interactions with LAMs and facilitated creation of PDL1^+^LAMs via their type I IFN/ISG response-signalling, rather than facilitating effector T cell-activity. In parallel, DCvax-IT also fuelled already pre-existing T cell-suppressive, M2-like, PDL1^+^TAMs within the TC1-tumours possibly via IFNγ-producing T cells and/or tumoural infiltration of PDL1^+^LAMs (**Fig.8**). This created a strong LN-to-tumour PDL1-driven LAM/TAM-based immunosuppressive circuit and was a key player in DCvax-IT immuno-resistance. It formed a barrier to effector T cell-activity in LNs and to Th1 cells or effector/stem-like memory CD8^+^T cells in the tumours. In fact, it required a DCvax-IT plus anti-PDL1 ICB combination to deplete PDL1^+^LAMs in LNs and PDL1^+^TAMs in the tumours, to unleash LN-to-tumour anti-cancer T cell-activity (**Fig.8**). This combination significantly suppressed TC1-tumour growth in a TAMs and CD8^+^T cells-dependent fashion. Using *Ifnar1*^-/-^DCvax-IT or *Ccr7*^-/-^DCvax-IT, we proved that IFNβ-driven immunogenicity and intra-LN homing of DCvax-IT were crucial for sustaining above LN-to-tumour PDL1^+^LAM/TAM-based immunosuppressive circuit and setting up the anti-tumour synergism between DCvax-IT plus anti-PDL1 ICB.

These results have major implications for changing the fundamental outlook of how anti-cancer moDC vaccines work. Currently the literature consensus unanimously agrees that LNs serve solely as the sites of efficacious T cell-priming for cancer antigens by the DCs, such that the less than proficient effector activation of LN-associated T cells after DC vaccination can be ascribed to the low immunogenicity of moDCs. In this sense, LNs have not been suspected to be sites for creation of T cell-suppressive LAMs by DC vaccines to the best of our knowledge.

Thus, our observations are highly original because a DC vaccines-driven T cell-suppressive pathway fuelled by PDL1^+^LAMs, making a circuit with PDL1^+^TAMs, that counters DC vaccines’ own ability to induce anti-tumour immunity has far-reaching clinical implications. Moreover, they immediately open novel combinatorial immunotherapy opportunities e.g., PDL1^+^LAM/TAM targeting via anti-PDL1 ICB (Laureano et al., 2022). Most proposals for immunotherapy combinations with DC vaccines revolve around anti-CSF1R ICB based targeting of TAMs or general T cell-targeting via anti-PD1 ICB (Perez and De Palma, 2019; Sprooten et al., 2019b; Wculek et al., 2020). However, such interventions are not tailored for DC vaccines. In fact, we clearly show that both anti-PD1 and anti-CSF1R ICBs did not synergize with DCvax-IT in regressing TC1-tumours. Although anti-PDL1 ICB is indeed already part of the clinically approved immunotherapy toolkit (Chae et al., 2018; Wei et al., 2018) yet the proposal to combine DC vaccines with anti-PDL1 ICBs is still novel because almost all current clinical trials integrating ICBs with DC vaccines are prioritising anti-PD1 ICBs (Laureano et al., 2022).

Finally, as an instance of mice-to-human translational validation for DC vaccines, we showed that DC vaccines also potentiated lymphocytes-suppressive, M2-like, PDL1^+^TAMs in patients with GBM, a prototypical T cell-depleted tumour. This, along with the data that TAMs and PDL1 co-association is negative prognostic and high PDL1^+^TAMs-signature predicts positive patient responses to anti-PDL1 ICB, suggests that DC vaccines-driven PDL1^+^TAMs-enrichment could be a predictive biomarker of non-responding patients. This claim is substantiated by some recent clinical studies e.g., DC vaccines increased tumoural PDL1 in lung cancer patients (Lee et al., 2017), or showed higher efficacy in breast cancer patients with PDL1 negative tumours (Santisteban et al., 2021).

In conclusion, our study shows that reverse translational optimization of DC vaccines is necessary but not sufficient for anti-tumour efficacy. Because the DC vaccines paradoxically induce a self-inhibitory LN-to-tumour circuit of T cell-suppressive PDL1^+^LAMs/TAMs that form a barrier to efficacious activation of T cell-immunity in the context of T cell-depleted, TAMs-enriched, tumours. This creates a new DC vaccines-driven immuno-resistant niche that opens novel avenues for combinatorial immunotherapy.

## MATERIALS & METHODS

Detailed materials and methods, as well as an overview of clinical or statistical approaches are discussed in detail within the **‘Supplementary Methods’** document going with this manuscript.

## AUTHOR CONTRIBUTIONS

JS was the lead researcher on this project and performed most of the experimental work, co-ordinated & managed the overall research as well as data handling efforts and created the figures. SN and DMB helped closely with bioinformatics analyses and critical discussion on biostatistics. IV, JG, RSL and AC performed several crucial experiments, helped in protocol standardizations and extensive manuscript/figure formatting. AD, MK, MS, MR, CKT and RVS collected relevant GBM-patient samples and performed the flow cytometric or BrdU experiments as well as associated data analyses. PL, LZ, OK and GK helped in generation of CRISPR/Cas9-modified TC1 cancer cell lines. LB helped with in vivo experiments involving murine antibodies blocking PD1, CTLA4, PD-L1. SS helped with PCR-analyses and critical assessment of flow cytometry data. SDV, JB and ST helped with critical data interpretation, provided expert conceptual or data-analyses views, and/or helped with manuscript writing. ADG and RVS supplied senior supervision for respective research tasks, took part in data analyses & representation decision-making, and critical revision of the manuscript. ADG was the principal investigator of the project and coordinated the overall project, (co-)supervised the *in silico*, *in vitro* or *ex vivo* research designs, as well as wrote the manuscript (and its revisions).

## COMPETING INTERESTS

Abhishek D Garg received consulting/advisory/lecture honoraria from Boehringer Ingelheim (Germany), Miltenyi Biotec (Germany), Novigenix (Switzerland), and IsoPlexis (USA). Oliver Kepp is a cofounder of Samsara Therapeutics.

## Supporting information

Key Resources

## ACKNOWLEDGEMENTS

We are immensely thankful to Prof. Roos Vandenbroucke (VIB-UGent) for providing us with the *Ifnar1*^-/-^C57BL/6 mice as well as Dr. Andrew Brown and Prof. Bart Lambrechts (VIB-UGent, Belgium) for providing us with the *Ccr7*^-/-^C57BL/6 mice. We acknowledge Prof. Patrizia Agostinis, Prof. Geert Bultynck, and Prof. Peter Vangheluwe (KU Leuven, Belgium) for supporting our research by sharing lab space and/or cell culture facilities. We would also like to thank Prof. Stefan Krautwald (Germany), Prof. Steven Verhelst (KU Leuven), Prof. An Coosemans (KU Leuven) and Prof. James Murphy (Australia) for providing us with various cell lines, tools or helping us with some experiments that unfortunately could not be accommodated in the final manuscript. We thank Prof. Pierre Coulie (UCL, Belgium), Prof. Adrian Liston (Babraham Institute, UK) and Elselien Frijlink (LUMC, The Netherlands) for attentively reading the manuscript and supplying valuable suggestions and feedback. This study is supported by Research Foundation Flanders (FWO) (Fundamental Research Grant, G0B4620N to ADG; Excellence of Science/EOS grant, 30837538, for ‘DECODE’ consortium, for ADG, JB), KU Leuven (C1 grant, C14/19/098; C3 grant, C3/21/037; and POR award funds, POR/16/040 to ADG), Kom op Tegen Kanker (KOTK/2018/11509/1 to SDV, ADG; and KOTK/2019/11955/1 to ADG) and VLIR-UOS (iBOF grant, iBOF/21/048, for ‘MIMICRY’ consortium to ADG and ST). IV is supported by FWO-SB PhD Fellowship (1S06821N). JS is funded by Kom op tegen Kanker (Stand up to Cancer), the Flemish cancer society via Emmanuel van der Schueren (EvDS) PhD fellowship (projectID: 12699). DMB is supported by KU Leuven’s Postdoctoral mandate grant (PDMT1/21/032), and the Belgian Federation against Cancer grant numbers 2018-127 and 2016-133 and by a grant from Fondation Roi-Baudouin to ST. ST is further supported by a Senior Clinical Investigator award of FWO. MCS, MR and RVS were supported by a grant from the Federal Ministry of Education and Research (BMBF; grant #01KG1242).

## Notes

### Summary of Updates

One of the co-author names was missing by mistake this has been corrected now.

