## Supplementary Figures S1-S17 for "A lymph node-to-tumour PDL1^+^macrophage circuit antagonizes dendritic cell immunotherapy"

### A lymph node-to-tumour PDL1$^+$macrophage circuit driven by dendritic cell immunotherapy inhibits anti-tumour immunity

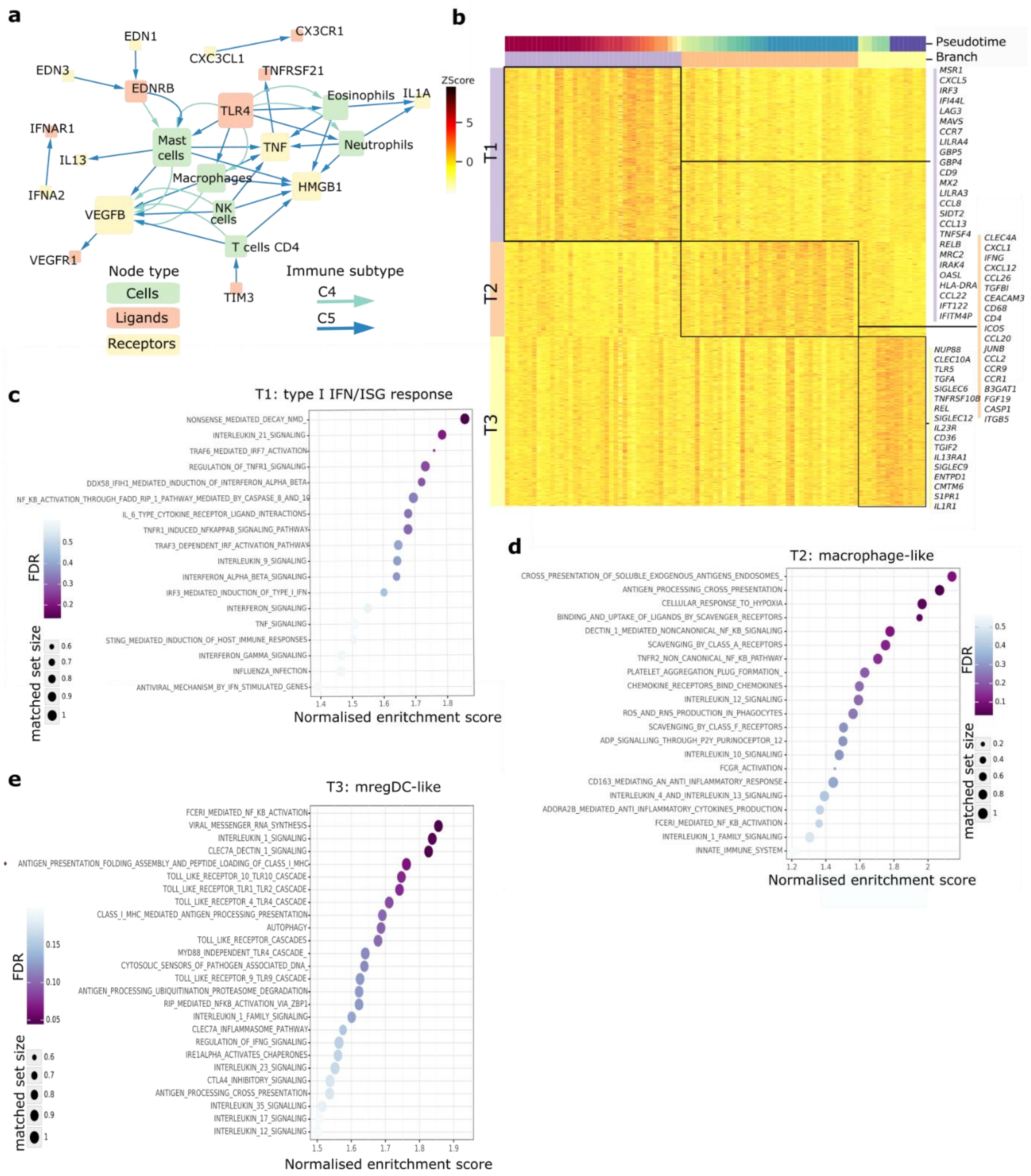


**Figure S1.**

(A) Discordance or non-correlative extracellular network analysis of the C4/C5 (T cell hostile) human tumours annotated in TCGA pan-cancer dataset (n=224). Of note, in this network, a connection or "pairing" between two nodes (i.e., a pair of two genes or a pair of a gene and an immune cell) indicates that those two nodes are non-correlated to each other thereby indicating discordance of pathway. Node size reflects the number of connections the node participates in (node degree).

(B-E) STREAM DC vaccine trajectory analysis of 93 autologous DC vaccines pulsed with LPS, IFNγ and TARP peptide generated for 18 prostate adenocarcinoma cancer patients vaccinated with 5-8 vaccines.

(B) Heatmap of upregulated expression per STREAM trajectory branch

(C-E) Set enrichment of REACTOME terms per trajectory branch. Dot size corresponds to matched set size and dot color corresponds to FDR (significance: FDR<0.05).

(C) T1: type I IFN/ISG response

(D) T2: macrophage-like.

(E) T3: mature regulatory DCs.

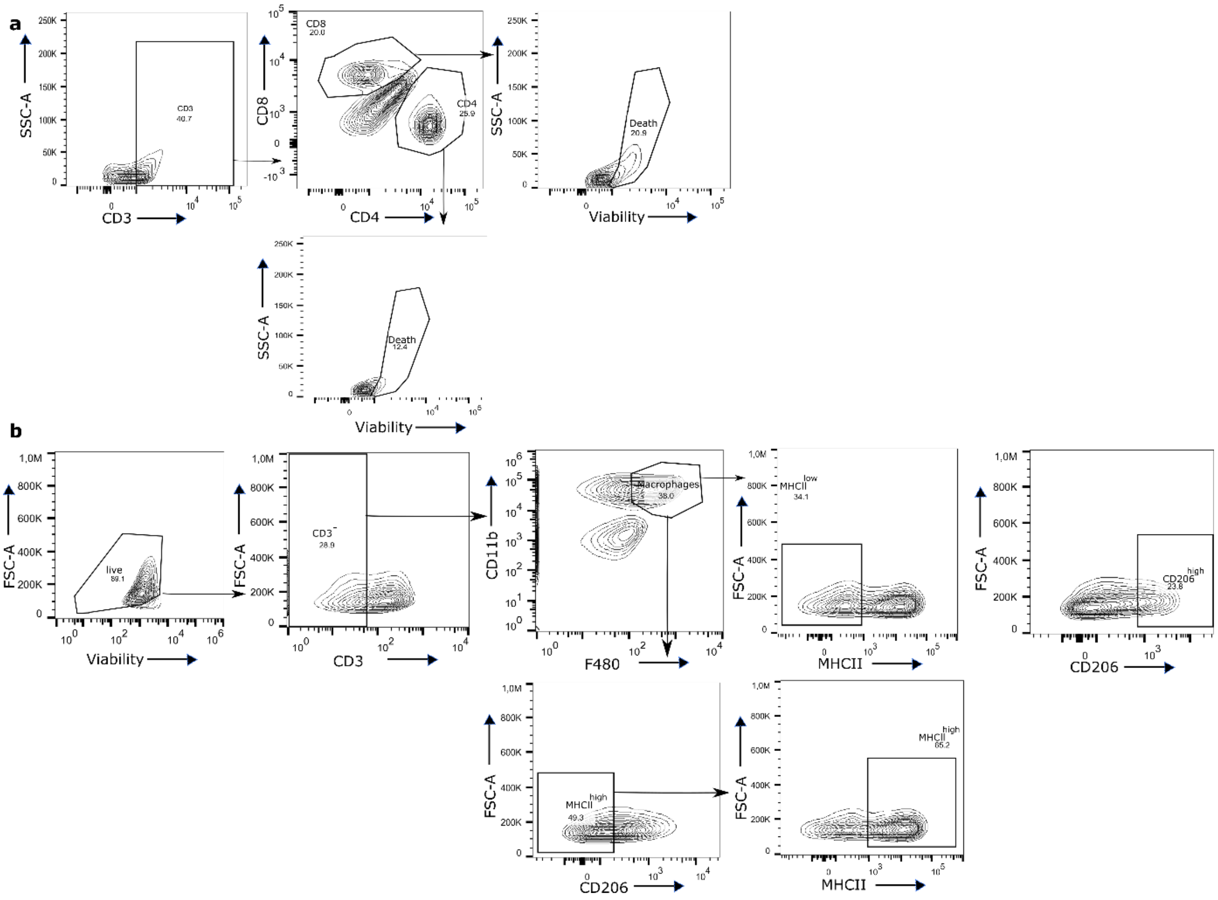


**Figure S2.**

(A) Representative gating strategy for frequency of zombie aqua$^{+}$ dying/dead cells of CD4$^{+}$ and CD8$^{+}$ T cells in the spleen of a wild-type TC1 tumor bearing mouse.

(B) Representative gating strategy for M1 (MHC-II$^{HIGH}$CD206$^{LOW}$) and M2 (MHC-II$^{LOW}$CD206$^{HIGH}$) macrophages gated for CD45$^{+}$ cells derived from wild-type TC1 tumors isolated on day 23 after wild-type TC1 cell injection.

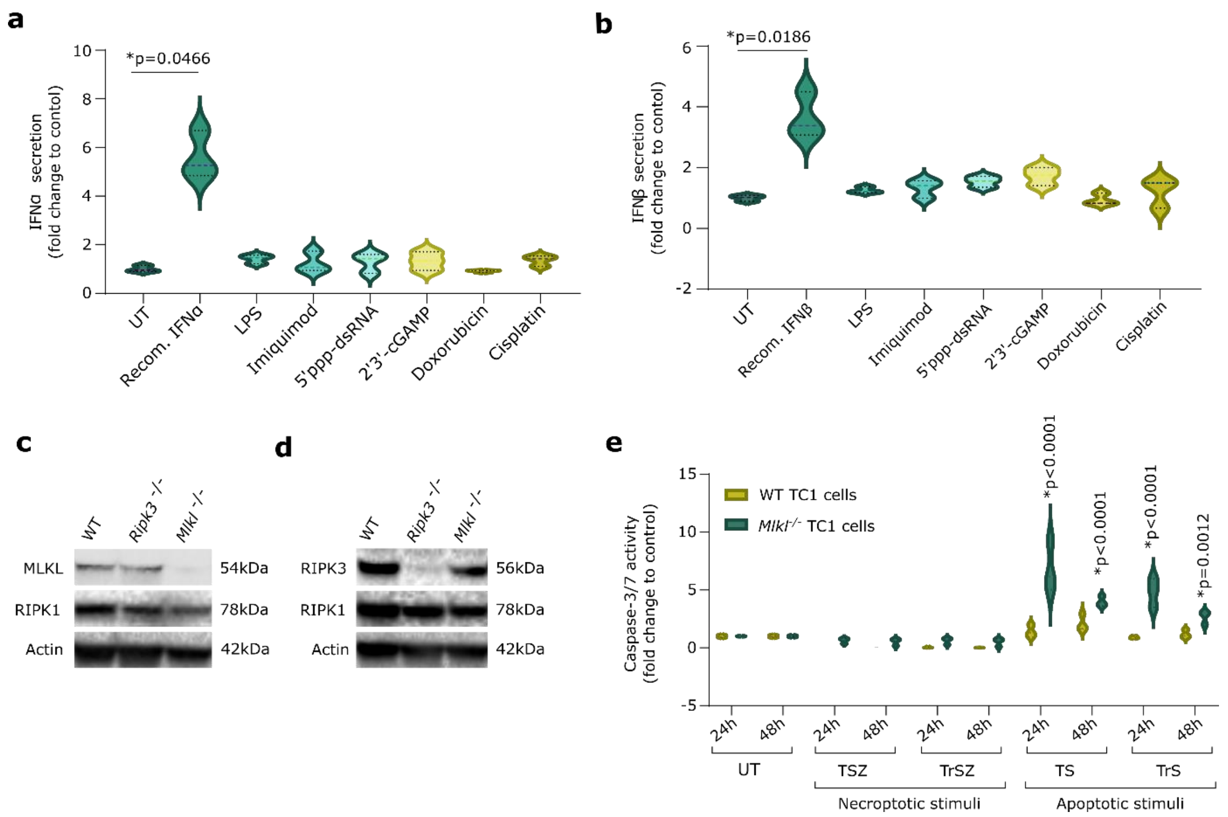


**Fig S3.**

(A-B) Secretion of interferon by wild-type TC1 stimulated with different stimuli for 24h. (A) for IFNα. (B)IFNβ. P-values depict comparison to untreated. (n=3; one-way ANOVA, Kruskal-Wallis test).

(C-D) Western blot of wild-type, *Ripk3*$^{-/-}$ and *MLKL*$^{-/-}$ TC1 for MLKL, RIPK1, RIPK3 and Actin.

(E) Caspase 3/7 activity of wild-type and *Mlkl*$^{-/-}$ TC1 stimulated with necroptotic and apoptotic stimuli. P-values depict comparison to 24h or 48h untreated. (n=3; 2way ANOVA, Dunnett's multiple comparisons test).

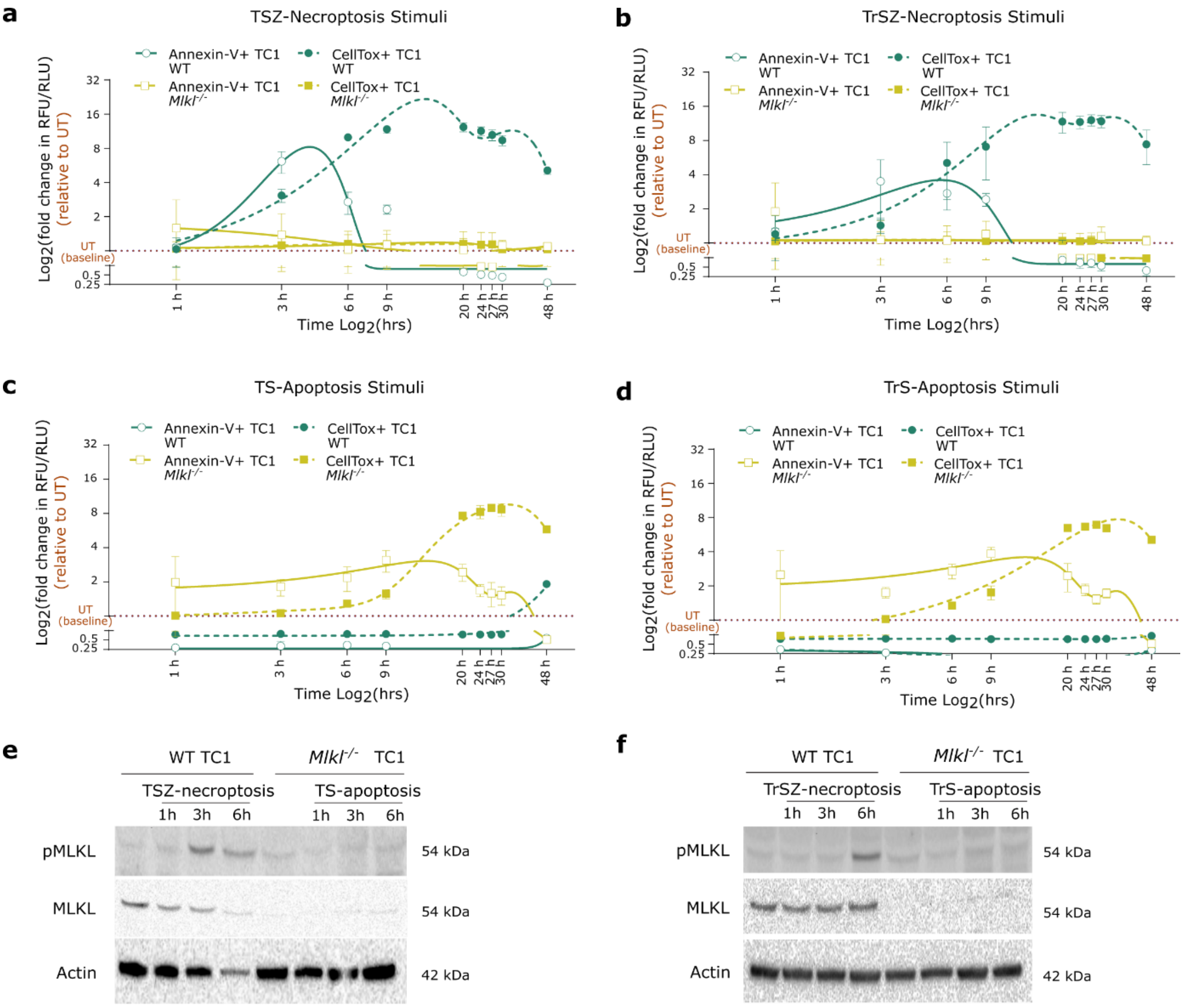


**Figure S4.**

(A-D) Kinetics of Annexin-V and CellTox staining at different timepoints for dying wild-type and *MLKL-/-* TC1 cells treated with (A) TSZ-necroptotic stimuli (B) TrSZ-necroptotic stimuli (C) TS-apoptotic stimuli (D) TrS-apoptotic stimuli TrS**.** (n=3).

(E-F) Western blot for phosphorylated MLKL (pMLKL), MLKL and Actin of wild-type and *MLKL-/-* TC1 cells stimulated with necroptotic stimuli and apoptotic stimuli.

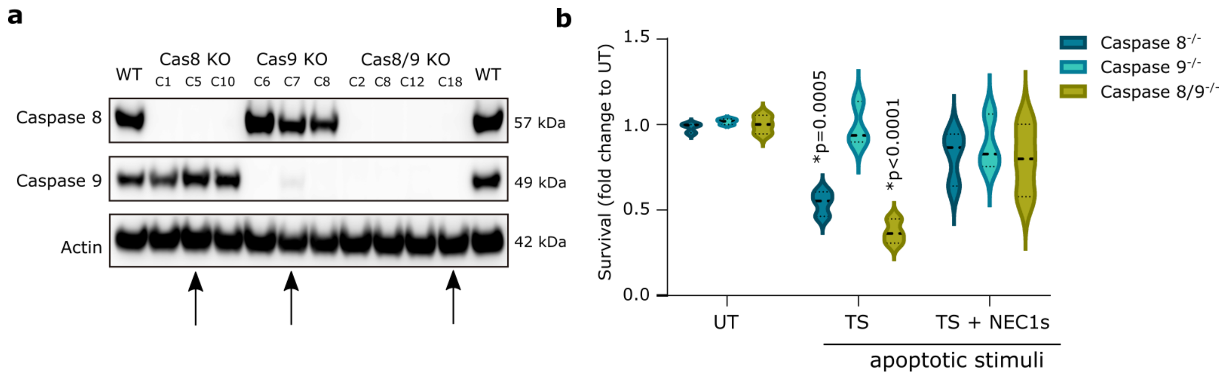


**Figure S5.**

(A) Western blot of caspase 8, caspase 9 and actin of different clones of TC1 *Caspase 8*$^{-/-}$, *Caspase 9*$^{-/-}$ and *Caspase8/9*$^{-/-}$. Arrows indicate the used clone.

(B) Survival of TC1 *Caspase 8*$^{-/-}$, *Caspase 9*$^{-/-}$ and *Caspase8/9*$^{-/-}$ stimulated with cell death cocktails. P-values depict comparison to untreated. (n=3; two-way ANOVA, Dunnett's multiple comparisons test)

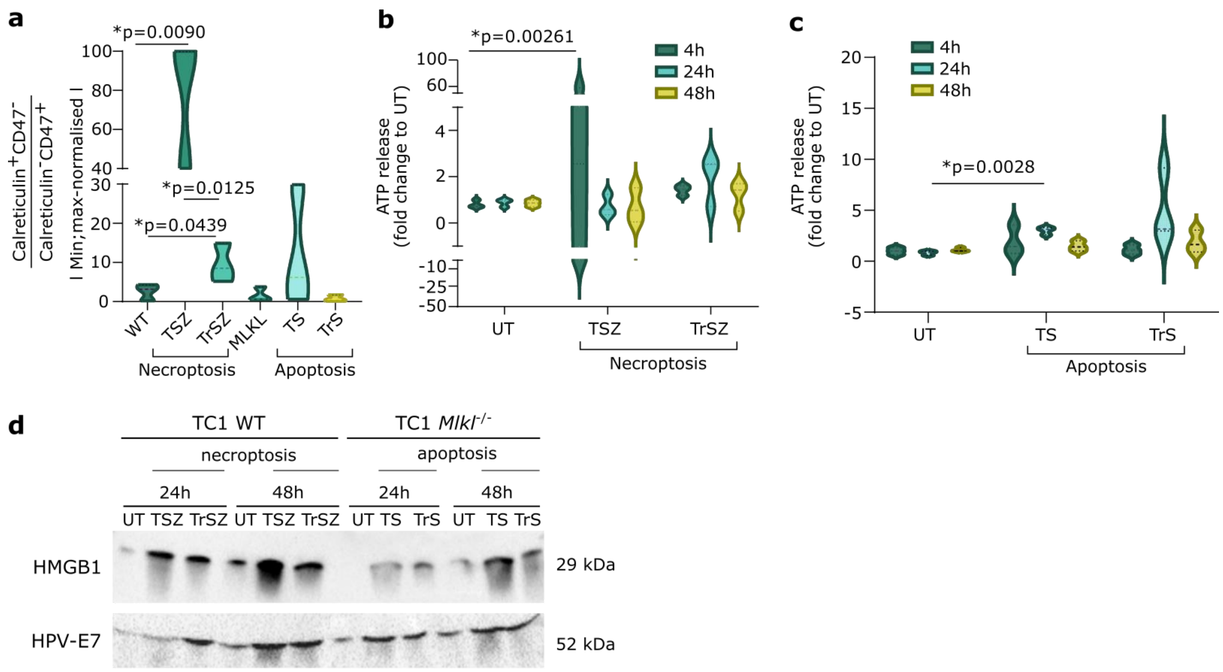


**Figure S6.**

(A) Calreticulin$^{+}$ CD47$^{-}$-to- Calreticulin$^{-}$ CD47$^{+}$ ratio of wild-type and *Mlkl*$^{-/-}$ TC1 cells treated with necroptotic and apoptotic stimuli. P-values depict comparison to untreated. (n=3; two-tailed student's t test).

(B-C) ATP release at 4h, 24h and 48h of wild-type and *Mlkl*$^{-/-}$ TC1 cells treated with necroptotic and apoptotic stimuli. P-values depict comparison to 4h, 24h or 48h untreated. (n=3; one-way ANOVA, Dunnett's multiple comparison test).

(D)Western blot for HMGB1 and HPV-E7 antigen secretion in the media of wild-type and *Mlkl*$^{-/-}$ TC1 cells treated with necroptotic and apoptotic stimuli.

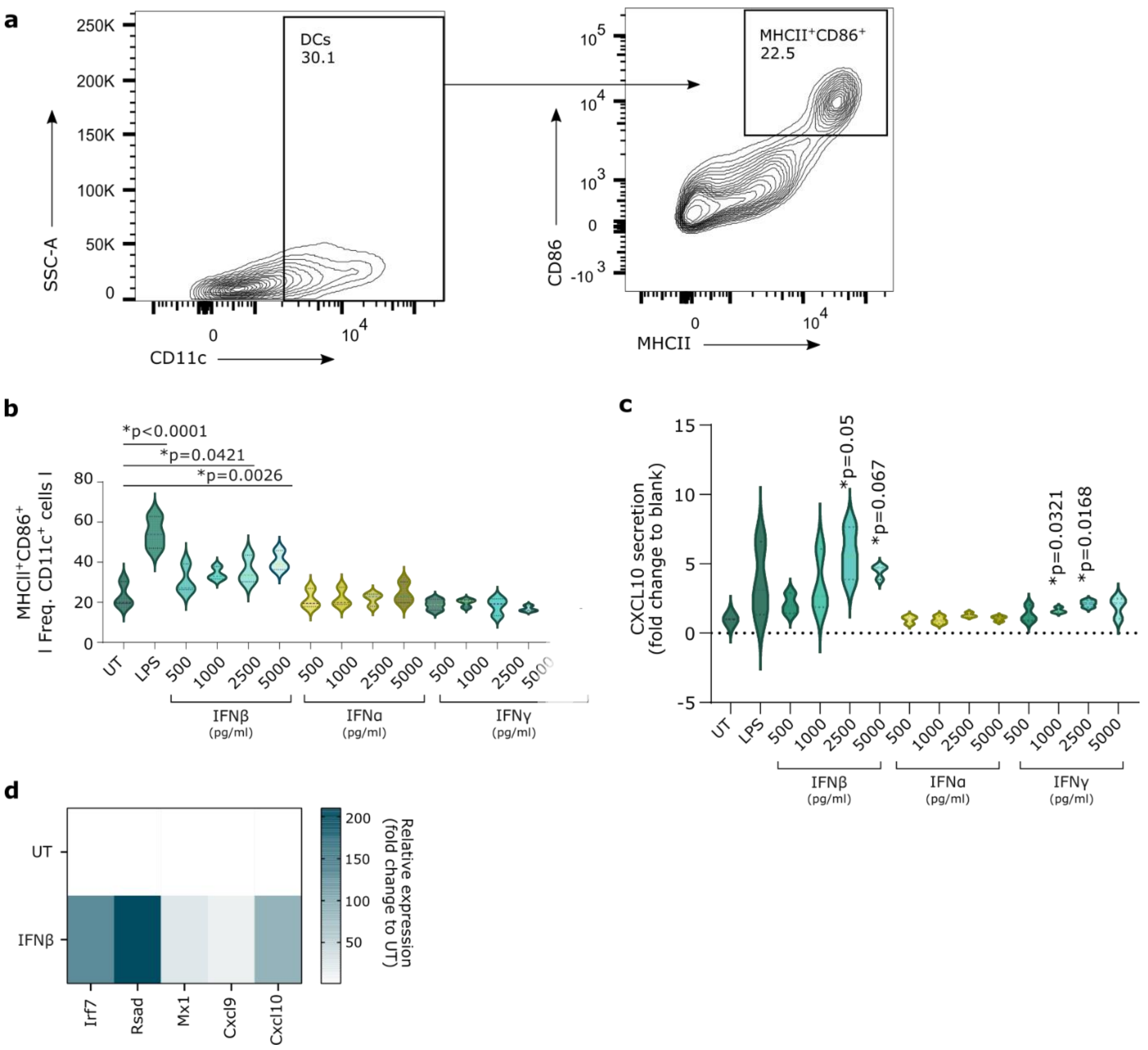


**Fig S7.**

(A) Representative gating strategy for DC maturation in DCs stimulated with LPS.

(B) MHCII$^+$CD86$^+$ of CD11c$^+$ of DCs treated with different concentrations of IFNα/β/γ.

(C) CXCL10 secretion of DCs treated with different concentrations of IFNα/β/γ.

(B-C) P-values depict comparison to untreated. (n=3; one-way ANOVA, Dunnett's multiple comparison test)

(D) Relative expression of different interferon-stimulated genes, interferon-response factor 7 (*Irf7*), Radical S-Adenosyl Methionine (*Rsad*), MX Dynamin Like GTPase 1 (*Mx1*), Chemokine (C-X-C motif) ligand 9 (*Cxcl9*), Chemokine (C-X-C motif) ligand 10 (*Cxcl10),* of which the IFN-signature is composed.

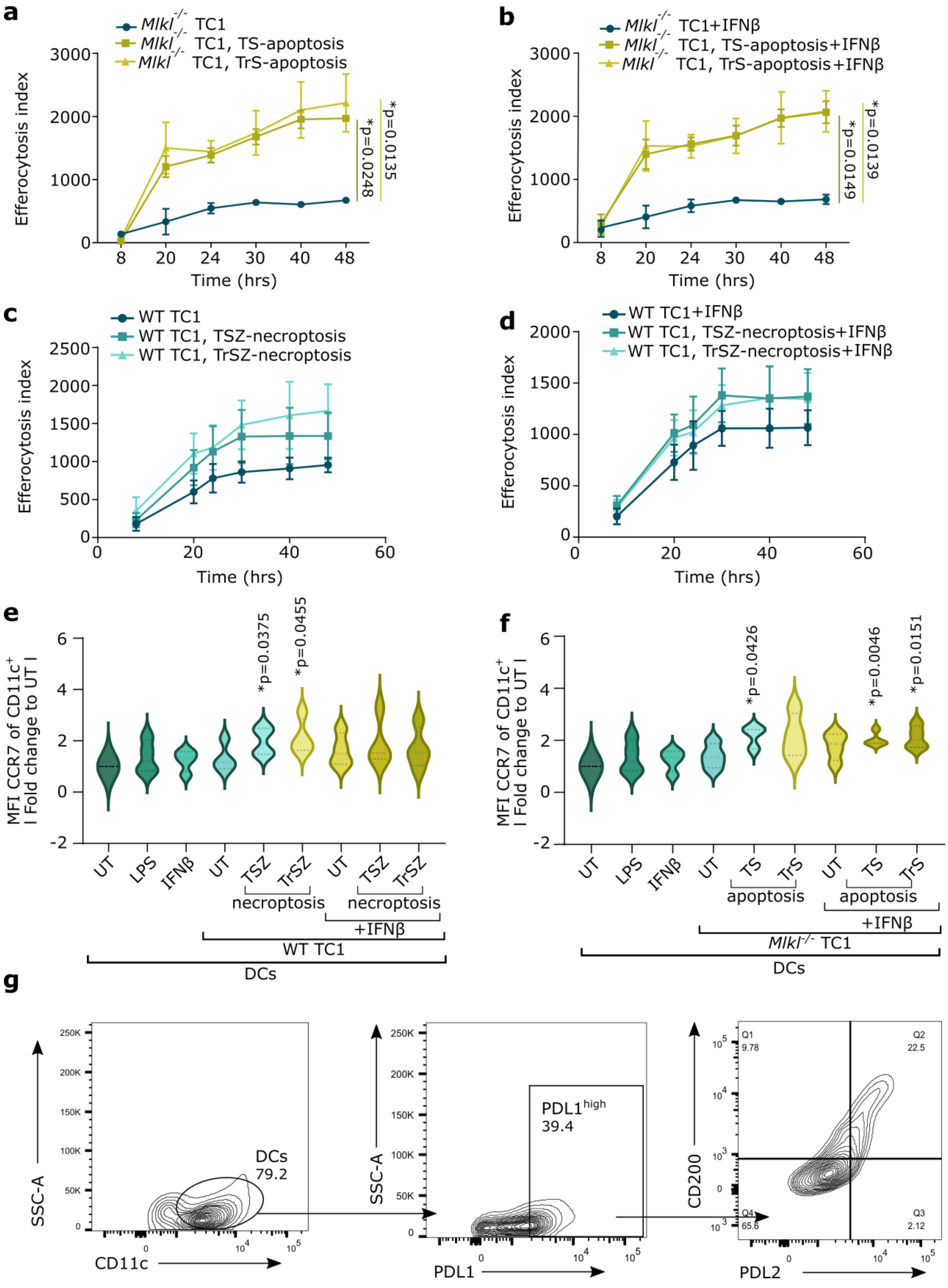


**Figure S8.**

(A-D) Efferocytosis index of DCs stimulated with pHRodo stained TC1 cancer cells calculated by the fluorescent intensity values, subtracted by the appropriate 4°C negative control.

(A-B) DCs stimulated with apoptotic *Mlkl*$^{-/-}$ TC1 cells. P-values depict comparison to untreated *Mlkl*$^{-/-}$ TC1 cells or *Mlkl*$^{-/-}$TC1 cells + IFNβ. (n=3; area under curve; one-way ANOVA, Dunnett's multiple comparison test).

(C-D) DCs stimulated with necroptotic wild-type TC1 cells. P-values depict comparison to untreated wild-type TC1 cells. (n=4; area under curve; one-way ANOVA, Dunnett's multiple comparison test).

(E-F) Fold change of mean fluorescence intensity of CCR7 of CD11c$^{+}$ cells to untreated DCs. DCs were stimulated with (E) necroptotic wild-type TC1 cells (F) apoptotic *Mlkl*$^{-/-}$ TC1 cancer cells with and without IFNβ. P-values depict comparison to untreated DCs. (n=4 (ts; n=3); one sample t-test).

(G) Representative gating strategy for mature regulatory DC gating of DCs treated with LPS.

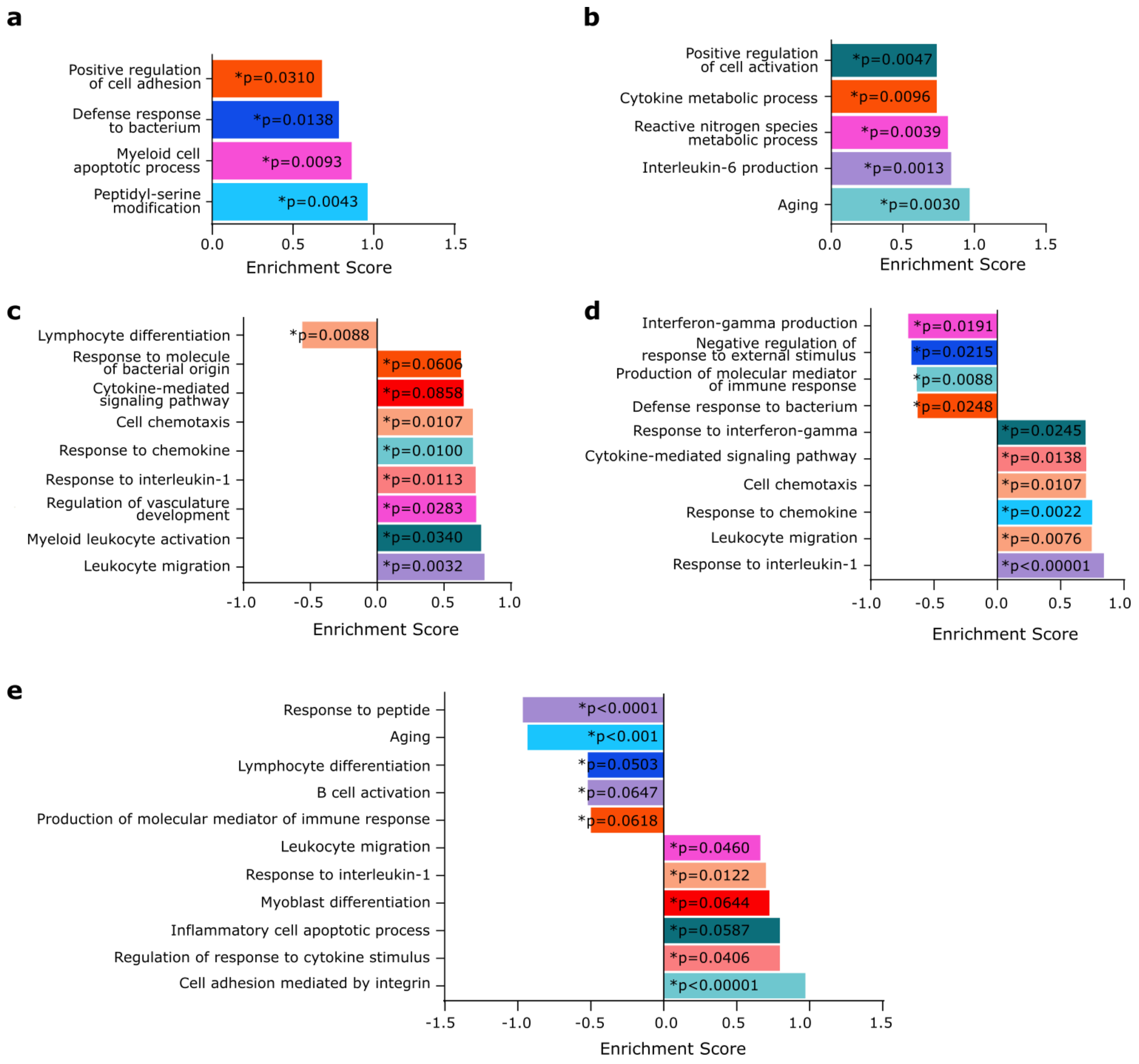


**Figure S9.**

(A-E) Set enrichment analysis of Gene Ontology Biological Process terms based on the secretome of DCs stimulated with (A) necroptotic wild-type TC1 cells, (B) apoptotic *Mlkl*$^{-/-}$ TC1 cells, (C) untreated wild-type or *Mlkl*$_{-/-}$ TC1 cells with IFNβ, (D) necroptotic wild-type TC1 cells with IFNβ or (E) apoptotic *Mlkl*$^{-/-}$ TC1 cells with IFNβ. P-values < 0.05 are marked with *.

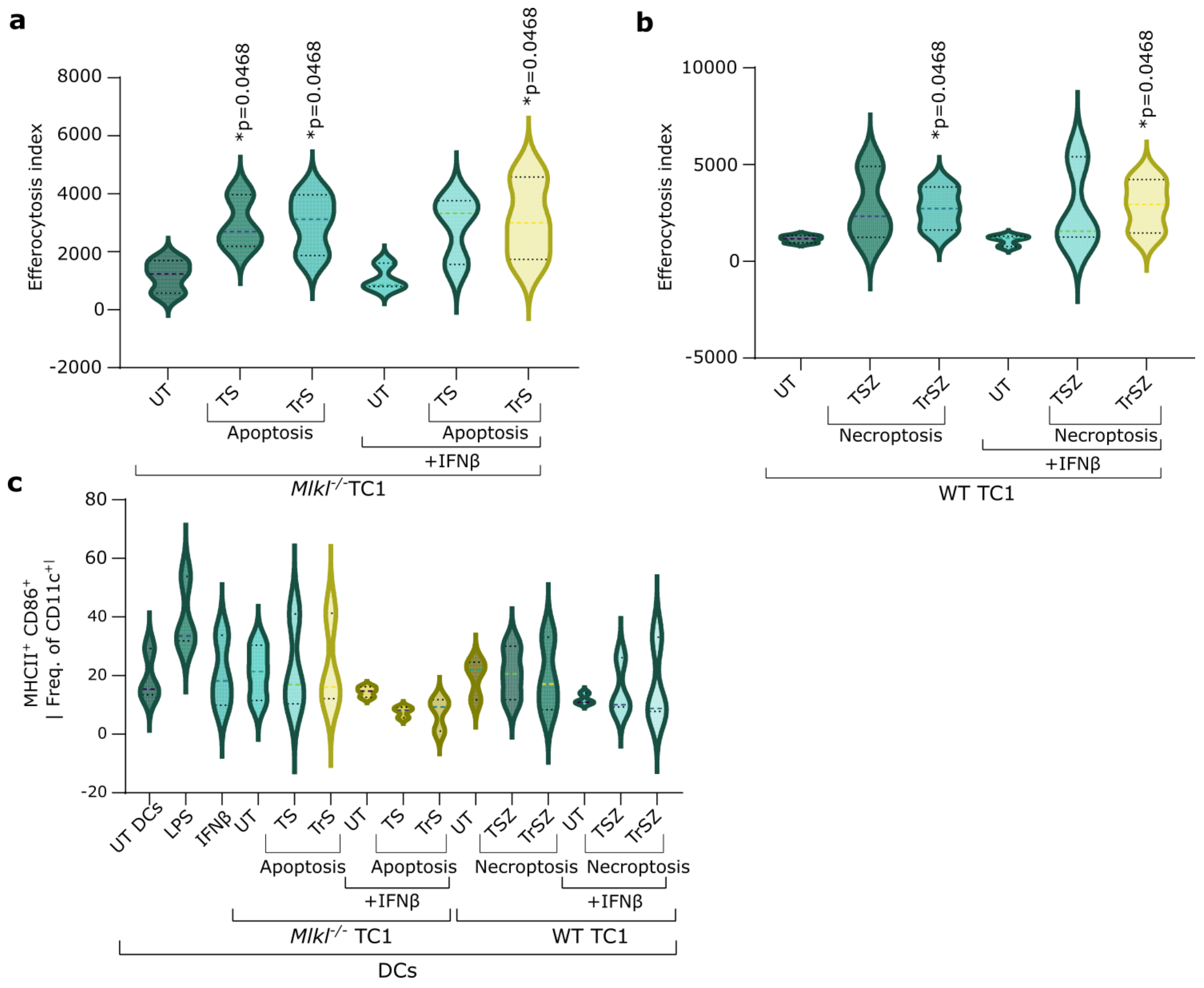


**Figure S10.**

(A-B) Efferocytosis index of *Ifnar1*$^{-/-}$ DCs stimulated with pHRodo stained (A) apoptotic *Mlkl*$^{-/-}$ TC1 cells or (B) necroptotic wild-type TC1 cells. Values were calculated by the fluorescence intensity values, subtracted by the appropriate 4°C negative control (n=3; one-way ANOVA, Kruskal-Wallis test).

(C) Maturation of DCs, assessed by MHCII$^{+}$CD86$^{+}$, untreated or treated with LPS, IFNβ or dying cancer cells with or without IFNβ (n=3; One-way ANOVA, Dunnett's multiple comparison).

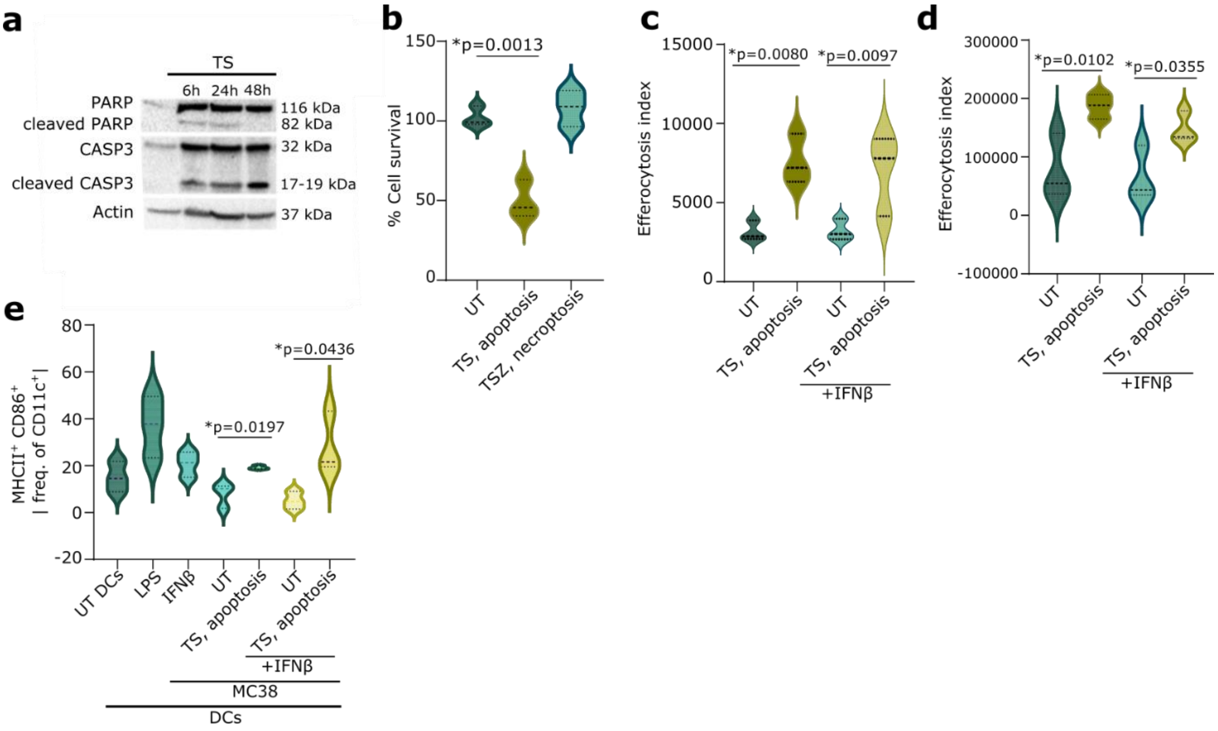


**Figure S11.**

(A) Western blot for PARP, Caspase3 and Actin of MC38 treated with apoptotic stimuli.

(B) Survival of MC38 stimulated with apoptotic (TS) or necroptotic (TSZ) stimuli measured with the MTT assay. P-values depict comparison to untreated MC38 cells. (n=3; One-way ANOVA, Dunnett's multiple comparison).

(C-D) Efferocytosis index of DCs stimulated with pHRodo stained apoptotic MC38 cells with or without IFNβ at (C) 24h (D) 48h. Values were calculated by the fluorescent intensity values, subtracted by the appropriate 4°C negative control. P-values depict comparison to untreated or IFNβ treated MC38 cells. (n=3; One-way ANOVA, FDR correction according to Benjamini, Krieger and Yekutieli method).

(E) Frequency of $MHCII^{+}CD86^{+}$ of $CD11c^{+}$ in DCs stimulated with untreated or apoptotic MC38 with or without IFNβ. P-values depict comparison to DCs with untreated or IFNβ treated MC38 cells. (n=3; two-tailed student's t test)

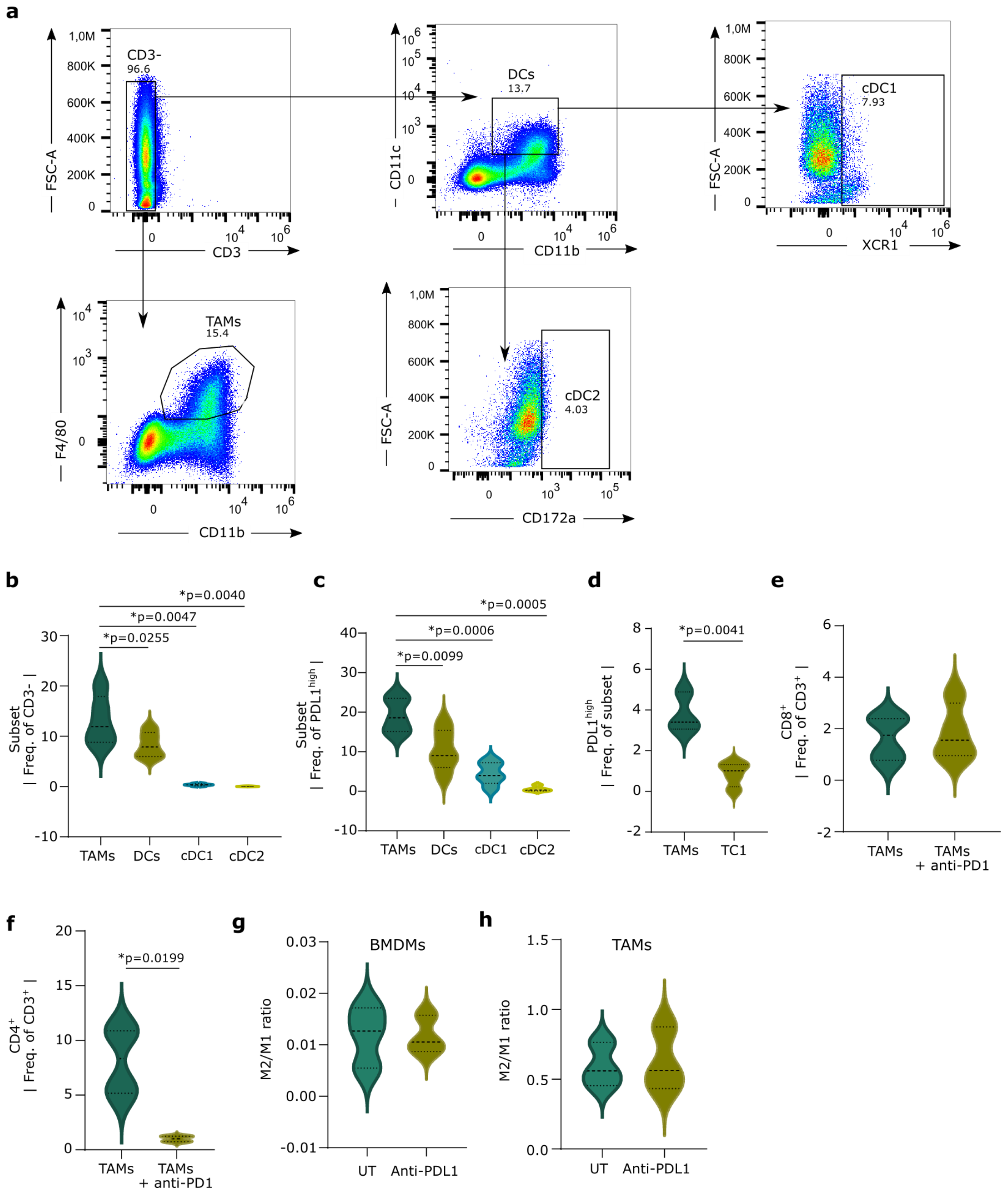


**Figure S12.**

(A) Gating strategy for Tumor associated macrophages, dendritic cells, and conventional DC1/2. Example demonstrates the gating for the CD45$^+$ population of an untreated wild-type TC1 tumor isolated on day 23 after wild-type TC1 cell injection.

(B) Frequency of subset of CD3$^-$ cells in CD45$^+$ cells isolated from wild-type TC1 tumors on day 23 after wild-type TC1 cell injection. Cell populations were assessed by CD11b$^+$ F4/80$^+$ (TAMs), CD11b$^+$CD11c$^+$

(DCs), CD11b$^+$CD11c$^+$XCR1$^+$ (cDC1), CD11b$^+$CD11c$^+$CD172a$^+$ (cDC2). P-values depict comparison to TAMs. (n=5; One-way ANOVA, FDR correction according to Benjamini, Krieger and Yekutieli method).

(C) Frequency of subset of PDL1$^+$ cells in CD45$^+$ cells isolated from wild-type TC1 tumors on day 23 after wild-type TC1 cell injection. Cell populations were assessed by CD11b$^+$ F4/80$^+$ (TAMs), CD11b$^+$CD11c$^+$ (DCs), CD11b$^+$CD11c$^+$XCR1$^+$ (cDC1), CD11b$^+$CD11c$^+$CD172a$^+$ (cDC2). P-values depict comparison to TAMs. (n=5; One-way ANOVA, FDR correction according to Benjamini, Krieger and Yekutieli method).

(D) Frequency of PDL1$^+$ cells in in vitro cultured TC1 cells or TAMS derived from wild-type TC1 tumors on day 23 after TC1 cell injection. (TAMs; n=3, TC1; n=4, two-tailed student's t test)

(E-F) Coculture experiments of T cells with wild-type TC1 tumor derived TAMs on day 23 after wild-type TC1 cell injection pretreated for 48h with or without anti-PD1. Frequency of (E) CD8$^+$ of CD3$^+$ cells (F) or CD4$^+$ of CD3$^+$ cells. (n=4, two-tailed paired t-test)

(G-H) MHCII$^{low}$ CD206$^{high}$/CD206$^{low}$ MHCII$^{high}$ (M2/M1) ratio with and without anti-PDL1 treatment for 48h of (G) bone marrow derived macrophages (BMDMs) (H) wild-type TC1 derived TAMs. (n=3; two-tailed paired t-test)

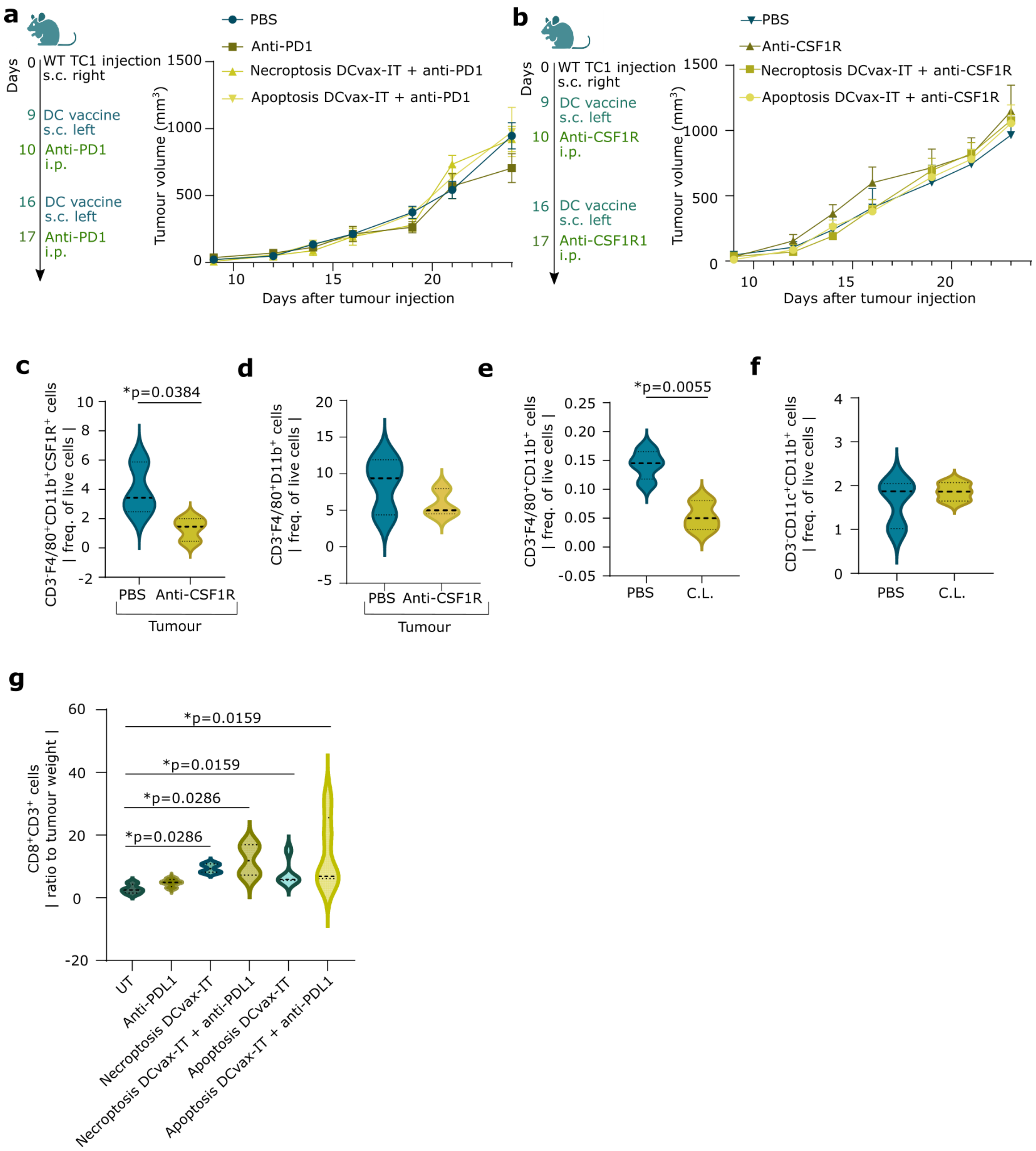


**Figure S13**

(A-B) Tumor volume curve of wild-type TC1 tumor bearing mice treated with DCvax-IT on day 9 and 16 in combination with (A) anti-PD1 on day 10 and 17 (B) anti-CSF1R on day 10 and 17. P-values depict comparison to PBS treated mice (n=4-8; area under curve, one-way ANOVA, Dunnett's multiple comparison test).

(C-D) Violin plots of the CD45$^+$ cell population of wild-type TC1 tumors isolated on day 23 after wild-type TC1 cell injection. TC1 tumors were treated with DCvax-IT on day 9 and 16 in combination with anti-CSF1R on day 10 and 17.

(C) Frequency of CD3$^-$F4/80$^+$CD11b$^+$CSF1R$^+$ of live cells (n=3; one-tailed student's t test).

(D) Frequency of CD3$^-$F4/80$^+$CD11b$^+$ of live cells (n=3; two-tailed student's t test).

(E-F) Violin plots of splenocytes of mice treated with clodronate liposomes on day -2,0,2,5,7,9,12,15,17,20,22. Frequency of (E) CD11b$^+$ F4/80$^+$of live or (F) CD11b$^+$ CD11c$^+$ of live (n=3-4; two-tailed student's t test).

(G) Frequency of CD8$^+$ of CD3$^+$ cell (normalized to tumor weight) of CD45$^+$ fraction of wild-type TC1 tumors treated with clodronate liposomes on day -2,0,2,5,7,9,12,15,17,20,22, anti-PDL1 on day 10 and 17 and with DCvax-IT on day 9 and 16 isolated on day 23 after wild-type TC1 cells injection. P-values depict comparison to clodronate liposome treated mice (n=4-5; Mann-Whitney test).

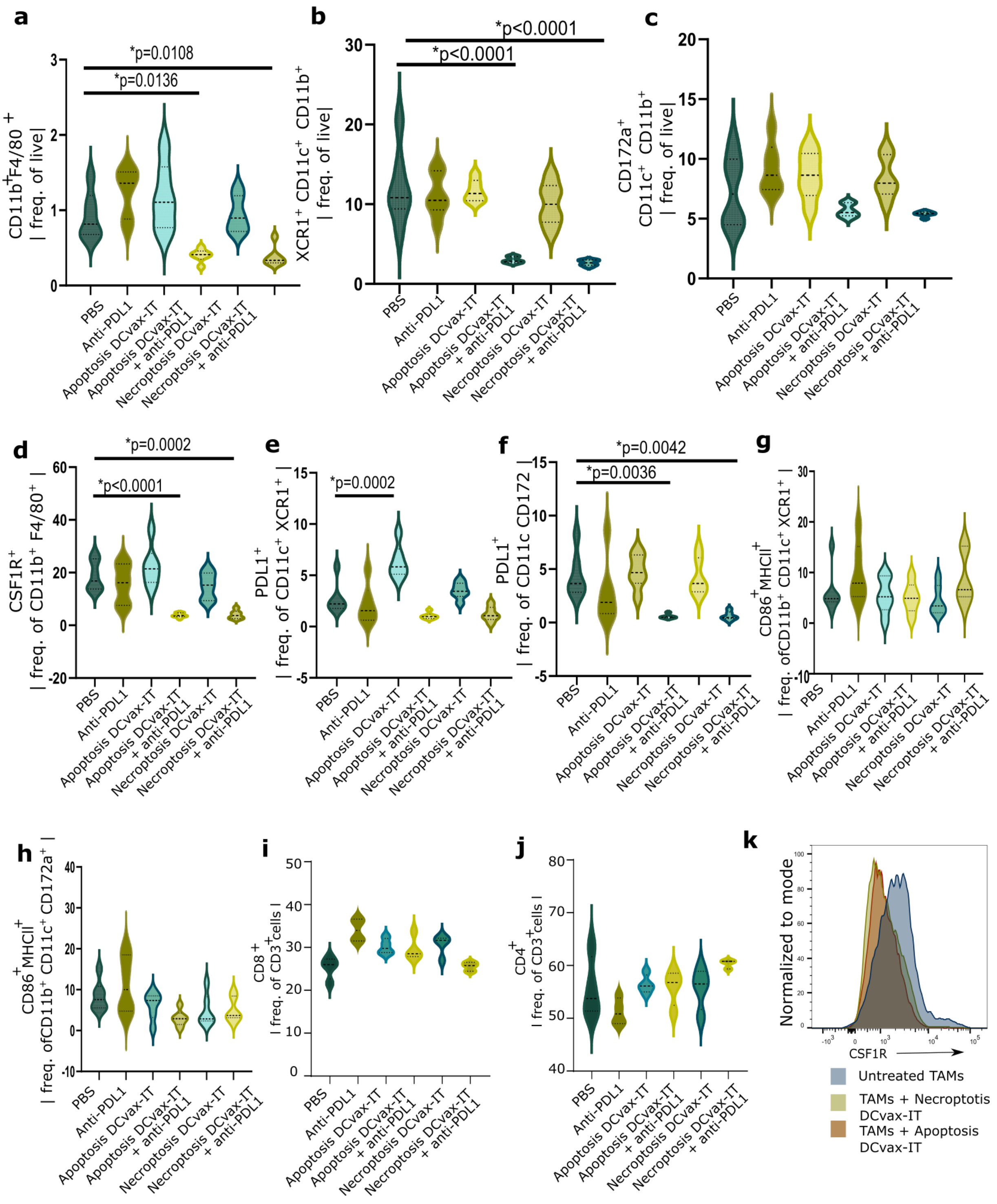


**Figure S14.**

(A-H) Lymph node analysis of wild-type TC1 tumor-bearing mice treated with DCvax-IT on day 9 and 16 and with anti-PDL1 on day 10 and 17. Frequency of (A)CD11b$^{+}$ F4/80$^{+}$ of live (B) CD11b$^{+}$ CD11c$^{+}$ XCR1$^{+}$ of live (C) CD11b$^{+}$ CD11c$^{+}$ CD172a$^{+}$ of live (D) CSF1R$^{+}$ of CD11b$^{+}$F4/80$^{+}$ (E) PDL1$^{+}$ of CD11b$^{+}$ CD11c$^{+}$ XCR1$^{+}$ (F) PDL1$^{+}$ of CD11b$^{+}$ CD11c$^{+}$ CD172a$^{+}$ (G) MHCII$^{+}$CD86$^{+}$ of CD11b$^{+}$ CD11c$^{+}$ XCR1$^{+}$ (H) MHCII$^{+}$CD86$^{+}$ of CD11b$^{+}$ CD11c$^{+}$ CD172a$^{+}$. p-values depict comparison to PBS treated mice (n=5-6; One-way ANOVA, Dunnett's multiple comparisons test).

(I-J) Lymph node analysis of wild-type TC1 tumor-bearing mice treated with treated with DCvax-IT on day 9 and with anti-PDL1 on day 10, isolated on day 12. Frequency of (I) $CD8^+$ of $CD3^+$ cells (J) $CD4^+$ of $CD3^+$ cells. P-values depict comparison to PBS treated mice (n=3-4; One-way ANOVA, Dunnett's multiple comparisons test).

(K) histogram of CSF1R on $CD11b^+$ $F4/80^+$ cells of wild-type TC1-derived TAMs-DCvax-IT cocultures.

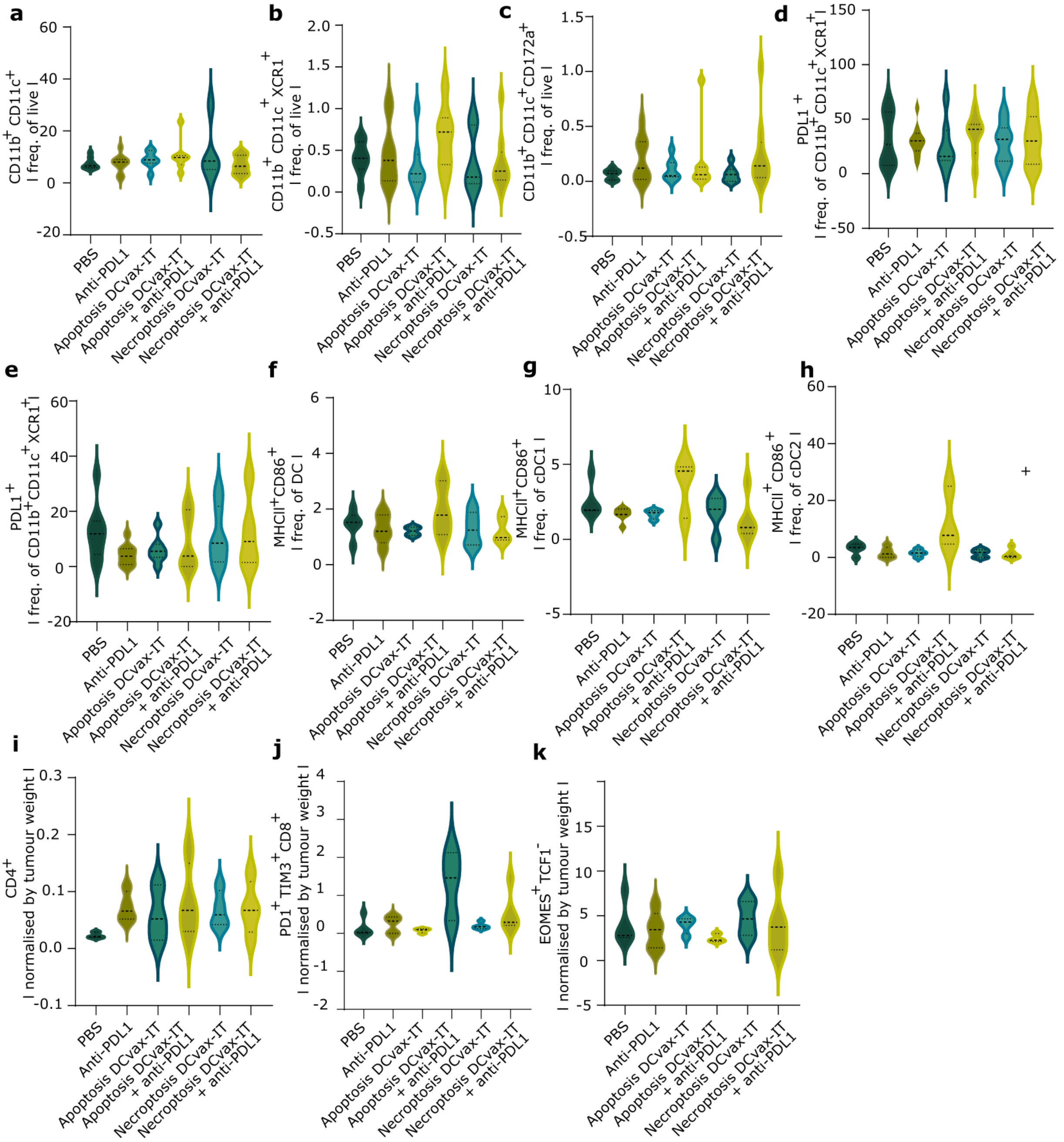


**Figure S15**

(A-H) TIL analysis of the $CD45^+$ fraction from wild-type TC1 tumor treated with DCvax-IT on day 9 and 16 and with anti-PDL1 on day 10 and 17, isolated on day 23 after wild-type TC1 cell injection. Violin plots of the percentage of (A) $CD11b^+CD11c^+$ of $CD3^-$ cells (B) $CD11b^+CD11c^+XCR1^+$ of $CD3^-$ cells (C) $CD11b^+CD11c^+CD172a^+$ of $CD3^-$ cells (D) $PDL1^+$ of $CD11b^+$ $CD11c^+XCR1^+$ cells (E) $PDL1^+$ of $CD11b^+$ $CD11c^+CD172a^+$ cells (F) $MHCII^+CD86^+$ of $CD11b^+$ $CD11c^+$ cells (G) $MHCII^+CD86^+$ of $CD11b^+$ $CD11c^+XCR1^+$ cells (H) $MHCII^+CD8$ of $CD11b^+$ $CD11c^+CD172a^+$ cells. Comparison to PBS treated mice (n=4-9; Mann-Whitney test).

(I-J) TIL analysis of the $CD45^+$ fraction from wild-type TC1 tumor treated with DCvax-IT on day 9 and 17 with or without anti-PDL1 injection on day 10 and 17 normalized by tumor weight at day of isolation. Frequency of (I) $CD4^+$ of $CD3^+$ (J) $PD1^+TIM3^+$ of $CD8^+$ cells (K) $EOMES^+$ $TCF1^-$ of $CD8^+$ cells. Comparison to PBS treated mice (n=3-5; Mann-Whitney test).

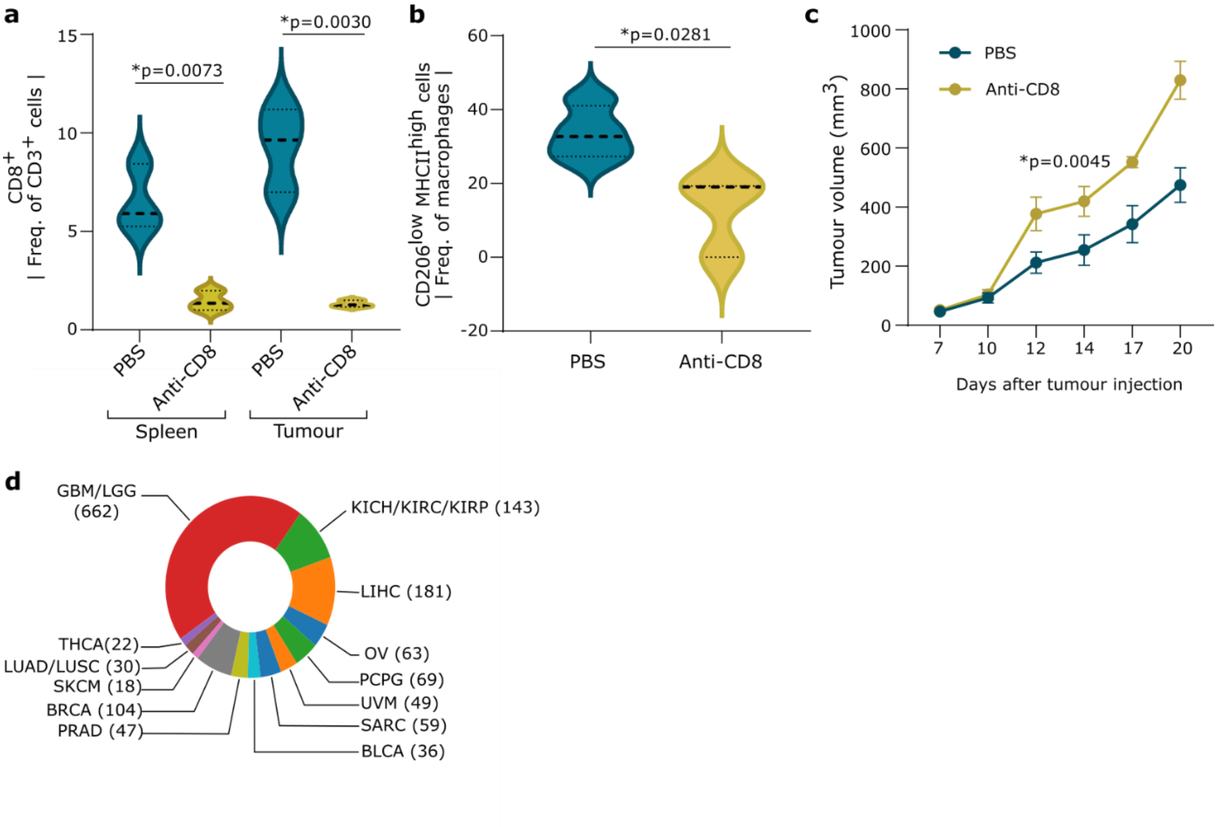


**Figure s16.**

(A) Frequency of $CD8^+$ of $CD3^+$ cells in the $CD45^+$ cell fraction of tumors isolated on day 23 after TC injection or spleen of untreated and anti-CD8 treated wild-type TC1 tumor bearing mice. P-values depict comparison to PBS treated mice (n=3; two-tailed student's t test).

(B) Frequency of $CD206^{low}MHCII^{high}$ in the $CD45^+$ cell fraction of tumors isolated on day 23 after TC injection of untreated and anti-CD8 treated wild-type TC1 tumors. (n=3; two-tailed student's t test).

(C) Tumor volume of untreated and anti-CD8 treated wild-type TC1 tumors. (n=3; area under curve; two-tailed student's t test).

(D) Cancer-type distribution analyses amongst all the TCGA C4/C5-tumours. (glioblastoma/low grade glioma GBM; n=622, kidney chromophobe/kidney renal clear cell carcinoma/kidney renal papillary cell carcinoma KICH/KIRC/KIRP; n=143, liver hepatocellular carcinoma LIHC; n=181, ovarian serous cystadenocarcinoma OV; n=63, pheochromocytoma PCPG; n=69, uveal melanoma UVM; n=49, sarcoma SARC; n=59, Bladder urothelial carcinoma BLCA; n=36, prostate adenocarcinoma PRAD; n=47, breast invasive carcinoma BRCA; n=104, skin cutaneous melanoma SKCM; n=18, lung adenocarcinoma/lung squamous cell carcinoma LUAD/LUSC; n=30, thyroid carcinoma THCA; n=22

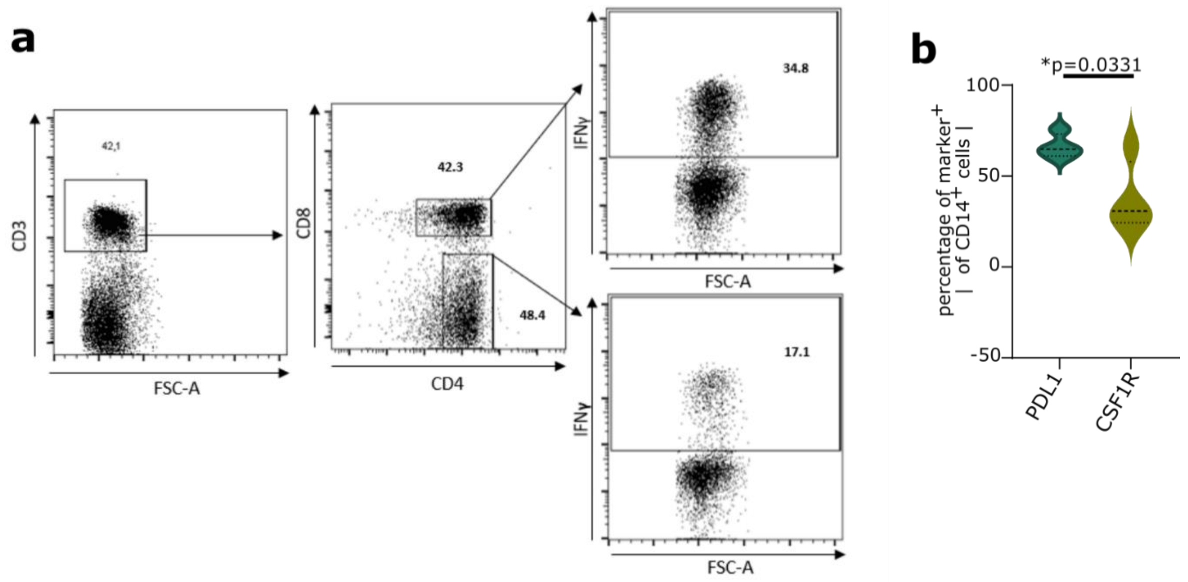


**Figure S17.**

(A) Representative gating strategy for IFNγ⁺ CD4⁺ and CD8⁺ T cells in CD45⁺ tumor-infiltrating leukocytes isolated from glioblastoma tumor samples.

(B) Frequency of PDL1⁺ or CSF1R⁺ of CD14⁺ macrophages isolated from glioblastoma tumor samples obtained at the day of resection at first diagnosis. (n=4; two-tailed student's t test).
