## Supplementary Methods for "A lymph node-to-tumour PDL1^+^macrophage circuit antagonizes dendritic cell immunotherapy"

### A lymph node-to-tumour PDL1$^{+}$macrophage circuit driven by dendritic cell immunotherapy inhibits anti-tumour immunity

#### Human study participants

Human samples were derived from glioblastoma patients included in the GlioVax trial (NCT03395587). Further trial details have been described elsewhere (Rapp et al., 2018).

#### Mice models

Wild type C57BL/6j were obtained from the KU Leuven breeding facility. The *Ifnar1*$^{-/-}$ mice (B6.129S2-*Ifnar1*$^{tm1Agt}$/Mmjax)(The Jackson Laboratory #010830) were a kind gift from the lab of Roos Vandenbroucke (VIB-Ugent) and the *Ccr7*$^{-/-}$ mice (*B6.129P2(C)-Ccr7*$^{tm1Rfor}$*/J)* (The Jackson Laboratory #006621) were a kind gift from the lab of Bart Lambrecht (VIB-UGent). All subcutaneous tumour experiments were done using 7- to 12-week-old female/male mice, maintained in the conventional mouse facility. Experiments were approved by the animal ethics committee at KU Leuven (project P114/2019 and p195/2020) following the European directive 2010/63/EU as amended by the Regulation (EU) 2019/1010 and the Flemish government decree of 17 February 2017.

#### Cell lines

TC1 wild type, Ripk3$^{-/-}$ , Mlkl$^{-/-}$ were a kind gift from the lab of Oliver Kepp (Université Paris) (Yang et al., 2016). Cells were cultured at 37°C under 5% $CO_2$ in DMEM media containing 2 mM L-glutamine, 3.7 g/L sodium bicarbonate, 4.5 g/L glucose and 1.0 mM sodium pyruvate) with 10% heat-inactivated fetal bovine serum (30 min at 56°C; FBS), penicillin 100 U/ml streptomycin 100 μg/ml DMEM and split when 90% confluency was reached through enzymatic dissociation (Trypsin).

#### Cell line creation

0.3 TU/cell LentiGuide Cas9 lentiviral particles was added. Thereafter, 48 hours later, the supernatant was replaced with fresh medium. The synthetic guide RNA complex was prepared by adding equal quantity of crRNA targeting *Casp8* + crRNA targeting *Casp9* + tracrRNA (25 nM), in 100 uL serum-free DMEM and these were incubated for 5 min at room temperature. 10 µL of DharmaFECT 1 in 100 µL of serum-free DMEM was incubated for 5 mins. The crRNA:tracrRNA complex and DharmaFECT 1 working solution were mixed and incubated for another 20 mins adding to lentiviral-transduced cells, which were incubated for another 48 hours to reach efficient gene knockout. The transfected cells were then sorted by a FACSAria cell sorter (BD Bioscience). Two weeks later, visible clones were picked and duplicated for seed reservation as well as western blotting detection of Casp8 (Thermofischer #PA5-77888) and Casp9(Thermofischer #PA5-16358) expression to confirm the knock-out phenotype.

##### *Cell death induction*

For cell death induction cells were seeded in the appropriate dish as such that they reached an 80% confluency the next day. Necroptosis was induced in TC1 WT by 100ng/mL tumor necrosis factor (TNF)

(Miltenyi #130-101-687), 1μM BV6 (Selleckchem #S7597 and 20μM Z-Val-Ala-Asp (OMe)-FMK (zVaD) (Bachem #4027403). For apoptosis, TC1 Mlkl$^{-/-}$, MC38 cells were pre-treated with 10μM BV6 for 3h and subsequently 100 ng/ml TNF was added. When applicable, a pre-treatment of 30μM of Necrostatin-1s (Bioke #17802S) for 3h was used to inhibit necroptosis.

***MTT assay***

TC1 WT and Mlkl$^{-/-}$ cells were seeded in a 96 well plate at a density of 5 000 cells per well 24h before cell death induction. Cell survival was obtained using the thiazolyl blue tetrazolium bromide (MTT) reagent (Abcam #ab197010). Absorbance was read on a microplate reader (Flexstation) at 490nm at 24h and 48h after cell death induction.

***RealTime-Glo Annexin V Apoptosis and Necrosis Assay***

A total of 5 000 cells TC1 WT and Mlkl$^{-/-}$ cells were plated in 96-well plates in 100 μl DMEM and incubated for 24h. Necroptotic and apoptotic stimuli cocktails were added to the plates. Apoptosis and necroptosis were detected using the RealTime-Glo Annexin V Apoptosis and Necrosis Assay (Promega #JA1011) including CellTox green (Promega #G8741). Bioluminescence and fluorescence at excitation 485nm and emission 530nm were measured at different time points; 1, 3, 6, 9, 20, 24, 27, 30 and 48 hours on a microplate reader (Flexstation). Fold changes to untreated (control) were taken. Percentage of cell death at 48h was estimated using following formula: Dead (%) = ((RFU TNF well/total RFU TNF well) − (RFU control well/total RFU control well)) × 100% as previously described (Degterev et al., 2014).

***Caspase 3/7 activity***

Cells were seeded in a 96-well plate at a density of 3000 cells/well. Cell death was induced and the Caspase 3/7 activity was measured at 24h and 48h using the caspase-glo kit (Promega #G8091). A fold change to untreated cells was taken to obtain caspase 3/7 activity.

***Western Blot***

For intracellular proteins, 800 000 cells were seeded in 10cm dishes 24h before cell death induction. Cells were scraped and collected at 1, 3, 6, 24 or 48 hours after cell death induction. Cells were centrifuged at 1500rpm for 5 min and the pellet was resuspended in 100μl NP-40 lysis buffer (Thermo Fischer #FNN0021) with protease (Thermo Fischer #A32953) and phosphatase inhibitors (Thermo Fischer #A32957). For secreted proteins, cells were 2 500 000 cells seeded in a 15cm dishes and media was collected 24h after cell death induction. Floating cells were removed by centrifugation. The supernatant was concentrated using Amicon Ultra-15 centrifuge filter units (Merck #UFC901024). A Bicinchoninic acid assay (BCA) for colorimetric quantification of total protein was done with a BCA protein assay kit (Thermo Fischer Scientific #23227) and a protein mixture of 90 μg was loaded onto the gel. Following proteins were detected; MLKL (clone 3H1), phosphorylated MLKL (pMLKL) (clone EPR9515(2)), RIPK1 (clone D94C12), RIPK3 (polyclonal), HMGB1(clone EPR3507), HPV-E7 (clone 8C9), PARP (clone 46D11) and β-Actin (clone AC-74 and AC-15). The primary antibodies were diluted in 5% BSA+TBST with a dilution factor of 1/1000. As secondary antibody, we used the anti-rabbit antibody labelled with HRP (Cell Signalling #7074S) or anti-mouse antibody labelled with HRP (Cell Signalling #7076S) diluted in 5% BSA+TBST with a dilution factor of 1/2000. The antibody's tethering to the proteins on the membranes were detected using the ECL substrate and the resulting precipitate was detected with the ChemiDoc MP imaging system. For detection of MLKL and HPV-E7, the membranes were stripped for 15 min at room temperature using stripping buffer (Abcam #Ab270550) with a dilution factor of 1/2000 and same procedure as described above was used to detect the protein of interest. The precision plus protein dual color standards (Biorad #161037) was used as a ladder.

***ATP secretion***

Media of TC1 WT and $Mlkl^{-/-}$ cells after cell death induction was harvested at 6, 24 and 48h. The presence of ATP in the media was analysed using the ATP assay system (Promega #FF2000). A fold change to untreated cells was taken to obtain ATP release.

***Flow cytometry-based detection of calreticulin and CD47***

A total of 200 000 TC1 WT and $Mlkl^{-/-}$ cells were plated in 12-well plates in 1 mL of DMEM and incubated for 24h. Cell death was induced for 48h. Then the cells were collected, washed with PBS and transferred to 5mL flow cytometry tubes. The cells were re-suspended in 50 µl of FACS buffer (0.5% BSA and PBS solution) and 1/100 anti-calreticulin primary antibody (clone B44). After 30 minutes incubation in the dark on ice, cells were washed with 1mL of FACS buffer, centrifuged and the supernatant was discarded. Then, cells were again re-suspended in 50 µl of FACS buffer with 1/500 of Goat Anti-Rabbit Alexa Fluor 488 (polyclonal) for CRT, anti-CD47 (clone miap301) antibody and Fixable Viability Dye eFluor 780. After 30 min incubation on ice in the dark, cells were fixed with cytofix (BD Bioscience #554655).

***DC vaccine formation***

For DC vaccine creation, bone marrow derived DCs were stimulated with dying cancer cells in a 1:1 ratio or with TC1 antigens (i.e., Human Papillomavirus (HPV) e6/e7 epitopes: VYDFAFRDL/DKKQRFHNI, RAHYNIVTF/LCVQSTHVD)), with or without 2.5 ng/ml interferon beta (IFNβ) (R&D sytems #8234-MB-010) for 48h. Where indicated, DCs were stimulated with 1000pg/ml lipopolysaccharide Escherichia coli (LPS-EB) (Invivogen #tlrl-eblps) as a positive control. After 48h, DCs were harvested by scraping and washed with PBS for injection; 1000 000 DCs per 100µl PBS or for further analysis. Media of the DC cultures was also collected for further analysis; ELISA, cytokine array.

***Flow cytometry***

Before staining procedure, FC receptor of all samples were blocked using TruStain FcX (Biolegend #101320) for 15min. Cells were further stained with the indicated antibodies listed in Key resource table, diluted in 0.5% BSA, for 1h and fixed with cytofix (BD Bioscience #554655). In case of intracellular markers, cells were fixed and permeabilised with the Cytofix/Cytoperm Kit (BD Bioscience #554714). For the staining of transcription factors, the true-nuclear transcription factor buffer (Biolegend #42441) set was used. After fixation, cells were maintained in 0.5% BSA. For intracellular cytokine staining, the cells were stimulated with Dynabeads Mouse T activator CD3/CD28 (Thermo Fisher Scientific #11456D). After 1h at 37°C 5% $CO_2$, 2µl Brefeldin A (Thermo Fisher Scientific #00-4506-51) was added. Cells were then placed at 37°C 5% $CO_2$ for 4h, transferred to 4°C overnight and then stained for intracellular cytokines. Flow cytometry was performed on the Attune NxT (Thermo Fisher Scientific), FACSCanto (BD Bioscience) or the ID7000 (SONY). Cell doublets were excluded based on FSC-A/FSC-H. Flow cytometry data was analysed using FlowJo.

***DC vaccine - TAM cocultures***

DC vaccines were harvested as previously described. DC vaccines were cocultures with TC1 tumour derived TAMs in a 1:1 ratio. After 48h cocultures were scraped to collect the cells, centrifuged and washed with PBS. Single cell suspension was stained with fluorescently labelled antibodies diluted in FACS buffer (0.5% BSA and PBS solution) for 1hour on ice and then washed with the same buffer. Cells were fixed with cytofix (BD Bioscience #554655). Following fluorochrome-conjugated antibody clones were used to analyse the isolated lymph nodes by flowcytometry; MHCII (clone M5/114.15.2), PDL1 (clone 10F.9G2), CD86 (clone GL1), F4/80 (clone T45-2342), CSF1R (clone AFS98), CD11b (clone ICRF44) and CD206 (clone C068C2).

***T cell - TAM cocultures***

TC1 derived TAMs and matched T cells (derived from the spleen) were harvested as described above. TAMs were plated in media alone or with 10µg/ml anti-PDL1(Polpharma Biologics), or 10µg/ml anti-PD1(Polpharma Biologics), for 24h. TAMs were scraped and washed and cocultured with their matched T cells in a 1:1 ratio with 100 IU/mL IL-2 for 48h. Then cocultures were scraped to collect the cells, centrifuged and washed with PBS. Single cell suspension was stained with fluorescently labelled antibodies diluted in FACS buffer (0.5% BSA and PBS solution) for 1hour on ice and then washed with the same buffer. Cells were fixed with cytofix (BD Bioscience #554655). The following fluorochrome-conjugated antibody clones were used: CD3 (clone 17A2), CD4 (clone GK1.5) and CD8a (clone 53-6.7).

***Enzyme-linked immunoassay (ELISA)***

Derived media of DC vaccines as described above was used to perform a CXCL10 ELISA (R&D systems#DY466. For IFNα and β secretion, TC1 cells were treated with 1µg/ml LPS (Invivogen# tlrl-eblps), 100µg/ml imiquimod (Invivogen #tlrl-imqs), 10µg/ml 5'ppp-dsRNA/lyovec (Invivogen #tlrl-3prnaclv), 2'3' cGAMP (Invivogen #tlrl-nacga23) 25µM doxorubicin (Merck #D1515) or 100µM cisplatin

(Merck #PHR1624) for 24h. An ELISA for IFN alfa (Invivogen #luex-mifnav2) and beta (Invivogen #luex-mifnbv2) was performed according to manufacturer's protocol.

***Quantitative polymerase chain reaction (qPCR)***

The RNA of DC vaccines was extracted using the Purelink RNA Mini Kit (ThermoFischer #12183025). Using the QuantiTect Reverse Transcription kit (Qiagen #205313) cDNA was synthesized from RNA. The qPCR was performed on the StepOnePlus Real-Time PCR system (Applied Biosystems) using SYBRgreen (Highqu #QPD0150) with the following primers; *Irf7, Rsad, Mx1, Cxcl9, Cxcl10* and *Actin* (all primers were ordered at Integrated DNA Technologies, sequences available in Table 1). Fold change was determined using the 2^ΔΔCT method compared to the house keeping gene, *Actin,* and untreated DCs.

**Antibody array**

Derived media of DC vaccines as described above was used for a mouse cytokine array panel A (R&D systems #ARY006) was used. Cytokine array was performed according to the manufacturer's protocol. Arrays were read on the ChemiDoc (Biorad). Dot intensities were determined using Image Lab (Biorad). From all values, the background was subtracted. Normalisation was done using DCs stimulated with live cancer cells. Fold-change values derived from above antibody array for different immunological factors, cytokines or chemokines per treatment condition were used to run a GSEA analysis using WebGestalt (WEB-based Gene SeT AnaLysis Toolkit), which is a functional enrichment analysis web tool (Liao et al., 2019). This GSEA analysis was run with Mus musculus as reference organism for Gene Ontology Biological Process term enrichment with the following analyses parameters: minimum number of genes for a category, 3; maximum number of genes for a category, 2000; significance level, top 10; number of permutations, 1000; p, 1; collapse method, mean; number of categories expected from set cover, 10.

**Efferocytosis assay**

30 000 bone marrow derived dendritic cells were plated in white clear bottom 96 well plate (Corning #3610) per well. Untreated or dying cancer cells were collected 24h after cell death induction. Cells were stained with 20 µg/mL pHRodo (ThermoFischer #P35373) for 1h and washed with FBS. Cells were added to BMDCs in a 1:1 ratio. Plates were kept in 37°C under 5% $CO_2$ or in 4°C as a negative control. Fluorescence at excitation 490nm and emission 520nm were measured at different time points; 8, 20, 24, 30, 40 and 48 hours after co-incubation on a microplate reader (BioTek). For the efferocytosis index calculations, fluorescent intensity values at 37 °C were subtracted by the appropriate 4°C negative control.

**5-bromo-2'-deoxyuridine assay (BrdU-assay)**

Isolated glioblastoma TAMs were plated with graded cell numbers from 20.000 to 100 cells in a dilutional series in 200 µL/well in a 96-flat-bottom plate and incubated overnight at 37°C and 5% $CO_2$. The following day, $1x10^5$ cells/well allogeneic CD14-depleted PBMC were added in 100 µl resulting in a final volume of 200 µl per well. Additionally, anti-PD-L1 (500 ng/ml) was added to some of the cultures to analyze its effect on macrophage-stimulated T-cell proliferation. X-Vivo 15 medium as well as PBMC ($1x10^5$ cells) or macrophages ($2x10^5$ cells) alone were used as negative controls. After five days of culture, cells were "pulsed" with 20 µl BrdU/well (BD Bioscience #550891) previously diluted 1:100 in X-Vivo-15 medium and incubated for 16 to 24 hours at 37°C and 5% $CO_2$ to allow the base analogue BrdU to be incorporated in place of thymidine during the cell division of the proliferating cells. After incubation, cells were washed and fixed with 200 µl/well FixDenat for 30 minutes, followed by an incubation with 100 µL/well of a 1:100 diluted anti-BrdU antibody for 90 min, during which the peroxidase-conjugated antibody binds to the BrdU incorporated into the newly synthesized DNA. After discarding the antibody, cells were washed three times with 200 µl/well washing buffer diluted 1:10 in distilled water to remove non-specifically bound antibody. Cells were then incubated for a maximum of 30 minutes with 100 µL/well of a substrate solution (tetramethylbenzidine - TMB), which is converted to a blue dye by the peroxidase bound to the BrdU-anti-BrdU complex. The optical density was analyzed using an ELISA reader (660 nm versus 490 nm).

***Murine bone marrow-derived dendritic cell and macrophage generation***

Bone marrow was isolated from wild type C57BL/6j, $Ifnar^{-/-}$ or $Ccr7^{-/-}$ mice. Both the femur and tibia were flushed using PBS and the cell suspension was centrifuged for 5min at 1500rpm. The pellet was resuspended in red blood cell lysis buffer (Merck life science), incubated for 5 min and centrifuged. Cells were resuspended in RPMI supplemented with 100 u/mL penicillin, 100 µg/L streptomycin, and 10% heat-inactivated fetal bovine serum (FBS). Bone marrow derived cells were differentiated into macrophages by adding 25 ng/ml M-CSF (Peprotech #315-02) into the media for 6 days. For differentiation into DCs, 20 ng/ml GM-CSF (Peprotech #315-03) and 10 ng/ml IL4 (Peprotech #214-14) was added to the media for 7 days. Media was replenished after 3 days. Differentiation of dendritic cells and macrophages was confirmed by flowcytometry with respectively following fluorochrome-conjugated antibody clones were used; CD11b (clone M1/70), CD11c (clone N418) and CD11b (clone M1/70), F4/80 (clone BM8).

***Murine splenocytes and T cells isolation***

Splenocytes were obtained from the spleen from wild type C57BL/6j mice. Spleens were minced and filtered through a 70-micron cell strainer. Cells were incubated in red blood cell lysis buffer for 5 min and centrifugated. For T cell isolation, splenocytes were purified using the negative selection pan-T cell isolation kit II (Miltenyi #130-095-130). Cells were maintained in RPMI supplemented with 100 u/mL penicillin, 100 µg/L streptomycin,10% heat-inactivated fetal bovine serum (FBS) and 100 IU/mL IL-2 (Peprotech #210-12). Following fluorochrome-conjugated antibody clones were used to confirm isolation of T cells by flowcytometry; CD3 (clone 17A2), CD4 (clone GK1.5) and CD8a (clone 53-6.7).

***Murine lymph node isolation***

3 days after DCvax-IT vaccination, mice were sacrificed and the axillary and inguinal lymph nodes of both sides were isolated. To create a single cell suspension, lymph nodes were minced and filtered through a

70-micron cell strainer and analysed by flow cytometry immediately. Following fluorochrome-conjugated antibody clones were used to analyse the isolated lymph nodes by flowcytometry; CD11c (clone N418), XCR1 (clone ZET), CD172A (clone P84), MHCII (clone M5/114.15.2), PDL1 (clone 10F.9G2), PD1 (clone RMP1-30), CD86 (clone GL1), F4/80 (clone T45-2342), CSF1R (clone AFS98), CD11b (clone ICRF44) and CD206 (clone C068C2).

***Murine tumour infiltrating leukocytes (TILs) and Tumour associated macrophages (TAMs) isolation***

Tumours were isolated at day 23 after tumour injection. A single cell suspension was made, using the tumour dissociation kit (Miltenyi #130-096-730). TILs were isolated through magnetic bead separation via CD45 (Miltenyi #130-110-618). For TAM isolation, anti-F4/80 microbeads (Miltenyi #130-110-443) were used. Isolated TILs and TAMs were either maintained in RPMI supplemented with 100 u/mL penicillin, 100 µg/L streptomycin, 10% heat-inactivated fetal bovine serum (FBS) or stained for flow cytometry immediately. Following fluorochrome-conjugated antibody clones were used; FOXP3 (clone MF-14), GATA3 (clone TWAJ), CD62L (clone MEL-14), TCF1/7 (clone S33-966), CD107a (clone 1D4B), TOX (clone REA473), Tbet (clone O4-46), CD3 (clone 17A2), CD8a (clone 53-6.7), Ki-67 (clone 11F6), CD45 (clone 30-F11), Eomes (clone Dan11mag), CD127 (clone SB/199), PD1 (clone RMP1-30), CD4 (clone GK1.5), TIM3 (clone 5D12/TIM-3), IL2 (clone JES6-5H4), TNFα (clone MP6-XT22), IFNγ (clone XMG1.2), PDL1 (clone 10F.9G2), PD1 (clone 29F.1A12), Siglec H (clone 551), CD11c (clone N418), XCR1 (clone ZET), CD172A (clone P84), MHCII (clone M5/114.15.2), PDL1 (clone 10F.9G2), PD1 (clone RMP1-30), CD86 (clone GL1), F4/80 (clone T45-2342), CSF1R (clone AFS98), CD11b (clone ICRF44) and CD206 (clone C068C2).

***Human TIL isolation***

For TIL isolation, surgically resected tumour samples of GBM patients at first diagnosis (primary GBM) or at recurrence after dendritic cell vaccination were collected and stored in MACS tissue storage solution (Miltenyi #130-100-008) for a maximum of 24 h at 4°C.  TILs were isolated from tumour samples by the tumour dissociation kit (Miltenyi #130-095-929) and magnetic bead isolation using $CD45^+$ beads (Miltenyi #130-045-801). For flow cytometric analysis, the $CD45^+$ cells were stained for T cell and TAM characterizing markers: CD3 (clone OKT3 or UCHT1), CD4 (clone okt/04 or REA636)), CD8 (clone RPA-T8 or Sk1 or BW135/80), CD45RO (clone UCHL1), CD45 (clone REA747), CD163 (clone REA812), CD45PcP (clone 2D1), CD14 (clone 63D3), CSF1R (clone AFS98), CD274 (clone M1H1), isotype control (clone X40 or MOPC-21 or 27-35 or P3.6.2.8.1) for 15 min at 4°C, washed with phosphate-buffered saline (PBS), fixed with 4% d1) paraformaldehyde (PFA) and analyzed on a Cytoflex flow cytometer, using CytExpert 2.3 software (Beckman-Coulter).

For detection of interferon-γ, TILs were stimulated in X-Vivo 15 medium (Lonza #BE02-060F) with 5 ng/mL phorbol myristate acetate (PMA) (Sigma-Aldrich #P8139) and 500 ng/mL Ionomycin (Sigma-Aldrich #I9657) for 5 h in the presence of 5 µg/mL Brefeldin A (BioLegend#420601) at 37°C and 5% $CO_2$ in a humidified atmosphere. After stimulation, cells were washed with PBS, stained for CD3, CD4, CD8 and CD45RO for 15 min at 4°C, washed with PBS and fixed for 15 min with 4% PFA. Subsequently, cells were permeabilised with 0,05% saponin in PBS (Sigma-Aldrich), stained intracellularly with: IFNγ (clone 4S.B3), CD3 (clone OKT3 or UCHT1), CD4 (clone okt/04 or REA636)), CD8 (clone RPA-T8 or Sk1 or BW135/80) and isotype control (clone X40 or MOPC-21 or 27-35 or P3.6.2.8.1) for 15 min and subjected to flow cytometric analysis.

**Human CD14$^+$ CD163$^+$ macrophages isolation**

To separate macrophages from tumor-infiltrating leukocytes, cells were adjusted to a cell titer of 2.5x $10^7$cells/mL in 0.5% HSA/D-PBS and stained with monoclonal antibodies directed against CD14 (clone 63D3) and CD163 (clone REA812). First, myeloid cells were identified by their typical FSC vs SSC. Doublets are excluded via a SSC (W) vs SSC (H) gating. Finally, within this single cell population, cells were gated based on their CD14 and CD163 expression, thereby targeting the double-positive population using an unstained control. This population was then sorted with a MoFlo XDP Sorter (Beckmann Coulter) into a 5 ml round bottom tube containing 1mL 100% FCS.

#### In Vivo Experiments

***Mouse experiments***

Seven to twelve-week-old female/male C57BL/6J mice were subcutaneously (s.c) injected with 1x$10^6$ TC1 or MC38 cells. For prophylactic vaccinations, 1x$10^6$ DCs were injected s.c twice (one week apart), before TC1 tumour inoculation. Mice were rechallenged with 1x$10^6$ TC1 cells after 30 days. For curative vaccinations, 1x$10^6$ DCs were injected s.c on day 9 and 16 after TC1 inoculation. When applicable, mice were cotreated with 250 µg of anti-PDL1 (clone MIH5; Polpharma Biologics), anti-CTLA4 (clone 4F10; Polpharma Biologics), anti-CSF1R (AFS98; BioXCell), or anti-PD1(RMP1-14; Polpharma Biologics), on day 10 and 17 via intraperitoneal (i.p.) injections. For CD8 depletion experiments, mice were given 200 µg of anti-CD8 (clone YTS169; Polpharma Biologics), i.p. one day before tumour inoculation and then every other day until the tumour reached 500 $mm^3$. When applicable, cisplatin (8mg/kg) (Sigma #p4394) was given on day 9 and 16. As indicated, 200µl clodronate liposomes (Liposoma) was given one day before tumour injected and every 2/3 days subsequently. Mice were monitored and weighed every other day and tumour volume was determined by height x width x length.

***CM-DIL***

DC vaccines were stained with 1µL/$10^6$ DCs of CellTracker CM-Dil dye (Thermofischer #C7000) prior to DC vaccination. 72 hours after vaccination mice were euthanised and both the left and right axillary and inguinal lymph nodes were isolated. Single cell suspension was stained with the following fluorochrome-conjugated antibody clones: CD11b (clone M1/70), CD11c (clone N418), F4/80 (clone QA17A29), XCR1 (clone ZET), CD172A (P84) and Siglec-H (551).

#### In silico Analyses

##### Transcriptomic analysis with The Cancer Genome Atlas (TCGA) datasets

##### Immuno-transcriptomic analyses

We used the bulk transcriptomes of TCGA as pre-processed by the Toil-recompute project(Vivian et al., 2017) in Xena(Goldman et al., 2020) Immune cell proportions were inferred using the Quantiseq approach for immune cell deconvolution(Finotello et al., 2019). Filtering by Overlapping the Thorsson immune subtype class labels(Thorsson et al., 2018) from TCGA pan-cancer analyses (C1 to C6; see results for more details), resulted in 9126 samples in total. Overall z-score normalized enrichment analyses was performed for each respective immune cell-fractions or as ratios per immune subtype class, specified in the figure legends and the text. Survival endpoints, were obtained from the same Xena Toil-Recompute TCGA data hub. A radar plot representation visualising the Spearman's correlations between *CD274* gene expression levels in M1 or M2 macrophage fractions, was built using all patients from same TCGA data for which C1-C6 immune subtype class labels were available.

##### Extracellular network analyses

The extracellular network analyses was performed following the CRI iAtlas portal tools and methodology (Eddy et al., 2020). Herein, this network analyses uses a database of well-established ligand-receptor, cell-receptor, and cell-ligand pairs previously published elsewhere (Ramilowski et al., 2015) and retrieved via FANTOM5(Ramilowski et al., 2015)). therefore, the network is built based on three criteria i.e., selected TCGA immune subtype class labels, Abundance threshold (%) and Concordance or Discordance threshold. Since the aim of the analyses was to reveal features of T cell-hostile tumours, it was hence restricted to only C4/C5 TCGA class-labels. hence, abundance threshold creates a selection threshold for the nodes in the network wherein the abundance threshold is the frequency of samples in the upper 2 tertiles of cell-abundance or gene expression-distributions. Here we used the default abundance threshold of 66%. Finally, concordance threshold values are relevant for the inclusion of the edges in the network between any two relevant nodes. Concordance threshold is computed based on a contingency table consisting of the ternary values of the two applicable nodes for an edge. However, we pre-programmed this threshold to capture discordance probability i.e., an edge with a concordance score of 0.5 that is considered discordant because the analyses with threshold of 0.5 captures information on one node being highly expressed while the other being lowly expressed (thus, discordant or uncorrelated) rather than both being simultaneously highly or lowly expressed.

##### GISTIC analysis

For genes derived from above extracellular network analyses (TNF, EDN3, IL13, EDN1, CX3CL1, VEGFB, IFNA2, IFNB1, IFNG, HMGB1, IL1A, IL1B), we performed high resolution copy number variation (CNV) analyses using the TCGA's genomic data via the GISTIC toolset poral (Mermel et al., 2011) (https://portals.broadinstitute.org/tcga/home). we used the Gene-Centric GISTIC Analyses wherein, amplifications and deletion tendencies were separately analysed per gene across a standardized pan-cancer TCGA dataset (i.e., 2013-08-16 TCGA Pan-Cancer data set), consisting of data from 4934 primary tumour samples passing GISTIC's default's quality control criteria from following cancer-types: bladder cancer (136), breast cancer (880), colorectal adenocarcinomas (585), glioblastoma multiforme (580), head and neck squamous cell cancer (310), kidney clear cell carcinoma (497), acute myeloid leukemia (200), lung adenocarcinoma (357), lung squamous cell carcimoma (344), ovarian serous carcinoma (563), and uterine endometrial carcinoma (496).

##### Prognostic impact analysis

For prognostic analyses i.e., impact of co-association between *CD274* and macrophage immune-fractions (TIMER-derived) on overall survival in multiple TCGA cancer-datasets spanning 19 cancer-types and 8493 patients, we utilized the TIMER2.0 portal (Li et al., 2017, 2020). the "Outcome module" was used for estimating the prognostic impact along with corrections for multiple covariates (stage, age, gender) in a multivariable Cox proportional hazard model. The hazard ratio from the Cox model was then computed into z-scores herein z-score > 0 meant negative prognostic impact (shorter survival) while z-score < 0 meant positive prognostic impact (prolonged survival). For this analyses, the following TCGA cancer datasets were utilized (number of patients in brackets): BLCA (n=408), BRCA (n=1100), COAD (n=458), GBM (n=153), HNSC-HPV- (n=422), HNSC-HPV+ (n=98), KICH (n=66), KIRC (n=533), KIRP (n=290), LGG (n=516), LIHC (n=371), LUAD (n=515), LUSC (n=501), OV (n=303), PAAD (n=179), PCPG (n=181), PRAD (n=498), READ (n=166), SARC (n=260), SKCM (n=471), STAD (n=415), THCA (n=509), UVM (n=80).

#### Immuno-oncology clinical trials analyses

Gene expression data for 12 clinical trials spanning 1142 patients in 5 cancer-types(Van Allen et al., 2015; Atkins et al., 2017; Choueiri et al., 2015; Cloughesy et al., 2019; Gide et al., 2019; Hoffman-Censits et al., 2016; Hugo et al., 2016; Kim et al., 2018; Liu et al., 2019; Miao et al., 2018; Riaz et al., 2017; Zhao et al., 2019), was downloaded from the Synapse server associated with the CRI iAtlas portal (syn24200710) (Eddy et al., 2020). We applied the immune subtype classifier (C1-C6) from the original paper (https://github.com/CRI-iAtlas/ImmuneSubtypeClassifier) on the normalized gene expression data to infer the immune subtypes. Cancer immune subtypes C2/3/6 and C4/5 were merged as separate categories and differences in patient survival (OS) were investigated using lifelines 0.27.0 in Python 3.9.5. Statistical significance was investigated using the log-rank test.

#### Transcriptomics analyses for murine subcutaneous tumours' bulk-RNAseq data

Affymetrix Mouse Exon 1.0 ST Array data, profiled from subcutaneous tumours based on B16-F10, TC1, CT26, MC38, LL2/LLC, RENCA, 4T1, TRAMPC1, EL4, P815 or PAN02 murine cancer cell lines implanted in syngeneic mice backgrounds, was derived from an existing published study (i.e., GSE85509) (Mosely et al., 2017) and analysed for expression levels of following metagene-signatures: pro-lymphocytic IFNγ/effector signalling, macrophages, type I IFN/ISG-response, or DCs (Mosely et al., 2017; Sprooten et al., 2021), and represented as a row-normalized heatmap.

In another case, for differential gene-expression (DGE) analyses, above transcriptomics data for TC1-tumours and MC38-tumours were analysed in R 4.2.0, using the package oligo 1.60.0(Carvalho and Irizarry, 2010). All corresponding CEL files were imported and RMA-corrected, then summarized to the gene level for which identifiers were mapped based on the NetAffx CSV files provided by Affymetrix (MoEx-1_0-st-v1.na28.mm9.transcript.csv). For the pairwise comparison between TC1 and MC38, we used limma 3.52.2(Ritchie et al., 2015) and applied t-tests with empirical Bayesian shrinkage of variance. P-values were adjusted in accordance with the Benjamini-Hochberg method.

Finally, for Correlation gene-set enrichment analyses (GSEA) analyses, following assorted genes specifically enriched in the TC1-tumours (compared to MC38-tumours) from above DGE analyses, that we further annotated as myeloid/macrophage-signalling relevant based on literature were utilized: *Dkk2, Figf, Akr1c18, Ptgs1, Ereg, Thbs1, Thbs2, Ptgs2, Aspa, Pla2g7, Pdpn, Pla2g4a, Areg, Cd80, Nox4, Abca1, Cd33, Il1rl1, P2ry5, Axl, Mrc1, Gas2, Ctsl.* Correlation GSEA analyses of this gene-set (gene-set 1) versus well-established translationally-relevant markers of tumour-associated macrophages (*Cx3cr1, Ccr2, Tek, Csf1r, Cd274, Pdcd1, Sirpa, Ido1, Pdcd1lg2, Cd40* (Duan and Luo, 2021; Liu and Wang, 2020)) (gene-set 2) was performed against the background of a reference murine macrophage transcriptomic-profile, using the standardized work-flow of Immuno-Navigator portal (Vandenbon et al., 2016).

**Maturation trajectory analyses for single-DC vaccine's transcriptomes**

Trajectory analyses of monocytic-derived DC vaccines micro-array transcriptomes was performed using the Gene Expression Omnibus (https://www.ncbi.nlm.nih.gov/geo/) dataset (GSE85698)(Castiello et al., 2017). This study included 93 DC vaccines that were administered to 18 prostate carcinoma (PDAC) patients. Each patient received multiple lots of autologous DC. Vaccination regimen included 5 vaccinations (one every 3 weeks) with boost vaccinations in case of response. At least 2 vaccines per patient are included. Normalized microarray expression dataset was extracted from GEO portal and further processed to filter mitochondrial genes and non-protein coding genes. Microarray's duplicated annotation probes was collapsed by per-gene average ratio of probe's expression values. Initial dimentionality reduction to further calculate trajectory inference was performed using high variable genes with additional threshold parameters of loess fraction 0.1 ad number of genes <= 4000. Hence UMAP dimensionality reduction was based on the top 9 principal components which covered 99.9% of genetic variance and was computed on a 2-dimensional scale of 4 neighbours. Trajectory inference was performed using STREAM (Huidong Chen, et al, Nature Comm, 2019) on the precomputed dimensionality reduction using *"seed elastic principal graph"* algorithm with default paramaters except for *4 neighbours* and *"ap" clustering* algorithm. Further branch pruning was performed using *"elastic principal graph"* with default parameters except for epg_alpha 0.01, epg_mu 0.2, epg_lambda 0.03, epg_n_nodes 5 and incr_n_nodes 3. Signature expression of several signatures was calculated on per DC vaccine sample of average expression genetic profiles. Leaf markers were calculated using "detect leaf markers" function against root S0 group using default parameters except for cutoff_zscore 1.0, cutoff_pvalue 1.0, min_num_cells 5 and percentile_expr 99. Pathway analysis for each leaf marker genes was computed using gseapy with signal to noise methodology of gene set permutation type for a maximum of 2000 permutations, including Reactome pathways between 1 and 2000 genes. Top ranked and biologically relevant Reactome pathways were plotted based on thir Normalized Enrichment Score (NES) and FDR values.

**Analysis of cancer patient's single-cell RNA-sequencing (scRNAseq) datasets**

For analyses of *CD274* or *Cd274* expression at single-cell resolution on pan-tumour level, various scRNAseq pan-tumour maps from existing peer-reviewed literature were accessed from the Tumor Immune Single Cell Hub (Sun et al., 2021), a large-scale curated database integrating single-cell transcriptomic data from >2 million single-cells (with a uniform/standardized workflow accounting for quality control and batch-effects) across >75 high-quality tumour-derived datasets. We used the Gene Exploration module of TISCH to short-list scRNASeq datasets with stable expression profiling available for at least most of the following tumour-associated cells: $CD4^{+}T$ cells, $CD8^{+}T$ exhausted cells, NK cells,

CD8$^{+}$T cells in general, fibroblasts, endothelial cells, B cells, malignant/cancer cells, DCs and macrophages. This delineated 21 eligible single-cell (sc)RNAseq datasets spanning 13 diverse cancer-types (SARC, CRC, SKCM, HNSC, UVM, Glioma/GBM, CHOL, OV, LIHC, NSCLC, BRCA, PAAD, BCC), 287 patients and 2 species (humans or mice), further accessible here: GSE119352, GSE122969, GSE120575, GSE103322, GSE120909, GSE139829, GSE123139, GSE139324, GSE141982, GSE125449, GSE118828, GSE131907, GSE139555, GSE136206, GSE111672, GSE131928, GSE146771, GSE115978, GSE99254, GSE110686, GSE123813. Across these datasets, we derived average gene expression of *CD274* or *Cd274* across an entire cell-type cluster per scRNAseq dataset and integrated all the values to create row-normalized heatmaps.

##### Predictive analyses of PDL1$^{+}$TAMs signature in immuno-oncology clinical trials

Pre-treatment tumour-derived gene expression data for 454 cancer patients (185 responders vs. 269 non-responders) treated with anti-PDL1 ICB (atezolizumab or durvalumab) spanning 5 cancer-types (4 patients with ureter/renal pelvis cancer, 345 with urothelial cancer, 31 with bladder cancer, 72 with esophageal cancer and 2 with renal cell carcinoma), or 761 cancer patients (504 responders vs. 257 non-responders) treated with anti-PD1 ICB (nivolumab or pembrolizumab) spanning 11 cancer-types (39 patients with lung cancer, 14 with glioblastoma, 7 with ureter/renal pelvis cancer, 45 with gastric cancer, 5 with colorectal cancer, 415 with melanoma, 59 with bladder cancer, 22 with hepatocellular carcinoma, 14 with breast cancer, 31 with renal cell carcinoma, and 110 with head & neck cancer) were accessed from the ROC Plotter server (Fekete and Győrffy, 2019; Kovács and Győrffy, 2022). We interrogated the impact of a PDL1$^{+}$TAM metagene (composed of *CD68, CD163, CD14, CD274*) on tumour-level objective response rate (ORR)-based responders vs. non-responders for either anti-PDL1 or anti-PD1 ICB treatments. Analyses was restricted to pre-treatment transcriptome while excluding any statistical outliers. Statistical significance was investigated using the Mann-Whitney U test.

##### Statistics

The statistical details of all the analyses are reported either in the figure legends, figures and/or methods sections, including statistical analysis performed, statistical significance thresholds/values and in most cases the counts/number of data-points. All the statistical tests used herein were always two-tailed unless otherwise explicitly mentioned. Gene signatures were estimated by considering the average expression of all the genes within that signature, unless otherwise mentioned. Details about used software for analysis can be found in the key resources table.

#### Study approval

Human samples were obtained with the informed consent of the patients and approval by the ethical committee of University Hospital Düsseldorf (MC-LKP-921 and 3005). Mouse Experiments were approved by the animal ethics committee at KU Leuven (project P114/2019 and p195/2020) following the European directive 2010/63/EU as amended by the Regulation (EU) 2019/1010 and the Flemish government decree of 17 February 2017.
