## Supplementary material for "A lymph node-to-tumour PDL1^+^macrophage circuit antagonizes dendritic cell immunotherapy": Key Resources

| Reagent or resorce | Source | Identifier |
|---|---|---|
| antibodies | | |
| Alexa Fluor 488 anti-mouse CCR7 (clone:4B12) | Biolegend | Cat#120110 RRID:AB_492841 |
| APC/Cyanine7 anti-mouse CD3 (clone:17A2) | Biolegend | Cat#100222 RRID:AB_2242784 |
| BV750 anti-mouse CD3 (clone:17A2) | Biolegend | Cat#100249 RRID:AB_2734148 |
| PE/Cyanine7 anti-mouse CD4 (clone: GK1.5) | Biolegend | Cat#100422 RRID:AB_312707 |
| PE anti-mouse CD4 (clone: GK1.5) | Biolegend | Cat#100408 RRID:AB_312693 |
| BUV563 anti-mouse CD4 (clone:GK1.5) | BD Biosciences | Cat#612923 RRID:AB_2870208 |
| BV711 anti-mouse CD8a (clone: 53-6.7) | Biolegend | Cat#100759 RRID:AB_2563510 |
| Pacific Blue anti-mouse CD8a (clone: 53-6.7) | Biolegend | Cat#100725 RRID:AB_493425 |
| PE anti-mouse/human CD11b (clone:M1/70) | Biolegend | Cat#101208 RRID:AB_312791 |
| APC anti-mouse/human CD11b (clone:M1/70) | Biolegend | Cat#101212 RRID:AB_312795 |
| BUV661 anti-mouse CD11b (clone:ICRF44) | BD Biosciences | Cat#612977 RRID:AB_2870249 |
| APC anti-mouse CD11c (clone:N418) | Biolegend | Cat#117310 RRID:AB_313779 |
| Pacific Blue anti-mouse CD11c (clone:N418) | Biolegend | Cat#117322 RRID:AB_755988 |
| FITC anti-mouse CD11c (clone:N418) | Biolegend | Cat#117305 RRID:AB_313774 |
| BV570 anti-mouse CD45 (clone:30-F11) | Biolegend | Cat#103136 RRID:AB_2562612 |
| PE/Cyanine7 anti-mouse CD47 (clone: miap301) | Biolegend | Cat#127523 RRID:AB_2629544 |
| BUV395 anti-mouse CD86 (clone:GL-1) | BD Biosciences | Cat#564199 RRID:AB_2738664 |
| APC/Cyanine7 anti-mouse CD86 (clone:GL-1) | Biolegend | Cat#105030 RRID:AB_2244452 |
| PerCP anti-mouse CD86 (clone: GL-1) | Biolegend | Cat#105026 RRID:AB_893417 |
| PE/Cyanine7 anti-mouse CD172a (clone:P84) | Biolegend | Cat#144008 RRID:AB_2563546 |
| PE anti-mouse CD200 (clone:OX-90) | Biolegend | Cat#123808 RRID:AB_2073942 |
| APC anti-mouse CD206 (clone:C068C2) | Biolegend | Cat#141708 RRID:AB_10900231 |
| BV785 anti-mouse CD206 (clone:C068C2) | Biolegend | Cat#141729 RRID:AB_2565823 |
| PerCP/Cyanine5.5 anti-mouse CSF1R (clone:AFS98) | Biolegend | Cat#135525 RRID:AB_2566461 |
| BV605 anti-mouse CSF1R (clone:AFS98) | Biolegend | Cat#135517 RRID:AB_2562760 |
| FITC anti-mouse F4/80 (clone:QA17A29) | Biolegend | Cat#157310 RRID:AB_2876535 |
| Pacific blue anti-mouse F4/80 (clone:BM8) | Biolegend | Cat#123124 RRID:AB_893475 |
| BUV737 anti-mouse F4/80 (clone:T45-2342) | BD Biosciences | Cat#749283 RRID:AB_2873658 |
| APC IFNγ anti-mouse (clone:XMG1.2) | Biolegend | Cat#505810 RRID:AB_315404 |
| PE anti-mouse IL2 (clone: JES6-5H4) | Biolegend | Cat#503808 RRID:AB_315302 |
| FITC anti-mouse MHCII (clone: M5/114.15.2) | Biolegend | Cat#107606 RRID:AB_313321 |
| BV650 anti-mouse MHCII (clone:M5/114.15.2) | Biolegend | Cat#107641 RRID:AB_2565975 |
| BUV615 anti-mouse PD1 (clone: RPM1-30) | BD Biosciences | Cat#752354 RRID:AB_2875871 |
| FITC anti-mouse PD1 (clone 29F.1A12) | Biolegend | Cat#135214 RRID:AB_10680238 |
| PE anti-mouse PD1 (clone RMP1-30) | Biolegend | Cat#109103 RRID:AB_313420 |
| APC anti-mouse PDL1 (clone 10F.9G2) | Biolegend | Cat#124312 RRID:AB_10612741 |
| Percp/Cyanine5.5 anti-mouse PDL2 (clone:TY25) | Biolegend | Cat#107218 RRID:AB_2728126 |
| Percp/Cyanine5.5 anti-mouse Siglec-H (clone:551) | Biolegend | Cat#129614 RRID:AB_10643995 |

| | | |
|---|---|---|
| FITC anti-mouse TNF (clone: MP6-XT22) | Biolegend | Cat#506304 RRID:AB_315425 |
| BV510 anti-mouse XCR1 (clone: ZET) | Biolegend | Cat#148218 RRID:AB_2565231 |
| Alexa Fluor488 anti-mouse FOXP3 (clone:MF-14) | Biolegend | Cat#126406 RRID:AB_1089113 |
| PE/Cyanine7 anti-mouse GATA3 (clone:TWAJ) | Thermo Fischer Scientific | Cat#25-9966-42 RRID:AB_2573568 |
| PE/Cyanine5 anti-mouse CD62L (clone:MEL-14) | Biolegend | Cat#104410 RRID:AB_313097 |
| PE anti-mouse TCF1/7 (clone:S33-966) | BD Biosciences | Cat#564217 RRID:AB_2687845 |
| Alexa Fluor700 anti-mouse CD107a (clone:1D4B) | Biolegend | Cat#121628 RRID:AB_2783063 |
| BV786 anti-mouse Tbet (clone:O4-46) | BD Biosciences | Cat#564141 RRID:AB_2738615 |
| BV650 anti-mouse KI-67 (clone:11F6) | Biolegend | Cat#151215 RRID:AB_2876504 |
| BV450 anti-mouse EOMES (clone:Dan11mag) | Thermo Fischer Scientific | Cat#48-4875-82 RRID:AB_2574062 |
| BUV737 anti-mouse CD127 (clone:SB/199) | BD Biosciences | Cat#612841 RRID:AB_2870163 |
| BUV395 anti-mouse TIM3 (clone:5D12/TIM-3) | BD Biosciences | Cat#747620 RRID:AB_2744186 |
| anti-mouse calreticulin (clone:B44) | abcam | Cat#ab2907 RRID:AB_303402 |
| Alexa Fluor 488 anti-rabbit IgG | abcam | Cat#ab150077 RRID:AB_2630356 |
| APC/Cyanine7 anti-human CD3 (clone:OKT3) | Biolegend | Cat#317342 RRID:AB_2563410 |
| Alexa Fluor594 anti-human CD3 (clone:UCHT1) | Biolegend | Cat#300446 RRID:AB_2563236 |
| PE/Cyanine7 anti-human CD4 (clone:okt/04) | Biolegend | Cat#317414 RRID:AB_571959 |
| Alexa Fluor597 anti-human CD8 (clone:RPA-T8) | Biolegend | Cat#301056 RRID:AB_2563232 |
| PE/Dazzle anti-human CD8 (clone:Sk1) | Biolegend | Cat#344744 RRID:AB_2566515 |
| VioGreen anti-human CD8 (clone:BW135/80) | Miltenyi Biotec | Cat#130-113-726 RRID:AB_2726267 |
| BV570 anti-human CD45RO (clone:UCHL1) | Biolegend | Cat#304226 RRID:AB_2563818 |
| Pacific blue anti-human IFNγ (clone:4S.B3) | Biolegend | Cat#502532 RRID:AB_2561398 |
| FITC anti-human CD45 (clone:REA747) | Miltenyi Biotec | Cat#130-110-769 RRID:AB_2658236 |
| FITC anti-human CD163 (clone:REA812 | Miltenyi Biotec | Cat#130-112-290 RRID:AB_2655475 |
| Percp anti-human CD45 (clone:2D1) | Biolegend | Cat#368506 RRID:AB_2566358 |
| APC anti-human CD14 (clone:63D3) | Biolegend | Cat#367118 RRID:AB_2566792 |
| PE/Dazzle anti-human CSF1R (clone:AFS98) | Biolegend | Cat#135528 RRID:AB_2566523 |
| PE anti-human PDL1 (clone:M1H1) | BD Biosciences | Cat#557924 RRID:AB_647198 |
| Alexa Fluor 488 isotype control (clone:MOPC-21) | BD Biosciences | Cat#557702 RRID:AB_396811 |
| Alexa Fluor 647 isotype control (clone:MOPC-21) | BD Biosciences | Cat#557783 RRID:AB_396871 |
| PE isotype control (clone:27-35) | BD Biosciences | Cat#555058 RRID:AB_395678 |
| PE/Cyanine5 isotype control (clone:P3.6.2.8.1) | Thermo Fischer Scientific | Cat#46-4714-82 AB_1834453 |
| FcX (truStain/CD16/32) anti-mouse (clone: 93) | Biolegend | Cat#101320 RRID:AB_1574975 |
| anti-mouse MLKL (clone:3H1) | Sigma Aldrich | Cat#MABC604 RRID:AB_2820284 |
| anti-mouse pMLKL (clone:EPR9515(2)) | Abcam | Cat#ab196436 RRID:AB_2687465 |
| anti-mouse RIPK1 (clone: D94C12) | Cell signaling | Cat#3493T RRID:AB_2305314 |
| anti-mouse RIPK3 | Biorad | Cat#AHP1797 RRID:AB_2178676 |
| anti-mouse HMGB1 (clone:EPR3507 | Abcam | Cat#ab79823 RRID:AB_1603373 |
| anti-HPV-E7 (clone:8C9) | Thermo Fischer Scientific | Cat# 28-0006 RRID:AB_2533057 |
| anti-human/mouse/rat caspase8 | Thermo Fischer Scientific | Cat#PA5-77888 RRID:AB_2735575 |

| | | |
|---|---|---|
| anti-Bovine/human/mouse/sheep/rat caspase9 | Thermo Fischer Scientific | Cat#PA5-16358 RRID:AB_10985523 |
| anti-mouse PARP (clone:46D11) | Cell signaling | Cat# 9532S RRID:AB_659884 |
| anti-mouse Actin (clone: AC-74) | Sigma Aldrich | Cat#A5316 RRID:AB_476743 |
| anti-mouse Actin | Abcam | Cat#ab49900 RRID:AB_867494 |
| anti-mouse Actin (clone: AC-15) | Sigma Aldrich | Cat#A5441 RRID:AB_476744 |
| anti-rabit IgG | Cell signaling | Cat#7074S RRID:AB_2099233 |
| anti-mouse IgG | Cell signaling | Cat#7076S RRID:AB_330924 |
| anti-rat IgG2a | Biolegend | Cat#400502 RRID:AB_326523 |
| anti-mouse CSF1R (clone:AFS98) | BioXCell | Cat#BP0213 RRID:AB_2687699 |
| Chemicals,peptides and recombinant proteins | | |
| Recombinant Murine TNF | Miltenyi Biotec | Cat#130-101-687 |
| Recombinant Murine IFNβ | R&D systems | Cat#8234-MB-010 |
| Recombinant Murine IL4 | Peprotech | Cat#214-14 |
| Recombinant Murine M-CSF | Peprotech | Cat#315-02 |
| Recombinant Murine GM-CSF | Peprotech | Cat#315-03 |
| Recombinant Murine IL2 | Peprotech | Cat#210-12 |
| lipopolysaccharide Escherichia coli | Invivogen | Cat#tlrl-eblps |
| imiquimod | Invivogen | Cat#tlrl-imqs |
| 5’ppp-dsRNA/lyovec | Invivogen | Cat#tlrl-3prnaclv |
| 2’3’ cGAMP | Invivogen | Cat#tlrl-nacga23 |
| Cisplatin | Sigma | Cat#P4394 |
| Doxorubicin | Merck | Cat#D1515 |
| NP-40 cell lysis buffer | Thermo Fischer Scientific | Cat#FNN0021 |
| Phenylmethylsulfonyl fluoride (PMSF) | Thermo Fischer Scientific | Cat#1083709 |
| Pierce ™ Protease Inhibitor Mini Tablets | Thermo Fischer Scientific | Cat#A32953 |
| Pierce ™ Phosphatase Inhibitor Mini Tablets | Thermo Fischer Scientific | Cat#A32957 |
| Pierce ™ ECL Western Blotting Substrate | Thermo Fischer Scientific | Cat#32106 |
| Bovine serum albumin (BSA) | Sigma | Cat#A2153-50G |
| Criterion™ XT Bis-Tris Precast Gels | Biorad | Cat#3450124 |
| Cell Staining Buffer | Biolegend | Cat#420201 |
| Western Blot Stripping Buffer | abcam | Cat#ab270550 |
| BV-6 | Selleckchem | Cat#S7597 |
| Z-Val-Ala-Asp (OMe)-FMK | Bachem | Cat#4027403 |
| Necrostatin-1s | Bioke | Cat#17802S |
| pHrodo™ iFL Red STP Ester | Thermo Fischer Scientific | Cat#P36011 |
| recombinant HPV-E6 antigen (VYDFAFRDL) | LifeTein | NA |
| recombinant HPV-E6 antigen (DKKQRFHNI) | LifeTein | NA |
| recombinant HPV-E7 antigen (RAHYNIVTF) | LifeTein | NA |
| recombinant HPV-E7 antigen (LCVQSTHVD) | LifeTein | NA |
| Cytofix/cytoperm™ fixation/permeabilization solution | BD Biosciences Biosciences | Cat#554714 |

| | | |
|---|---|---|
| True-nuclear™ transcription factor buffer set | Biolegend | Cat#424401 |
| Dynabeads® Mouse T-activator CD3/CD28 | Thermo Fischer Scientific | Cat#11456D |
| Brefeldin A Solution (1000x) | Thermo Fischer Scientific | Cat#00-4506-51 |
| Cytofix | BD Biosciences | Cat#554655 |
| SYBRgreen | Highqu | Cat#QPD0150 |
| Bromodeoxyuridine | BD Biosciences | Cat#550891 |
| MACS tissue storage solution | Miltenyi Biotec | Cat#130-100-008 |
| phorbol myristate acetate | Sigma-Aldrich | Cat#P8139 |
| Ionomycin | Sigma-Aldrich | Cat#I9657 |
| Brefeldin A | Biolegend | Cat#420601 |
| Clodronate liposomes | Liposoma | NA |
| CellTracker CM-Dil dye | Thermo Fischer Scientific | Cat#C7000 |
| anti-mouse PDL1 (clone:M1H5) | JJP Biologics, Warsaw, Poland. Louis Boon | NA |
| anti-mouse CTLA4 (clone:4F10) | JJP Biologics, Warsaw, Poland. Louis Boon | NA |
| anti-mouse PD1 (clone:RMP1-14 | JJP Biologics, Warsaw, Poland. Louis Boon | NA |
| anti-mouse CD8 (clone:YTS169) | JJP Biologics, Warsaw, Poland. Louis Boon | NA |
| Fixable Viability Dye eFluor™ 780 | Thermo Fischer Scientific | Cat#65-0865-14 |
| Zombie Aqua™ Fixable Viability Kit A | Biolegend | Cat#423102 |
| ZOMBI NIR™ fixable viability kit | Biolegend | Cat#423106 |
| Edit-R Lentiviral CAG-Blast-Cas9 Nuclease Particles | Dharmacon Horizon discovery | Cat#VCAS10129 |
| Edit-R CRISPR-Cas9 Synthetic tracrRNA | Dharmacon Horizon discovery | Cat#U-002005-50 |
| **Critical commercial assays** | | |
| Caspase-Glo kit | Promega | Cat#G8091 |
| RealTime-Glo Annexin V Apoptosis and Necrosis assay | Promega | Cat#JA1011 |
| MTS assay kit | abcam | Cat#ab197010 |
| ELITEN ATP assay system kit | Promega | Cat#FF2000 |
| Bicinchoninic Acid (BCA) Pierce Protein Assay Kit | Thermo Fischer Scientific | Cat#23227 |
| Purelink RNA Mini Kit | Thermo Fischer Scientific | Cat#12183025 |
| QuantiTect Reverse Transcription kit | Qiagen | Cat#205313 |
| Proteome Profiler Mouse Cytokines Array Kit Panel A | R&D systems | Cat#ARY006 |
| pan-T cell isolation kit II | Miltenyi Biotec | Cat#130-095-130 |
| Murine tumour dissociation kit | Miltenyi Biotec | Cat#130-096-730 |
| anti-F4/80 microbeads | Miltenyi Biotec | Cat#130-110-443 |
| Human tumour dissociation kit | Miltenyi Biotec | Cat#130-095-929 |
| anti-human CD45 microbeads | Miltenyi Biotec | Cat#130-045-801 |
| anti-mouse CD45 microbeads | Miltenyi Biotec | Cat#130-110-618 |
| CXCL10 ELISA | R&D systems | Cat#DY466 |
| IFN alfa ELISA | Invivogen | Cat#luex-mifna |
| IFN beta ELISA | Invivogen | Cat#mifnbv2 |
| **Deposited data** | | |

| | | |
|---|---|---|
| Rational selection of syngeneic preclinical tumor mode | (Mosely et al., 2017) | GEO:GSE85509 |
| Analysis of DC vaccines used for phase II clinical trial in | (Castiello et al., 2017) | GEO:GSE85698 |
| High Dimensional Analysis Delineates Myeloid and Lyn | (Gubin et al., 2018) | GEO:GSE119352 |
| Checkpoint blockade immunotherapy induces dynami | (Kurtulus et al., 2019) | GEO:GSE122969 |
| Defining T cell states associated with response to chec | (Sade-Feldman et al., 2018) | GEO:GSE120575 |
| Single cell RNA-seq analysis of head and neck cancer | (Puram et al., 2017) | GEO:GSE103322 |
| Combination immunotherapy can rescue CD8+ T cell d | (Wang et al., 2018) | GEO:GSE120909 |
| Single-cell analysis reveals new evolutionary complexity in uveal melanoma | (Durante et al., 2020) | GEO:GSE139829 |
| Dysfunctional CD8+ T cells form a proliferative, dynam | (Li et al., 2019) | GEO:GSE123139 |
| Immune landscape of viral- and carcinogen-drived hea | (Cillo et al., 2020) | GEO:GSE139324 |
| Ensemble learning for classifying single-cell data and projection across reference atlases | (Wang et al., 2020) | GEO:GSE141982 |
| Tumor cell biodiversity drives microenvironmental rep | (Ma et al., 2019) | GEO:GSE125449 |
| Identification of grade and origin specific cell populatio | (Shih et al., 2018) | GEO:GSE118828 |
| Single cell RNA sequencing of lung adenocarcinoma | (Kim et al., 2020) | GEO:GSE131907 |
| Peripheral clonal expansion of T lymphocytes associat | (Wu et al., 2020) | GEO:GSE139555 |
| Mutagenesis sensitizes murine models of triple negative breast cancer to immunotherapy | (Hollern et al., 2019) | GEO:GSE136206 |
| Integrating microarray-based spatial transcriptomics a | (Moncada et al., 2020) | GEO:GSE111672 |
| single cell RNA-seq analysis of adult and paediatric IDH | (Neftel et al., 2019) | GEO:GSE131928 |
| Single-Cell Analyses Inform Mechanisms of Myeloid-Ta | (Zhang et al., 2020) | GEO:GSE146771 |
| Single-cell RNA-seq of melanoma ecosystems reveals s | (Jerby-Arnon et al., 2018) | GEO:GSE115978 |
| T cell landscape of non-small cell lung cancer revealed | (Guo et al., 2018) | GEO:GSE99254 |
| Single-cell RNA-seq of six thousand purified CD3+ T cells from human primary TNBCs | (Savas et al., 2018) | GEO:GSE110686 |
| Clonal replacement of tumor-specific T cells following | (Yost et al., 2019) | GEO:GSE123813 |
| **Experimental models:Cell lines** | | |
| TC1 | (Yang et al., 2016) | NA |
| TC1 *Mlkl -/-* | (Yang et al., 2016) | NA |
| TC1 Ripk3 -/- | (Yang et al., 2016) | NA |
| TC1 *Caspase8 -/-* | This paper | NA |
| TC1 *Caspase9 -/-* | This paper | NA |
| TC1 *Caspase8/9 -/-* | This paper | NA |
| MC38 | Kerafast | Cat#ENH204-FP |
| **Experimental models: Organisms/strains** | | |
| Mouse:C57BL/6J | the Jackson Laboratory | JAX: 000664 |

| | | |
|---|---|---|
| Mouse:B6.129S2-*Ifnar1* $^{tm1Agt}$ /Mmjax | the Jackson Laboratory | JAX:010830 |
| Mouse:B6.129P2*(C)-Ccr7* $^{tm1Rfor}$ */J* | the Jackson Laboratory | JAX:006621 |
| **Oligonucleotides** | | |
| murine CXCL10_forward | IDT | CGATGACGGGCCAGTGAGAA |
| murine CXCL10_reverse | IDT | CAGCCACTTGAGCGAGGACT |
| murine CXCL9_forward | IDT | AAACAGTTTGCCCCAAGCCC |
| murine CXCL9_reverse | IDT | CGAGTCCGGATCTAGGCAGG |
| murine RSAD2_forward | IDT | CGACAGCTTCGATGAGCAGG |
| murine RSAD2_reverse | IDT | ACACCTCTTTGTGACGCTCCA |
| murine MX1_forward | IDT | GGGGTCTTGACCAAGCCTGA |
| murine MX1_reverse | IDT | ACCGGCTGTCTCCCTCTGATA |
| murine IRF_forward | IDT | TGCTGAGCGAAGAGAGCGAA |
| murine IRF_reverse | IDT | CCTGCCATGCTGCATAGGGT |
| murine ACTIN_forward | IDT | CATTGCTGACAGGATGCAGAAGG |
| murine ACTIN_reverse | IDT | TGCTGGAAGGTGGACAGTGAGG |
| **Recombinant DNA** | | |
| crRNA Caspase 8 | Dharmacon™, Horizon Discovery | CCAGATTTCTCCCTACAGGT |
| crRNA Caspase 9 | Dharmacon™, Horizon Discovery | CTGTCCCATAGACAGCACCC |
| **Software and algorithms** | | |
| GraphPad Prism 8 | GraphPad Software | https://www.graphpad.com/ |
| FlowJo | BD Biosciences | https://flowjo.com |
| Image Lab 6.1 | Biorad, Image Lab | https://www.bio-rad.com/en-be/product/image-lab-software?ID=KRE6P5E8Z#fragment-6 |
| Biorender | Biorender | https://biorender.com/ |
| Phyton (version 3.9.7) | Phyton | https://www.python.org/ |
| Gistic | Broad institute | https://portals.broadinstitute.org/tcga/gistic/browseGisticByGene |
| Morpheus | Broad institute | https://software.broadinstitute.org/morpheus/ |
| Stream | Human Cell atlas data portal | https://data.humancellatlas.org/analyze/methods/stream |
